# Three-dimensional Virtual Adult Cardiomyocyte Transcriptomics

**DOI:** 10.64898/2026.04.14.718375

**Authors:** Cheng Luo, Yuyan Lyu, Xinning Guo, Li Cheng, Qiantong Liang, Songmao Wang, Yanzhi Wang, Shikuan Zhang, Shengnan Wang, Tianyu Liu, Yongjiang Luo, Fang Lu, Boli Ran, Yaou Zhang, Xiao Liu, Yi Wang, Gangjian Qin, Jiangbin Wu, Qing Rex Lyu

**Affiliations:** Department of Cardiology, Chongqing Academy of Medical Science, Chongqing General Hospital, Chongqing University, Chongqing 401147, China; Tsinghua Shenzhen International Graduate School, Shenzhen 518055, China; Department of Cardiology, Renji Hospital, School of Medicine, Shanghai Jiao Tong University, Shanghai 200127, China; Medical Research Center, Chongqing Academy of Medical Science, Chongqing General Hospital, Chongqing University, Chongqing 401147, China; Department of Pharmacology, Cardiovascular Research Institute, School of Medicine, Southern University of Science and Technology, Shenzhen 518055, China; Department of Cardiovascular Surgery of First Affiliated Hospital and Institute for Cardiovascular Science, Suzhou Medical College, Soochow University, Suzhou 215000, China; State Key Laboratory of Silkworm Genome Biology, Biological Science Research Center, Southwest University, Chongqing 400715, China; School of Medicine, Chongqing University, Chongqing 400030, China

## Abstract

Whole-cell transcriptomic profiling of adult cardiomyocytes at single-cell resolution remains challenging because of their large size, elongated morphology and frequent multinucleation ^1–4^. Spatial transcriptomics (ST) preserves tissue architecture while measuring gene expression *in situ*. However, many existing analytical frameworks rely on nuclear segmentation, limiting their ability to delineate adult cardiomyocytes accurately ^5–11^. In this study, we introduce 3D-VirtualCM (three-dimensional virtual cardiomyocyte), a membrane-guided framework for reconstructing volumetric cardiomyocyte transcriptomes. By integrating cell-membrane contour similarity with cross-sectional optimal transport, 3D-VirtualCM provides a panoramic transcriptomic view of the mouse heart. Applying 3D-VirtualCM to adult mouse hearts after myocardial infarction, we identified spatially and transcriptionally distinct cardiomyocyte populations within the infarct border zone. It also enabled high-throughput quantification of cardiomyocytes exhibiting cell-cycle re-entry signatures and characterization of their associated molecular programs. Moreover, 3D-VirtualCM revealed heterogeneous RNA distributions along the longitudinal axis of individual cardiomyocytes. By integrating three-dimensional cellular morphology with *in situ* transcriptomes, 3D-VirtualCM provides a scalable approach for resolving cardiomyocyte states and spatial organization during physiological and pathological cardiac remodeling.

---

Adult cardiomyocytes are highly specialized cells that form the contractile myocardium and generate the force required to sustain blood circulation ^12,13^. Their large size, elongated rod-shaped morphology and frequent multinucleation hinder efficient isolation and capture, limiting whole-cell transcriptomic profiling using current single-cell RNA-sequencing platforms ^6,14–17^. As a result, the transcriptomic landscape of adult cardiomyocytes, particularly their cytoplasmic RNA profiles, remains incompletely characterized across physiological and pathological states ^18,19^.

Spatial transcriptomics (ST) profiles gene expression in tissue sections while preserving spatial context ^20,21^. However, adult cardiomyocytes are elongated cells that typically span 100-150 μm in length ^22^. A single ST section therefore often captures only a fraction of each cardiomyocyte, limiting direct reconstruction of volumetric transcriptomes at single-cell resolution. Several methods align and integrate serial ST sections, but they have primarily targeted tissue-scale reconstruction in brain and tumor tissues ^23–28^. Their reliance on nucleus-centered segmentation also makes them poorly suited to specialized adult cardiomyocyte morphology. Thus, reconstructing spatially resolved volumetric transcriptomes of adult cardiomyocytes remains technically challenging.

Here, we introduce 3D-VirtualCM, a membrane-guided framework for reconstructing adult cardiomyocyte morphology and spatial transcriptomes in three dimensions. We used Cellpose to segment wheat germ agglutinin (WGA)-stained cell membranes across 10 consecutive ST sections from an infarcted mouse heart ^29^. For volumetric reconstruction, we developed HiDTW, a hierarchical contour-tracking algorithm that matches cardiomyocyte boundaries across serial sections and integrates their transcriptomes into three-dimensional single-cell representations. Benchmarking and simulation analyses showed that HiDTW outperformed existing methods and baselines on WGA-stained cardiomyocyte contour datasets. Compared with single-section ST and single-nucleus RNA sequencing (snRNA-seq), 3D-VirtualCM recovered more UMIs and genes per cell and resolved post-infarction cardiomyocyte heterogeneity at greater granularity. The framework also supported high-throughput quantification of cardiomyocytes exhibiting cell-cycle re-entry signatures and characterization of their associated transcriptional programs. Furthermore, 3D-VirtualCM revealed heterogeneous RNA distributions along the longitudinal axis of individual cardiomyocytes, which we validated using orthogonal experiments. Together, 3D-VirtualCM reconstructs the volumetric architecture of adult cardiomyocytes and links three-dimensional cell morphology with spatial gene expression across thick tissues. It therefore provides a framework for investigating adult cardiomyocytes and other structurally complex cell types across physiological and pathological states.

## Membrane-based cell segmentation of adult cardiomyocytes

Owing to their elongated morphology and frequent multinucleation, single-section ST captures only a fraction of each adult cardiomyocyte and its transcriptome (**Fig. 1a**). Current ST analysis frameworks rely predominantly on nucleus-centered segmentation, limiting accurate delineation of cardiomyocyte boundaries (**Extended Data Fig. S1a**) ^30,31^. To address these limitations, we developed a membrane-guided framework that combines cell-boundary recognition with cross-sectional contour tracking to reconstruct volumetric transcriptomes of adult cardiomyocytes at single-cell resolution (**Fig. 1b**). We first used a human-in-the-loop (HITL) workflow to generate ground-truth membrane annotations from WGA-stained cardiac sections (**Extended Data Fig. S1b**) ^29^. These annotations and the corresponding raw images from serial sections were then used to train an adult cardiomyocyte membrane recognition (ACMR) model based on *Cellpose* ^29^. Across four independent validation datasets, ACMR achieved a mean contour overlap of 84.38% with HITL-defined cardiomyocyte boundaries (**Extended Data Fig. S1c-d**).

**Fig. 1.**
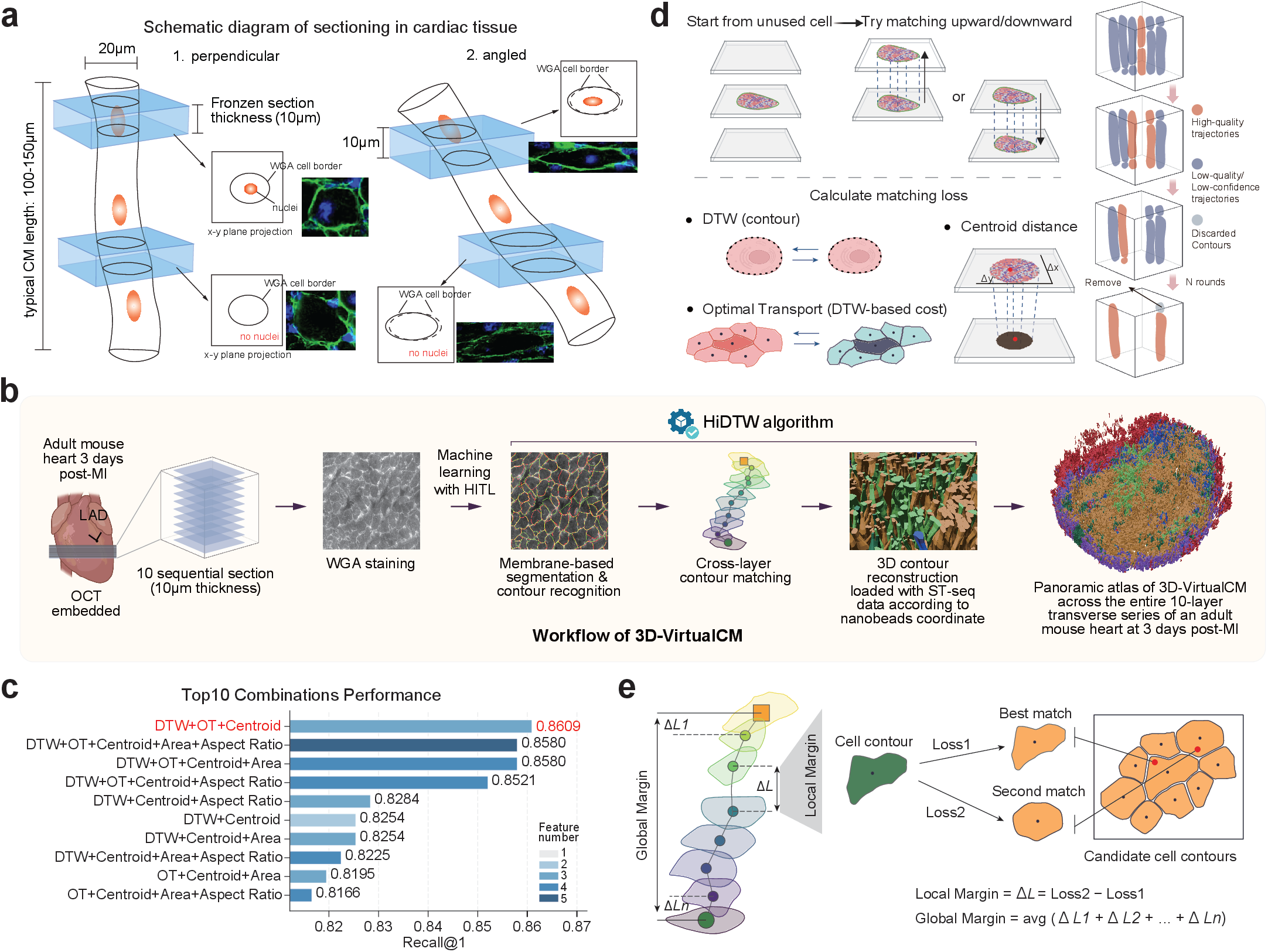
Three-dimensional reconstruction of adult cardiomyocytes. **a.** Schematic diagram of single-layer sectioning in cardiac tissue, illustrating cardiomyocytes with longitudinal axes oriented either perpendicular or oblique to the sectioning plane; **b.** Workflow of 3D-VirtualCM; **c.** Top 10 feature combinations ranked by Recall@1 across all tested metric combinations. Recall@1 denotes the proportion of cases in which the ground-truth match was ranked first; **d.** Schematic illustration of the HiDTW algorithm integrating DTW, centroid displacement and optimal transport-based neighborhood matching, together with the trajectory evaluation process; **e.** Schematic illustration of the margin-based z-stitching strategy across serial layers. Two margin metrics were used: local margin for pairwise ranking and global margin for trajectory-level ranking.

## Three-dimensional reconstruction of adult cardiomyocyte contours

To track individual adult cardiomyocytes across serial sections and reconstruct their three-dimensional contours, we generated a reference dataset using sparse, stochastic cardiomyocyte-specific GFP labeling in adult mouse hearts (**see Methods**). GFP-labeled cardiomyocytes extended across multiple sections, and their continuous signals provided empirical ground-truth for contour matching and optimization of membrane-based three-dimensional reconstruction **(Extended Data Fig. S2a, Supplemental Data S1)**. Analysis of this dataset showed that correctly matched contours retained consistent geometric and morphological features across adjacent sections **(Extended Data Fig. S2b)**. We then assessed five candidate matching metrics using positive and negative contour pairs. Most metrics reliably distinguished true matches from negative pairs **(Extended Data Fig. S2c).**

To test whether nonlinear feature interactions improved matching accuracy, we compared logistic regression and random forest models across multiple metric combinations (**Extended Data Fig. S2d-f**). The two models performed comparably; therefore, we selected logistic regression in downstream analysis for its greater interpretability. Next, we performed exhaustive feature-combination screening and ablation analyses, and identified dynamic time wrapping (DTW), optimal transport (OT), and centroid displacement as the best-performing feature set for contour matching **(Fig. 1c, Extended Data Fig. S2g-h)**. Incorporating OT substantially improved pairing performance in cases that remained ambiguous when using DTW and centroid displacement alone **(Extended Data Fig. S3a-b)**. Based on these results, we developed HiDTW, a contour-tracking algorithm that combines the selected geometric features with local neighborhood information to trace individual cardiomyocytes across serial sections for three-dimensional reconstruction **(Fig. 1d)**.

During z-stitching, pairing accuracy and coverage exhibited a clear trade-off **(Extended Data Fig. S4a)**. We therefore introduced a margin criterion (Top1−Top2) to balance matching confidence against reconstruction completeness. For each contour, candidate matches were ranked by a composite matching loss (*L*), and matching ambiguity was quantified from the relative difference between the best and second-best candidates. Consistent with this rationale, low-margin pairs were preferentially enriched for incorrect matches **(Extended Data Fig. S4b; see Methods)**. We incorporated the margin threshold as an explicit z-stitching hyperparameter and jointly optimized it with the DTW, OT, and centroid displacement weights using Bayesian optimization. To improve trajectory consistency across serial sections, we implemented a dual-normalization framework combining local and global confidence measures **(Fig. 1e)**. Specifically, the local margin (Δ*L*) quantified contour-level confidence at each z-step, whereas the global margin was computed on a task-calibrated scale and aggregated across sections to assess overall trajectory quality.

After generating a pool of candidate trajectories in each round, contour-level conflicts were resolved by retaining trajectories with higher trajectory scores. Unassigned contours were passed to subsequent rounds, and the procedure was iterated until no additional trajectories could be generated **(Fig. 1d, see Methods)**. We then examined margin distributions and trajectory acceptance rates across serial sections and optimization rounds. Both local and global margin distributions, as well as trajectory acceptance rates, remained stable throughout reconstruction **(Extended Data Fig. S4c-f)**. Statistical validation further showed that the HiDTW reconstruction-quality metrics were significantly higher than those produced by two complementary null models **(Extended Data Fig. S4g-h)**. Parameter-sensitivity analysis identified that the local-margin threshold and candidate-retention fraction primarily governed the trade-offs among coverage, trajectory length, and matching confidence (**Extended Data Fig. S4i-n**). Correspondingly, higher local margins were associated with smaller inter-section centroid displacement and greater leave-one-out (LOO) stability (**Extended Data Fig. S4o-p**). Together, these analyses support the robustness of HiDTW for membrane-referenced three-dimensional reconstruction of adult cardiomyocytes.

## Benchmarking HiDTW using ground-truth and simulated datasets

We independently applied Bayesian optimization to each of three ground-truth datasets (**Extended Data Fig. S5a**). The optimization used a global score combining average trajectory length, coverage ratio, and global margin as the surrogate objective for reconstruction quality. Comparisons with null distributions generated by ranking permutations and morphology shuffling showed that the global score distinguished empirical constructions from both null models (**Extended Data Fig. S5b**). For each dataset, we selected the parameter combination yielding the highest global score and used it to summarize HiDTW reconstruction performance. Across the three datasets, HiDTW achieved pair-level accuracies of 84.11%, 83.64%, and 81.72%; coverage rates of 94.09%, 93.69%, and 90.83%; and track-level accuracies of 76.92%, 77.22%, and 77.71%, respectively. Thus, the global score correlated positively with both pair- and track-level accuracy across all ground-truth datasets (**Extended Data Fig. S5c-d; Supplemental Data S2**).

Error analysis further showed that mispairings were enriched among contour pairs with low local margins **(Extended Data Fig. S6a-e)**. Decomposition of the loss components for mispaired contours identified centroid displacement as the largest contributor to matching error (**Extended Data Fig. S6f)**. Potential sources of this displacement included inter-section shifts, tissue deformation, compression and local mechanical perturbations introduced during sectioning. To test whether sectioning contributed to these errors, we performed two-photon microscopy of thick GFP-labelled cardiac tissue, eliminating serail sectioning and reducing associated geometric distortions **(Extended Data Fig. S7a-d; Supplemental Data S3)**. Applying HiDTW to these data yielded cardiomyocyte contours with 95.08% pair-matching accuracy, 90.91% track-level accuracy, and 94.41% coverage **(Extended Data Fig. S7e)**. Among 177 ground-truth pairings, only three were incorrectly predicted **(Extended Data Fig. S7f)**. These results support robust HiDTW reconstruction when sectioning-induced deformation is minimized.

We next established a phantom-based benchmark comprising 54 simulated datasets across 18 conditions (**Fig. 2a**). The benchmark systematically varied cell density, spatial clustering, inter-section translation, local positional jitter, interfering cells and contour deformation. Using a unified evaluation framework, we compared HiDTW with three baselines: centroid nearest neighbor matching (Centroid_NN), DTW-only matching (DTW_only), and OT-only matching (OT_only). Overall, HiDTW achieved higher matching accuracy and pair-level F1 scores, together with lower trajectory fragmentation, than all three baselines **(Fig. 2b-d)**. Centroid_NN showed the largest performance decline under inter-section displacement and local clustering, whereas DTW_only and OT_only were more susceptible to matching errors under contour perturbation (**Fig. 2e**). Together, these benchmarks support the accuracy and robustness of HiDTW for three-dimensional reconstruction of adult cardiomyocyte contours across the ground-truth and simulated conditions tested.

**Fig. 2.**
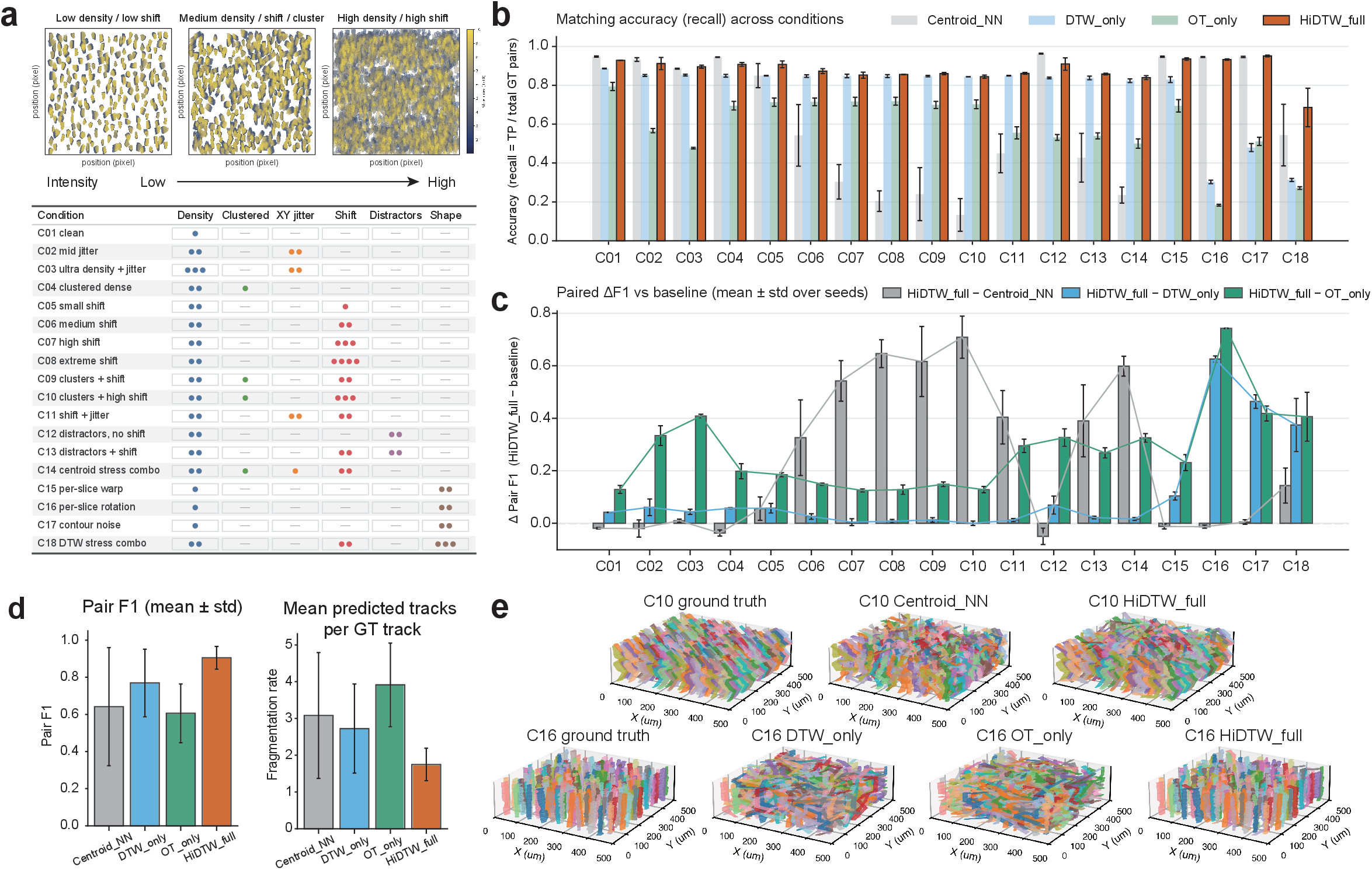
HiDTW maintains high contour-matching accuracy across diverse perturbation conditions. **a.** Benchmarking design for systematic evaluation of contour matching across adjacent sections. Representative simulated datasets illustrate increasing reconstruction difficulty. The table summarizes the 18 benchmark conditions (C01-C18), incorporating different levels of contour density, clustering, XY jitter, inter-slice shift, distractors, centroid stress, and shape perturbation. **b.** Recall of ground-truth contour pairs across all benchmark conditions for Centroid_NN, DTW_only, OT_only, and HiDTW_full. Accuracy was calculated as the proportion of true positive matched pairs among all ground-truth pairs. Bars show mean values and error bars indicate s.d. across random seeds. **c.** Paired comparison of pair F1 score between HiDTW_full and each baseline method across all benchmark conditions. Bars indicate the mean difference in pair F1 (Δ pair F1) over random seeds, with error bars denoting s.d. **d.** Aggregate performance summary across benchmark conditions. Left, mean pair F1 score for each method. Right, mean predicted tracks per ground-truth track, reflecting the degree of reconstruction fragmentation. Data are presented as mean ± s.d. across all conditions and random seeds. **e.** Representative 3D reconstruction examples from challenging simulation settings illustrating failure modes of individual baselines and the corresponding reconstructions by HiDTW_full.

## Generation of panoramic 3D transcriptional profiles of adult cardiomyocytes

To generate a three-dimensional, single-cell-resolution panoramic transcriptomic atlas of the adult mouse heart, we collected 10 consecutive 10-µm cryosections from adult mouse hearts after experimental myocardial infarction. The sections were mounted directly onto BMKMANU S3000 spatial-transcriptomics chips and subsequently stained for nuclei and WGA (**Extended Data Fig. 8a-b**). Nuclear and WGA images were registered with the barcoded nanobeads coordinates in a shared coordinate system, enabling retrivel of corresponding RNA-sequencing data (**Extended Data Fig. S8c**). Macroscopic alignment of the serial images was then performed using *Linear Stack Alignment with SIFT* (**Extended Data Fig. S9a**)^32^. The resulting image stack showed consistent registration across all 10 sections (**Extended Data Fig. S9b-d**).

We next applied HiDTW across the serial sections, yielding volumetric transcriptomic profiles of 23,308 adult cardiomyocytes (**Fig. 3a**). These reconstructed profiles contained a median of 15,470 unique molecular identifiers (UMIs) and 2,716 detected genes per cardiomyocyte (**Fig. 3b; Extended Data Fig. S10a-c; Supplemental Data S4**). For reference, snRNA-seq of an adjacent tissue recovered a median of 4,741 UMIs and 2,031 detected genes per nucleus. Single-section 10× Visium HD data processed using a nuclear-centered segmentation yielded medians of 739 UMIs and 459 detected genes per cardiomyocyte (**Fig. 3b; Supplemental Data S5**).

**Fig. 3.**
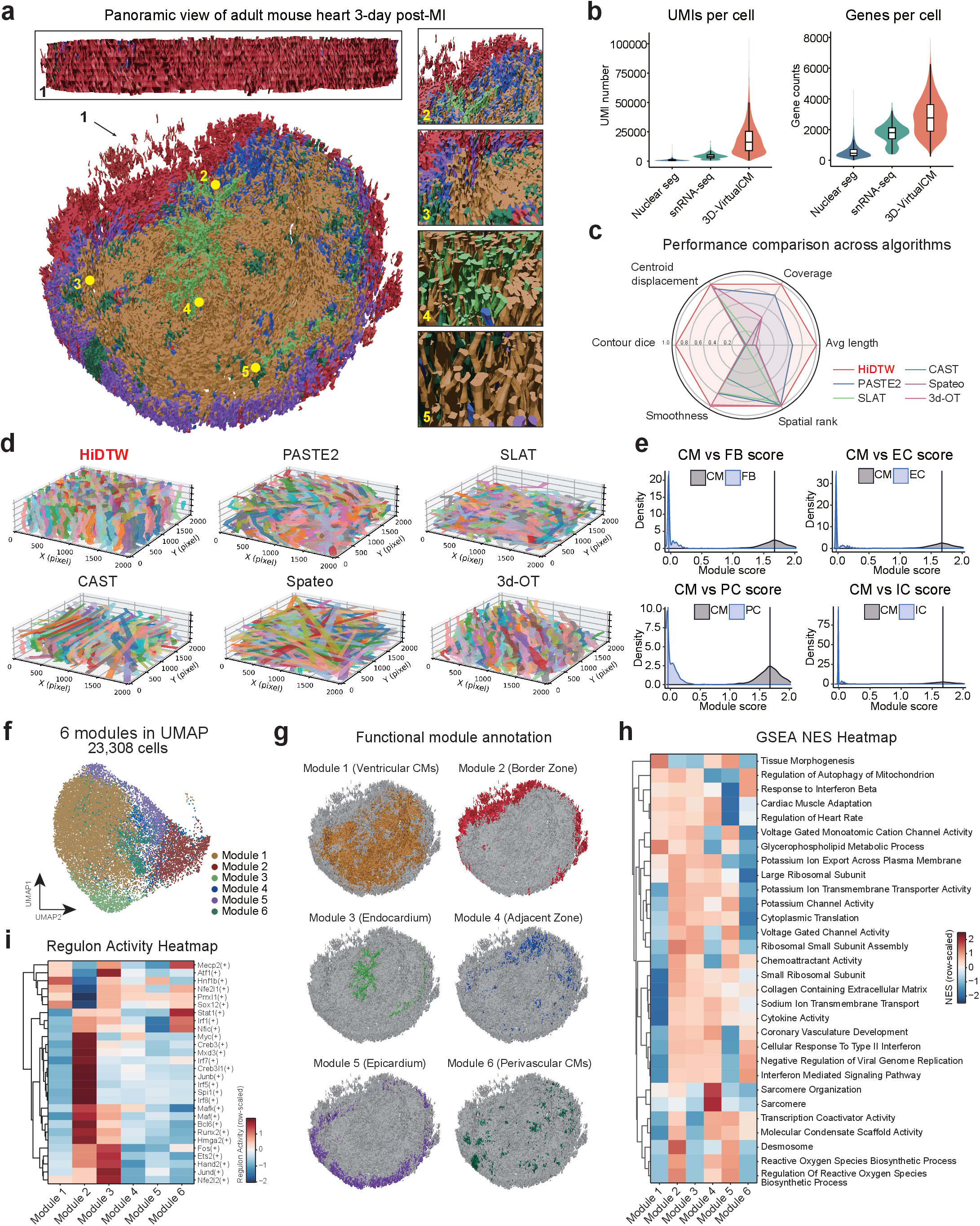
Single-cell panoramic mapping and benchmarking of the mouse heart 3 days after myocardial infarction. **a.** Panoramic 3D reconstruction of a 100µm-thick cardiac tissue section from an adult mouse at day 3 post-myocardial infarction. Each cell is integrated with *in situ* RNA transcriptomic and spatial information; **b.** Comparison of UMI and gene counts per cardiomyocyte across different platforms; **c.** Radar plot summarizing direction-normalized benchmark performance metrics for HiDTW, PASTE2, SLAT, CAST, Spateo and 3d-OT; **d.** Representative reconstructed cardiomyocyte trajectories generated by different algorithms; **e.** Comparison of density distributions of 3D-VirtualCM module scores between cardiomyocytes (CM) and non-CM cell types (FB, fibroblasts; EC, endothelial cells; PC, pericytes; IC, immune cells); **f.** UMAP of adult cardiomyocytes reconstructed by 3D-VirtualCM and annotated by functional modules; **g.** Spatial projection of functional modules in the adult mouse heart, with module annotations mapped back to their original spatial coordinates; **h.** Heatmap of GESA NES across different functional modules; **i.** Assessment of transcription factor activity across different functional modules.

To evaluate reconstruction accuracy, we benchmarked HiDTW against representative methods for cross-sectional alignment within a unified evaluation framework. Because cardiomyocyte nuclei were not consistently captured across adjacent sections, nucleus-position-based reconstruction methods could not be evaluated directly. We therefore compared HiDTW with five representative gene-expression-based methods: PASTE2^23^, SLAT^27^, CAST^26^, Spateo^33^, and 3d-OT^24^. HiDTW achieved the strongest overall performance across the evaluation metrics (**Fig. 3c-d, Extended Data Fig. S11a-g**). To isolate the contribution of contour-based trajectory matching, we also constructed three morphology-based baselines: local centroid nearest-neighbor matching (Baseline 1), global Hungarian assignment (Baseline 2), and neighborhood-structure matching (Baseline 3) (**Extended Data Fig. S11h, see Supplemental Methods**). HiDTW outperformed all three baselines under every tested condition, demonstrating the benefit of integrating geometric similarity with local tissue context for cross-sectional contour reconstruction (**Extended Data Fig. S11i-k**).

To assess the transcriptomic specificity of 3D-VirtualCM, we examined potential non-cardiomyocyte contamination using cell-type-specific signatures derived from our snRNA-seq dataset and two independent published references (GSE76092 and GSE293209) ^34,35^. Reconstructed cardiomyocytes showed only low and sporadic expression of non-cardiomyocyte marker genes (**Fig. 3e; Extended Data Fig. S12a-i**), supporting that 3D-VirtualCM faithfully preserved cardiomyocyte-specific transcriptional identities and limited detectable cross-cell-type contamination.

3D-VirtualCM resolved 14 transcriptional clusters among adult cardiomyocytes (**Extended Data Fig. S13a**). Based on their spatial distribution, we grouped these clusters into six modules annotated as ventricular, border-zone (BZ), endocardial, adjacent-zone (AZ), epicardial, and perivascular cardiomyocytes (**Fig. 3f-g; Extended Data Fig. S13b**). We next identified representative genes enriched in each module. Module 1 was marked by *Myl3*, *Fabp3* and *Ckmt2* expression and was distributed throughout the left and right ventricles. Module 2 showed high expression of *Nppb*, *Postn* and *Spp1* and was enriched in the BZ. Module 3 was marked by *Atp2a2*, *Myom2* and *Npr3* and was localized preferentially to the endocardial region. Module 4 expressed *Myh7*, *Trim63* and *Ttn* and was enriched in the AZ surrounding the infarct. Module 5 was marked by *Kcnd2*, *Fxyd1* and *Tnnt2* and localized to the epicardial region. Finally, Module 6 expressed *Mylk*, *Tagln* and *Tcap* and localized to perivascular regions (**Extended Data Fig. S13c; Supplemental Data S6**).

Previous microdissection-based studies reported substantial cardiomyocyte heterogeneity within the infarct BZ ^35,36^. Consistent with these observations, 3D-VirtualCM resolved two transcriptionally distinct cardiomyocyte populations adjacent to the ischemic zone (IZ), corresponding to the BZ (Module 2) and AZ (Module 4) (**Extended Data Fig. S14a-d; Supplemental Data S7**). Functional enrichment analysis revealed marked differences between these modules. Module 2 was enriched for genes involved in protein synthesis and reactive oxygen species production, whereas Module 4 preferentially expressed genes associated with muscle development and sarcomere organization (**Fig. 3h; Extended Data Fig. S14e; Supplemental Data S8**). Regulon analysis further identified distinct regulatory programs between the two modules. Specifically, *Junb*- and *Myc*-associated regulons showed higher activity in Module 2 than in Module 4, whereas *Sox12*- and *Atf1*-associated regulons showed lower activity (**Fig. 3i**). Together, these analyses show that 3D-VirtualCM integrates volumetric reconstruction with high-depth spatial transcriptomics to resolve peri-infarct cardiomyocyte heterogeneity at single-cell resolution not readily achieved using conventional approaches.

## Identifying cardiomyocytes in the cell cycle

Cardiomyocytes that re-enter the cell cycle are of interest because they may contribute to cardiac repair after injury ^37^. However, their rarity and the lack of definitive cardiomyocyte-specific cell-cycle markers hinder their identification *in situ* ^18^. Consequently, existing studies often rely on manual quantification and provide limited information about their broader transcriptomic states ^38^. By integrating volumetric reconstruction with ST, 3D-VirtualCM supports high-throughput detection, quantification and molecular characterization of cardiomyocytes in the cell-cycle (CM-CC)-associated transcriptional signatures.

Using established cell-cycle markers, including *Mki67*, *Aurkb*, and *Top2a*, we identified 619 *Mki67*^+^, 234 *Aurkb*^+,^ and 530 *Top2a*^+^ cardiomyocytes. Of all reconstructed cardiomyocytes, 44 co-expressed *Mki67*^+^ and *Aurkb*^+^ (0.19%), 68 co-expressed *Mki67*^+^ and *Top2a*^+^ (0.29%), and 13 co-expressed all three markers (0.056%) (**Fig. 4a; Extended Data Fig. S15a-f**). As an orthogonal validation, we performed RNA FISH and quantified transcript puncta in mouse hearts 3 days after myocardial infarction. RNA FISH detected *Mki67*^+^/*Aurkb*^+^ and *Mki67*^+^/*Top2a*^+^ cardiomyocytes at frequencies of 0.31% and 0.68%, respectively (**Extended Data Fig. S16a-c**). In agreement with previous reports^38^, the CM-CC population was predominantly localized to the BZ (**Fig. 4b**).

**Fig. 4.**
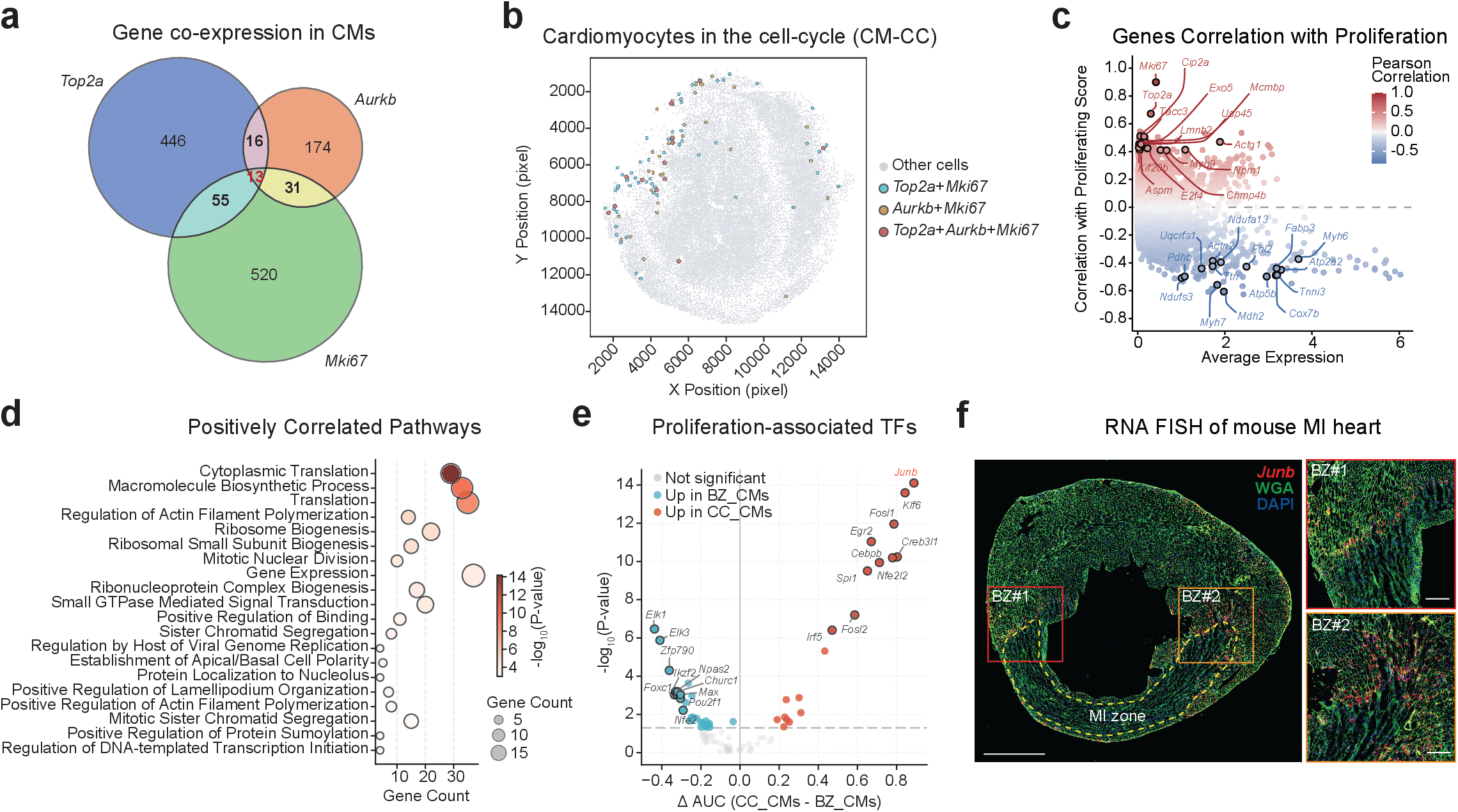
Identification of cardiomyocyte in the cell cycle using 3D-VirtualCM. **a.** The Venn diagram shows the expression and co-expression of *Mki67*, *Aurkb*, and *Top2a* in 3D-VirtualCM-derived cardiomyocytes; **b.** The mapping of *Mki67*^+^/*Top2a*^+^, *Mki67*^+^/*Aurkb*^+^, *Mki67*^+^/*Aurkb*^+^/*Top2a*^+^ cardiomyocytes onto an adult mouse heart projection; **c.** The Pearson correlation assessment of genes associated with proliferation marker genes *Mki67*, *Aurkb*, and *Top2a*; **d.** The Gene Ontology functional annotation of cardiomyocytes is positively associated with a proliferation marker in (**c**); **e.** The volcano plot of proliferation-associated transcription factors between cardiomyocytes in the cell-cycle (CM-CC) and other cardiomyocytes in the Border Zone (CM-BZ); **f.** Co-staining of WGA and *Junb* RNA FISH in a transverse mouse heart section.

To characterize the molecular features associated with cell-cycle activity, we defined a CM-CC score based on the combined expression of *Mki67*, *Aurkb* and *Top2a*. We then calculated Pearson correlations between the expression of each gene and the CM-CC score across cardiomyocytes in the cell-cycle (**Fig. 4c**). Gene Ontology biological process (GO-BP) analysis showed that positively correlated genes were enriched in cytoplasmic translation, actin filament polymerization, DNA replication and cell-cycle phase transition processes (**Fig. 4d; Supplemental Data S9**). To characterize regulatory programs associated with CM-CC, we inferred transcription factor (TF) activity and compared CM-CC with other BZ cardiomyocytes (CM-BZ) ^39^. *Junb* showed the largest increase in inferred TF activity in CM-CC (**Fig. 4e**). As an independent validation, RNA FISH detected higher *Junb* transcript abundance in neighboring CM-BZ cardiomyocytes (**Fig. 4f**).

## 3D-VirtualCM reveals asymmetric RNA distribution in adult cardiomyocytes

Localized translation in cardiomyocytes raises the possiblity that intracellular RNA distribution is spatially heterogeneous ^40,41^. By reconstructing adult cardiomyocytes through approximately 100-μm tissue, 3D-VirtualCM enables systematic analysis of intracellular RNA localization at single-cell resolution. For these analyses, the cardiomyocyte end oriented toward the atrium was defined as the proximal, and the opposite end was defined as distal (**Extended Data Fig. S17a**). We focused on ventricular cardiomyocytes reconstructed across 10 consecutive sections and compared transcriptomes from the three sections nearest each end. Dimensionality reduction revealed a gradual and asymmetric transcriptomic transition along the longitudinal axis of individual cardiomyocytes (**Fig. 5a**). Sequencing depth and the number of detected genes remained stable across sections, arguing against systematic technical variation (**Extended Data Fig. S17b-f**). Housekeeping genes and canonical cardiomyocyte markers likewise showed no systematic section-dependent bias (**Extended Data Fig. S17g**).

**Fig. 5.**
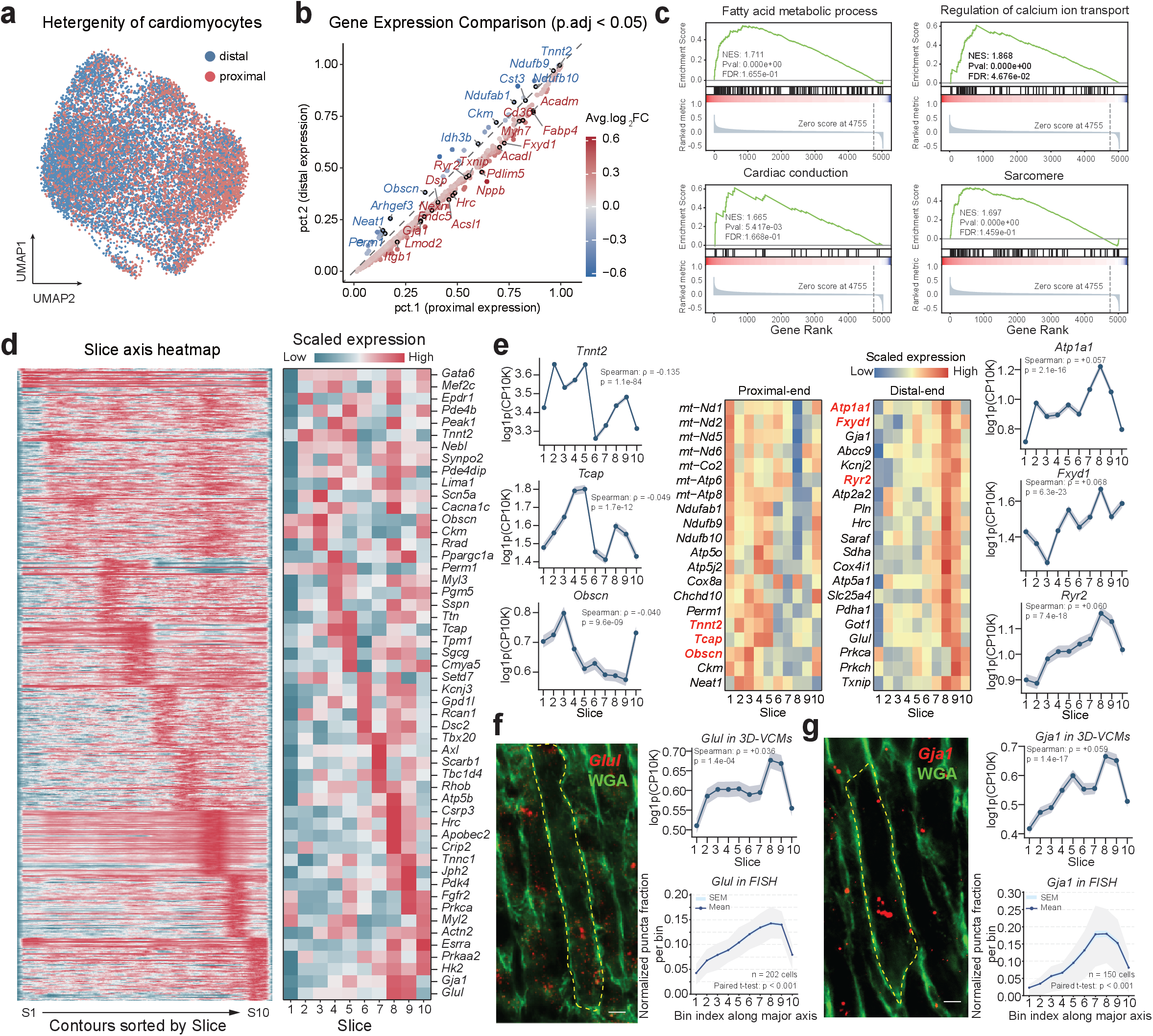
3D-VirtualCM reveals heterogeneity of adult mouse cardiomyocytes. **a.** UMAP of the distal and proximal ends of cardiomyocytes reveals the heterogeneity of adult cardiomyocytes; **b.** Scatter plot displays differentially expressed genes (DEG) (adj.p<0.05) between the distal and proximal regions; the x-axis represents the proportion of expression in the proximal end, and the y-axis represents the proportion in the distal end; **c.** The representative gene ontology biological processes (GO-BP) of the upregulated functional pattern in the proximal end. NES, normalized enrichment score; FDR, false discovery rate; **d.** Heatmap showing gene expression dynamics along the slice axis in reconstructed cardiomyocytes. Left, heatmap of slice-dependent expression patterns; right, selected representative genes of heatmap; **e.** Identification of genes with significant longitudinal expression gradients across slices. Middle, heatmaps of representative genes enriched toward the proximal end or distal end. Left and right, representative examples of genes showing significant positive or negative correlations with slice position after sequencing-depth-corrected regression analysis. Expression is shown as log1p(CP10K), CP10K indicates counts per 10,000. Spearman’s correlation coefficients and corresponding p values are indicated. **f-g.** Validation of asymmetric subcellular RNA distribution by 3D RNA fluorescence *in situ* hybridization (FISH). Representative images and quantification of *Glul* **(f)** and *Gja1* **(g)** transcripts in adult cardiomyocytes. Cardiomyocyte boundaries are outlined by WGA staining. Right, upper panels show transcript abundance across slices in 3D-VirtualCM (3D-VCMs); lower panels show normalized puncta fraction per bin along the major axis measured by RNA FISH. The shaded band indicates the mean ± SEM across cells within each slice, with SEM reflecting the precision of the per-slice mean.

We next identified genes preferentially enriched at either end, revealing distinct intracellular expression programs (**Fig. 5b; Supplemental Data S10**). Proximal, intermediate and distal regions also exhibited distinct transcriptional signatures. Proximal-enriched genes were associated with oxidative phosphorylation, mitochondrial electron transport and translational elongation. By contrast, distal-enriched genes were associated with synapse assembly, muscle structure development and regulation of energy metabolism (**Fig. 5c-d; Extended Data Fig. S17h**). Together, these analyses revealed longitudinal polarization of RNA distribution within adult cardiomyocytes, with distinct biological processes represented at opposite ends.

To systematically identify genes with polarized intracellular localization, we fitted sequencing-depth-corrected linear regression models using the z-axis slice index of each reconstructed contour as the independent variable. This analysis identified genes with significant positive or negative expression gradients along the longitudinal axis (**Fig. 5e; Supplemental Data S11**). Recent studies have reported neuron-like properties of cardiomyocytes, including endogenous glutamatergic and cholinergic signaling systems ^42,43^. In line with these reports, three-dimensional mapping showed that metabolic genes were preferentially localized proximally, whereas genes associated with glutamatergic and cholinergic signalings were enriched distally (**Fig. 5e**). These asymmetric patterns remained evident in cardiomyocytes reconstructed from 7 to 10 consecutive sections, indicating robustness to variation in reconstruction length (**Extended Data Fig. S18a-b**). As an independent validation, three-dimensional RNA FISH (z-stacking) showed asymmetric intracellular distribution of *Glul* and *Gja1* transcripts within adult cardiomyocytes (**Fig. 5f-g**). Together, these results revealed longitudinal RNA polarization within adult cardiomyocytes and showed that 3D-VirtualCM can resolve intracellular RNA organization *in situ*.

## Discussion

In this study, we developed 3D-VirtualCM, a membrane-guided framework that reconstructs volumetric representations of adult cardiomyocytes across serial tissue sections while retaining matched *in situ* transcriptomic information. Adult cardiomyocytes are particularly difficult to profile using existing methodology because their large size, elongated morphology and frequent multinucleation make nucleus-centered segmentation an unreliable proxy for cell boundaries. By using WGA-stained membrane contours as the primary structural reference and tracking them across the z-axis, 3D-VirtualCM links cellular morphology, tissue position and molecular state within the same reconstructed cardiomyocytes. Benchmarking using empirical ground-truth and simulated datasets further indicates that this strategy can recover cross-sectional cell trajectories despite variation in local tissue geometry.

3D-VirtualCM builds on advances in tissue reconstruction, spatial transcriptomics and volumetric imaging. We noticed that CODA, PASTE and 3d-OT support three-dimensional reconstruction, and CaMVIA-3D enables direct volumetric imaging of cardiomyocytes in thick tissue sections ^24,28,44,45^. However, these approaches operate primarily at the level of tissue architecture, spatial spots or cellular morphology. None explicitly tracks membrane-defined adult cardiomyocytes across serial spatial transcriptomic sections to recover matched volumetric transcriptomes for individual cells. The distinction of 3D-VirtualCM lies in preserving cardiomyocyte identity across serial sections and coupling spatially distributed transcripts to membrane-defined volumetric cells (**Supplementary Table S1**). 3D-VirtualCM enables, for the first time, integrated analysis of morphology, spatial position and molecular state within the same reconstructed cardiomyocytes.

By recovering transcripts distributed across multiple cross-sections, 3D-VirtualCM increased transcript detection relative to single-section ST and snRNA-seq, improving the resolution of peri-infarct cardiomyocyte heterogeneity. It also enabled scalable identification and molecular characterization of cardiomyocytes expressing cell-cycle-associated transcripts, although marker expression alone does not establish completed mitosis, cytokinesis or new cardiomyocyte formation. Beyond intercellular heterogeneity, 3D-VirtualCM revealed longitudinal RNA asymmetry within individual cardiomyocytes, including polarized *Gja1* and *Glul* distributions, suggesting structured intracellular RNA organization without establishing discrete compartments or functional consequences. Nevertheless, RNA FISH validation was restricted to these two prespecified transcripts in approximately 150-200 cardiomyocytes per transcript, limiting transcriptome-wide generalization. Higher-density axial sampling, multiplexed RNA imaging and functional perturbation will be required to establish the prevalence and biological significance of this polarization.

Several technical constraints define the current scope of 3D-VirtualCM. Reconstruction accuracy depends on membrane-staining quality, section continuity and cross-sectional alignment. Tissue loss, compression, local deformation or poorly preserved membranes may reduce reconstruction confidence and coverage. The current workflow is best suited to large cells with clearly delineated membrane boundaries and will require cell-type-specific training and validation for smaller or densely packed cells. In addition, the 10-μm axial sampling interval and the lateral resolution of the spatial transcriptomic platform limit the spatial scale at which RNA localization can be resolved. The detected gradients should therefore be interpreted as coarse longitudinal patterns rather than fine-scale molecular localization.

In summary, 3D-VirtualCM extends serial-section reconstruction from tissue-level alignment to cell-resolved molecular measurement. By integrating membrane-referenced morphology, native spatial context and matched transcriptomic information, it provides a new experimental layer for examining how adult cardiomyocyte architecture and molecular signatures vary during cardiac diseases. This framework provides a foundation for investigating morphologically complex cells in intact tissues while preserving explicit links among cellular shape, spatial position and molecular state.

## Supporting information

All suppl files

## Acknowledgements

This study was supported by grants from the National Natural Science Foundation of China (82270908, 82470464, 82470396, 82570558), the Shanghai Municipal Commission of Science and Technology (24ZR1445100), and startup funding from Chongqing General Hospital to Q.R.L. and Soochow University to J.B.W.

AI-assisted tools, including Grammarly and ChatGPT, were used to correct grammatical errors and improve the clarity of the English text and assist coding. The authors have reviewed the manuscript in full and take responsibility for all content presented in this study.

## Author Contributions

Drs. Q.R.L., Y.Y.L., J.B.W., and C.L. conceived and designed the study. Dr. C.L. performed the bioinformatics analyses, developed algorithm scripts, analyzed the data, and generated the figures. Dr. Y.Y.L. conducted animal experiments, sectioned and stained tissues, and assisted with data analysis and figure preparation. Drs. Y.Z.W., and S.K.Z. performed RNA *in situ* hybridization. Drs. X.N.G., S.M.W., S.N.W., and T.Y.L. contributed to human-in-loop cell segmentation. Dr. L.C. prepared samples for two-photon microscopy. Dr. Q.T.L. provided consultation on mathematical algorithms and formulas. Drs. B.L.R., Y.O.Z., and X.L. provided consulting support. Drs. Y.J.L., F.L., and Y.W. trained the cell segmentation model and assisted with data processing. Drs. Q.R.L., C.L., and Y.Y.L. wrote the draft. Drs. Q.R.L., J.B.W., G.J.Q., and Y.W. supervised the study and revised the manuscript.

## Materials and Methods

### Animal models

The animal experiments were conducted in accordance with the National Institutes of Health Guidelines on the Care and Use of Laboratory Animals (NIH Publication, 8th edition, 2011) and were approved by the Institutional Animal Care and Use Committee of Yishang Biotech. (Shanghai, China). All mice were maintained under controlled conditions: temperature of 24 ± 2°C, humidity of 40-60%, and a 12-hour light/dark cycle, with *ad libitum* access to food and water.

#### Myocardial infarction model

Coronary artery ligation was performed to establish the MI model. Briefly, 8-week-old male C57BL/6J mice were anesthetized and positioned on a surgical platform. An incision was made in the fourth intercostal space of the left chest to expose the heart, and the left anterior descending (LAD) coronary artery was ligated 2mm distal to its origin with a 6-0 silk suture after the heart was smoothly and gently externalized. Afterwards, the heart was placed back into the intrathoracic space, followed by manual evacuation of air and closure of the muscle and the skin. Three days post-LAD ligation surgery, the heart was harvested and embedded in optimal cutting temperature (OCT) compound (Cat# 4583; Tissue-TEK, USA).

#### GFP labeling of mouse cardiomyocytes in vivo

To establish a sparsely GFP-expressed pattern in the adult mouse heart, AAV9-cTnT-EGFP and AAV9-cTnT-Cre-EGFP were produced and administered subcutaneously to postnatal day 1 (P1) C57BL/6J mice with 1×10^12^ vg virion per mouse. The adult mouse heart was harvested at P26 and embedded in OCT compound.

### Tissue sectioning

The OCT-embedded hearts were cryosectioned at varying depths (10μm, 20μm and 100μm) according to experimental requirements, using a Leica cryostat (CM1950; Leica, USA).

### RNA *in situ* hybridization

RNA *in situ* hybridization (RNA-FISH) was performed on frozen tissue sections (10μm for slide scanner and 20μm z-stack imaging) using RNAscope Multiplex Fluorescent Reagent Kit v2 (#323100, ACD, USA) following the official RNAscope protocols. Target-specific probes, including *Mki67* (#416771, ACD, USA), *Aurkb* (#461761, ACD, USA), *Top2a* (#491221, ACD, USA), *Junb* (#556651, ACD, USA), *Gja1* (#486191, ACD, USA), *Glul* (#426231, ACD, USA), together with ACD positive and negative control probes, were used. Signals were developed using the RNAscope Multiplex Fluorescent (or chromogenic) detection workflow. WGA (#32466, Thermo Fisher, USA) was used to visualize the cell border of cardiomyocytes. Sections were counterstained, mounted, and imaged using identical acquisition settings for quantitative analysis. We used a Pannoramic MIDI scanner (3DHISTECH, Hungary) for low-power imaging and a laser confocal microscope (#FV3000, Olympus, Japan) for high-power imaging.

### GFP-labeled cardiomyocyte imaging

For establishing a ground truth, 10μm-thick cryosections were fixed in 4% PFA, and anti-GFP antibody (#CL488-66002, Proteintech, China) was incubated to enhance signal for imaging using a slide scanner. 20× panoramic view of the whole tissue sections was imaged by Panoramic MIDI scanner (3DHISTECH, Hungary).

For two-photon imaging, 100μm thick cryosections were fixed in 4% PFA and washed in PBS. Overnight WGA (#32466, Thermo Fisher, USA) incubation was performed at 4°C. After temperature equilibration and washing steps, sections were imaged directly on a Nikon AX R MP two-photon microscope (Nikon, Japan) without a coverslip using a 25× water-immersion objective lens (CFI75 Apo 25XC W 1300) (#MRD77225, Nikon, Japan).

### Code and data availability

All code used in this study is publicly available on GitHub: https://github.com/qlyu1/3D-Virtual-Cardiomyocyte-Transcriptomics. The related datasets associated with the algorithm are available on Figshare: https://doi.org/10.6084/m9.figshare.31305214.

### Spatial transcriptomics RNA sequencing

#### 1. BMKMANU S3000 platform

Sequencing libraries were prepared according to the manufacturer’s instructions using the BMKMANU Library Construction Kit (#ST03002-34, Biomarker, China). The libraries were constructed using Truseq Illumina library preparation kits. Sequencing was performed on a NovaSeq platform (Illumina) by Integragen (Evry). The sequencing was performed according to the recommended protocol.

For each sample, FASTQ files and manually aligned histology images were analyzed using BSTMatrix version 2.4.f.2. The resulting data were then mapped to the genome using the STAR genome aligner (version 2.5.1b). The processed data were subsequently imported into R using Seurat (version 4.0.1). Genes with expression counts in fewer than 5 spatial spots were discarded, and spots with evidence of folding were also removed.

#### 2. 10× Genomics Visium HD platform

Freshly collected mouse heart tissues were submitted to Majorbio Bio-pharm Technology Co., Ltd. (Shanghai) and processed on the 10× Genomics Visium HD platform following the manufacturer’s instructions. OCT blocks were cryosectioned at 10μm and placed within the fiducial frame of Visium Spatial slides. H&E staining and decrosslinking were performed following the Visium Spatial Gene Expression workflow for CytAssist and imaging optimization. Brightfield images were acquired with a 20× objective on a Pannoramic MIDI scanner (3DHISTECH, Hungary), stitched in Pannoramic MIDI software. Spatial transcriptomics was performed using 10x Genomics Visium HD Slide (#1000670, Millipore Sigma, USA) with mouse transcriptome probe hybridization (#1000667, Millipore Sigma, USA); ligated probe products were released and captured by spatially barcoded oligonucleotides, and libraries were constructed using Dual Index Plate TS Set A (i7/i5) according to the Visium HD Spatial Gene Expression Reagent Kits User Guide (#CG0006855, 10× Genomics, USA). Library quality and quantity were assessed using an Agilent 5300 TapeStation (Agilent, USA) and a Qubit Fluorometer (Thermo Fisher, USA).

Raw FASTQ files and corresponding histology images were processed per sample using Space Ranger v1.3.1 (10× Genomics, USA). Reads were aligned with STAR to the 10× Genomics Cell Ranger GRCm39 reference. Per-spot QC metrics were assessed in R using Seurat v4.0, and spots with fewer than 200 detected UMIs were excluded.

### Single-nucleus RNA-seq

Samples were homogenized using a glass Dounce tissue grinder (#D8938, Sigma, USA) and lysed in ice-cold Nuclei EZ lysis buffer. Nuclei were pelleted by centrifugation (500g, 5 min, 4°C). The nuclei pellet was then washed and passed through a 35μm cell strainer (#352235, Corning-Falcon, USA). Single-nucleus RNA-seq libraries were generated using the Chromium Next GEM Single Cell 3′ Reagent Kits v3.1 on a Chromium Controller (10× Genomics, USA). Nuclei from cultured cell lines were resuspended and loaded onto a Chromium Next GEM Chip G, and partitioned into GEMs following the manufacturer’s protocol. Within GEMs, nuclei were lysed and RNA was barcoded during reverse transcription, followed by cDNA amplification and library construction per the kit instructions. Libraries were sequenced on an Illumina NovaSeq 6000 (PE150). Reads were aligned with 10x Cell Ranger v7.0 (STAR) to the mouse mm10 reference genome, and UMI counts were computed per barcode-gene. Cell Ranger’s barcode-calling was used to identify nuclei-containing barcodes for downstream analyses. Biomarker Technologies (Beijing, China) performed the assay.

### Algorithms and Bioinformatic Analysis

#### 1. Cell contour preprocessing and feature extraction

To reconstruct 3D trajectories from 2D sections, we convert each cell contour into a fixed-length, translation-invariant point-sequence representation, compute per-cell geometric features, and build an intra-slice adjacency graph used for neighborhood-based OT computation.

**Contour ordering and resampling.** For each cell *j*, we represent its original contour as a point set 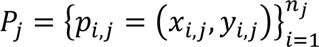, where *n_j_* denotes the number of contour vertices for cell *j* and may vary across cells. Then points are sorted clockwise in ascending order of the centroid-referenced polar angle defined as:

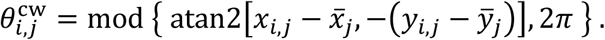

The starting point is chosen as the contour vertex with the smallest *y*-coordinate (corresponding to the 12 o’clock direction in image coordinates), which ensures consistent parameterization across cells. The reordered contour sequence is denoted by (*q*_1,*j*_, …, *q_nj_*_,*j*_). We close the contour by setting *q_nj_*_+1,*j*_ = *q*_1,*j*_ and index the ordered contour by cumulative arc length:

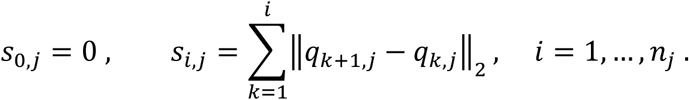

To facilitate downstream contour-based computation, we resample each cell contour to a fixed number *N* = 200 of boundary points used throughout this work, so that all cells are represented in a fixed-length point sequence. Specifically, we resample *N* uniformly spaced positions on [0, *s_nj_*_,*j*_] via linear interpolation of *x* and *y*; if *s_nj_*_,*j*_ = 0 (a degenerate zero-perimeter contour), we replicate the single point *N* times. For translation invariance, we center the sampled contour by subtracting its centroid and get 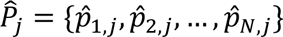.

**Geometric features.** From the original contour, we compute per-cell features including centroid, polygon area (shoelace formula; absolute value), and an ellipse-based aspect ratio obtained by least-squares fitting (OpenCV fitEllipse). If ellipse fitting fails or the minor axis is zero, we set the aspect ratio to 1.

**Intra-slice adjacency graph.** To construct the intra-slice adjacency graph, we compute the minimum pairwise Euclidean distance between the resampled contour points of cells *i* and *j* in the original image coordinates, denoted by *d_ij_*. Cells *i* and *j* are defined as neighbors if *d_ij_* < 5 pixels; we store neighbor IDs and the neighbor count for each cell.

#### 2. Dynamic Time Warping (DTW)

To quantitatively measure contour similarity between adjacent slices, we adopt a Dynamic Time Warping (DTW)–based distance and use the CUDA-accelerated Soft-DTW implementation for large-scale batch computation. For a cell *c_i_^z^* on slice *z* and a candidate cell *c_i_^z+1^* on slice *z* + 1, we extract their 2D boundary contours, each uniformly resampled to *N* points (Section 1) and get 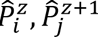. The DTW distance between the two contours is defined as:

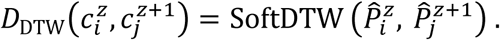

#### 3. Optimal Transport (OT)

To complement the DTW contour distance with local context, we introduce a neighborhood distance based on optimal transport (OT), which measures the overall matching quality between the two surrounding neighborhood blocks. For a cell *c_i_^z^* on slice *z* and a candidate cell *c_i_^z+1^* on slice *z* + 1, their neighborhood blocks are denoted by *C_i_^z^*, *C_i_^z+1^* respectively, each consisting of the cell and all of its intra-slice neighbors defined in Section 1.

##### 3.1 Cost matrix via Soft-DTW

All contours are preprocessed and uniformly resampled to *N* points as described in Section 1. Denote the preprocessed contour point sequences of *a_r_* µ *C_i_^z^* and *b_s_* µ *C_i_^z+1^* by *P*^(*a_r_*) and *P*^(*b_s_*), respectively. Then the pairwise shape cost is defined as:

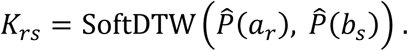

and the cost matrix is ***K*** = (*K_rs_*). Thus, K collects the pairwise Soft-DTW costs for all cell pairs between the two neighborhood blocks.

##### 3.2 Area-weighted entropic OT

To represent the relative contribution of each segmented object within its neighborhood block, we use its 2D projected area to define the OT mass distribution. This area-based weighting reduces the influence of small truncated regions, fragmented detections, and other small non-target structures during neighborhood matching. Let *A*(*a_r_*) and *A*(*b_s_*) be the polygon areas (Section 1) of *a_r_* and *b_s_*. The source and target masses are:

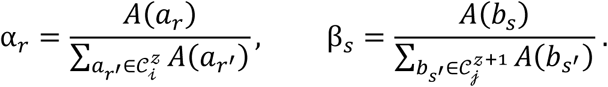

If both neighborhood blocks contain only the cell itself, with no adjacent neighbors, we set the neighborhood OT cost to zero by convention, since no additional neighborhood context is available.

Given ***K***, ***α***, ***β*,** we compute an entropy-regularized OT plan *T* using a GPU-accelerated Sinkhorn algorithm:

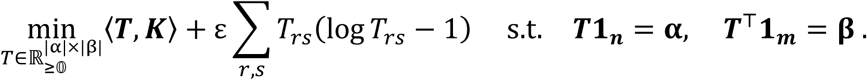

with adaptive regularization strength:

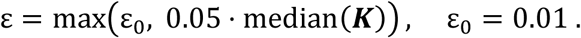

The lower bound ε_0_ = 0.01 prevents numerical instability when the median cost is close to zero. After calculating the entropy-regularized coupling, we evaluate only the primal transport-cost term at the regularized solution:

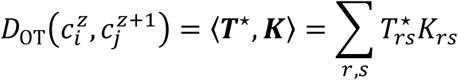

This quantity is used as a transport-based neighborhood-matching score rather than as a classical, parameter-independent OT distance. The entropy term is introduced primarily to improve numerical stability and smooth the transport solution.

#### 4. Dual-Scale Normalization for Stitching

During stitching, correspondences are estimated by a joint cost combining contour (DTW), neighborhood (OT), and centroid terms. To ensure comparability across tissues/regions/slice sets, we use dual-scale normalization for DTW and OT: a step-wise local scale for candidate selection and a region-wise global scale for quality reporting. Locally, we set the scale as the 75th percentile of valid raw distances with outlier clipping, compute OT only for top-ranked candidates (others receive a constant penalty); globally, we fix a normalization factor per region by randomly sampling cell pairs and taking the 75th percentile of their raw DTW/OT distances, and use it for trajectory-level assessment and cross-region/parameter comparisons.

#### 5. Margin-based match quality estimation

After computing the contour (DTW), neighborhood (OT), and centroid distances, we make matching decisions using a local margin criterion and use the margin statistics to define trajectory-level and global quality scores.

##### 5.1 Candidate-set scoring with local/global normalization

For each current cell, we first identify candidate cells in the next slice within a spatial search window. For every candidate, we compute three complementary distances to the current cell: a contour-based distance, a neighborhood-based distance, and a centroid distance. These three terms are combined with fixed weights to form two parallel losses. The locally normalized loss is used to rank candidates within the current decision point, while the globally normalized loss uses fixed reference statistics estimated in advance and is used for quality reporting across different decision points.

##### 5.2 Local margin acceptance rule

We convert candidate losses into scores using a monotone decreasing transform, so that lower loss corresponds to higher score. Candidates are ranked by the local score. To determine whether the best candidate is sufficiently better than the second-best one, we define a local margin (Local Margin):

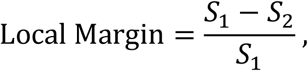

where *S*_1_ and *S*_2_ are the top-1 and top-2 local scores, respectively. A linkage is accepted only if this local margin exceeds a preset threshold α otherwise, it is rejected as unreliable for trajectory extension. For accepted linkages, we also record the corresponding globally normalized margin for later quality aggregation.

##### 5.3 Trajectory-level margin quality and conflict resolution

For each trajectory, we summarize linkage reliability by averaging the globally normalized local margins across all accepted linkages. To support conflict resolution, we combine this margin-based quality measure with a normalized trajectory-length term. The length term acts as a prior that reduces the tendency of very short trajectories to dominate the conflict-resolution process. The final trajectory score is computed by combining the margin quality and normalized trajectory length. After each round of candidate trajectory generation, we rank all candidate trajectories according to this final score and retain the top ρ fraction for selection, with ρ = 40%. Contours assigned to the retained trajectories are removed from subsequent matching, whereas unused contours are passed to the next round to generate additional candidate trajectories.

##### 5.4 Global margin statistics and overall scoring

To characterize overall matching ambiguity, we record a margin statistic at every decision point. If candidates are available, we record the local margin and whether it passes the acceptance threshold; otherwise, we assign a margin value of zero and mark the position as having no candidate. The average of these per-point margins is then rescaled to a confidence score. We further compute a coverage score and a length score, and define the final global score as a weighted combination of confidence, coverage, and length.

#### 6. Bayesian-optimization-based parameter search framework

We wrap the outer layer of our margin-based matching module into a Bayesian optimization (BO) loop. BO automatically tunes the local margin threshold and distance-mode weights by maximizing a region-wise reconstruction-quality score. We implement the optimizer with gp_minimize (scikit-optimize) and perform multi-run evaluations for each candidate setting.

##### 6.1 Parameter space and constraints

Each single-step matching decision is controlled by the hyperparameter vector ***θ*** = (α, *w*_DTW_, *w*_OT_, *w*_cen_), where α is the local margin acceptance threshold and (*w*_DTW_, *w*_OT_, *w*_cen_) are the weights for contour (Soft-DTW), neighborhood (OT), and centroid distances. We search a 4D continuous space:

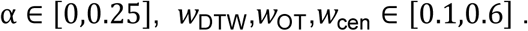

##### 6.2 Outer-loop objective

Given θ, we run the full 3D cell-trajectory reconstruction pipeline in the target region and compute three global metrics: Conf_glob_, CovScore and LenScore. We combine them into a scalar objective:

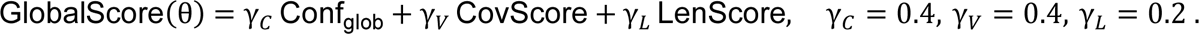

The length term serves as a mild regularizer to discourage trivially short trajectories.

##### 6.3 GP surrogate and sampling strategy

We adopt GP-based BO using gp_minimize with a zero-mean GP regressor and a Matérn kernel. The acquisition function is gp_hedge, which adaptively selects among LCB, EI, and PI; acquisition maximization uses the default random sampling plus L-BFGS. BO starts with *n*_init_ uniformly sampled initial points (8 in this study). For each evaluated *θ_i_*, we minimize the negative score:

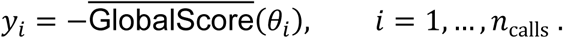

The GP is fitted to {(*θ_i_*, *y_i_*)} to obtain posterior mean *μ*(*θ*) and variance *σ*^2^(*θ*). Subsequent evaluations choose *θ_t_*_+1_ by maximizing the acquisition function under the current posterior until the evaluation budget is exhausted. After optimization, we select the best configuration from all evaluated candidates:

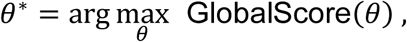

and use *θ*^∗^ as the final parameter setting for that region.

#### 7. Simulation data and controlled benchmarking design

To evaluate inter-slice cell matching under known ground truth, we generated parameterized three-dimensional cardiomyocyte phantoms within a fixed 500 × 500 × 100 *μ*m volume and virtually sectioned them into ten consecutive two-dimensional slices of 10μm thickness. Each simulated cell was represented as a capsule-like three-dimensional phantom whose cross-sectional contour was randomly sampled from a library of real cardiomyocyte contours, with configurable geometric properties, and each ground-truth trajectory comprised the observations of the same cell across adjacent sections.

The simulation included multiple controlled conditions with different spatial densities and perturbations, including Poisson-like or clustered cell placement, slice-wise rigid translation, per-cell lateral jitter across sections, and slice-specific distractor contours without cross-slice identity. We compared three single-cue baselines based on centroid proximity, contour similarity, or neighborhood optimal transport with the full HiDTW pipeline used in practice, with HiDTW parameters optimized by 20 rounds of Bayesian optimization.

#### 8. Benchmarking against alternative methods

To compare HiDTW with published methods that could be adapted for cross-slice alignment, we benchmarked it against five representative nucleus-independent approaches based on spatial coordinates and gene expression, including PASTE2, SLAT, CAST, Spateo, and 3d-OT. For the alternative methods, inter-slice correspondences were inferred from spatial and transcriptomic information; for HiDTW, the same cells were represented by contour geometry. To ensure comparability, all resulting adjacent-slice correspondences were normalized to one-to-one matches and linked in slice order using the same downstream trajectory-construction and conflict-resolution procedure. All alternative methods were run using the scripts provided in their respective GitHub repositories with default parameter settings.

Performance was assessed using label-free metrics computed from adjacent links along predicted trajectories, including centroid displacement, contour Dice coefficient, trajectory smoothness, trajectory length, cell coverage, and spatial rank. Here, spatial rank refers to the rank of the matched target cell among all candidate cells in the corresponding target slice when candidates are ordered by centroid distance to the source cell from nearest to farthest. These metrics were designed to quantify geometric consistency, continuity, and biological plausibility of the reconstructed cross-slice trajectories under a unified evaluation framework.

#### 9. 3D-VirtualCM expression data filling

To estimate gene expression at the single-cell level from spatial transcriptomics data, we mapped spatial barcodes to corresponding cell contours using segmentation masks derived from tissue images. For each cell contour, we summed per-gene counts across all mapped barcodes to generate a contour-by-gene expression matrix. Image registration, barcode assignment, and aggregation were implemented in custom scripts. (see code repository).

#### 10. 3D-VirtualCM data preprocessing

Each section was split into a 10×10 grid and processed/stiched per subregion to improve efficiency and preserve local structure. Raw count matrices were imported into R (v4.1.3) and converted to Seurat objects (Seurat v4.3.0). QC metrics included nFeature_RNA, nCount_RNA, and percent.mt. Subregions were merged and processed with: LogNormalize (scale factor 10,000), 2,000 HVGs (“vst”), scaling, PCA (20 PCs), SNN graph on PCs (FindNeighbors, dims = 1:10), clustering (FindClusters, resolution = 1.0), and UMAP for visualization. Downstream analyses used this processed dataset.

#### 11. Differential expression and module-enriched genes

Differential expressions were performed in Seurat using the Wilcoxon rank-sum test with Benjamini-Hochberg (BH) FDR correction; genes with BH-adjusted p (FDR)<0.05 were considered significant. Module-enriched genes were identified with FindAllMarkers (adjusted p<0.05). To account for abundance, we computed each marker’s UMI fraction using the raw count matrix and compared gene-wise UMI percentages within the target module versus all remaining cells.

#### 12. GSEA and GO enrichment

Preranked GSEA was run with gseapy (v1.1.8). For each module, genes were ranked by Seurat avg_log_2_FC and tested against MSigDB mouse gene sets (release 2024.1.Mm; m5.go collection). We used 1,000 permutations and controlled multiple testing by BH FDR correction; gene sets with FDR<0.05 were considered significant. GO enrichment was additionally performed using Enrichr (https://maayanlab.cloud/Enrichr/).

#### 13. pySCENIC regulon inference and activity analysis

Regulons were inferred with pySCENIC (v0.12.1): (i) GRNBoost2 network inference using a mouse TF list (mm_mgi_tfs.txt); (ii) cisTarget motif enrichment/pruning with mm9 TSS-centered 5 kb, 7-species mc9nr rankings and v10nr motifs; and (iii) AUCell scoring to obtain per-cell regulon AUC. For CC-CMs vs BZ-CMs, we binarized regulon activity, computed group-wise mean AUC and activation rate, and tested AUC differences using two-sided Mann-Whitney U tests with BH FDR correction across regulons (statsmodels).

#### 14. 3D-VirtualCM contour-expression analysis

Contour-based expression profiles from 10-layer 3D-VCMs (Module 1) were imported into Seurat. QC metrics (nFeature_RNA, nCount_RNA, percent.mt) were computed and visualized; we retained cells with intermediate library sizes (nCount_RNA between the 15th and 75th percentiles). Slice identity was parsed from cell barcodes and stored as metadata. The filtered data were re-normalized and scaled while regressing out nCount_RNA, followed by PCA. Clustering used PCs 1-10 (FindNeighbors, dims = 1:10; FindClusters, resolution = 0.6), and UMAP was computed on PCs 1-10 (RunUMAP, dims = 1:10).

