## Supplementary material for "Three-dimensional Virtual Adult Cardiomyocyte Transcriptomics": All suppl files: 518649_3_extended_data_5189828_tpd5v8_convrt.pdf

### Extended Data Fig. S1. HITL-assisted training and validation of a cardiomyocyte membrane-boundary segmentation model.

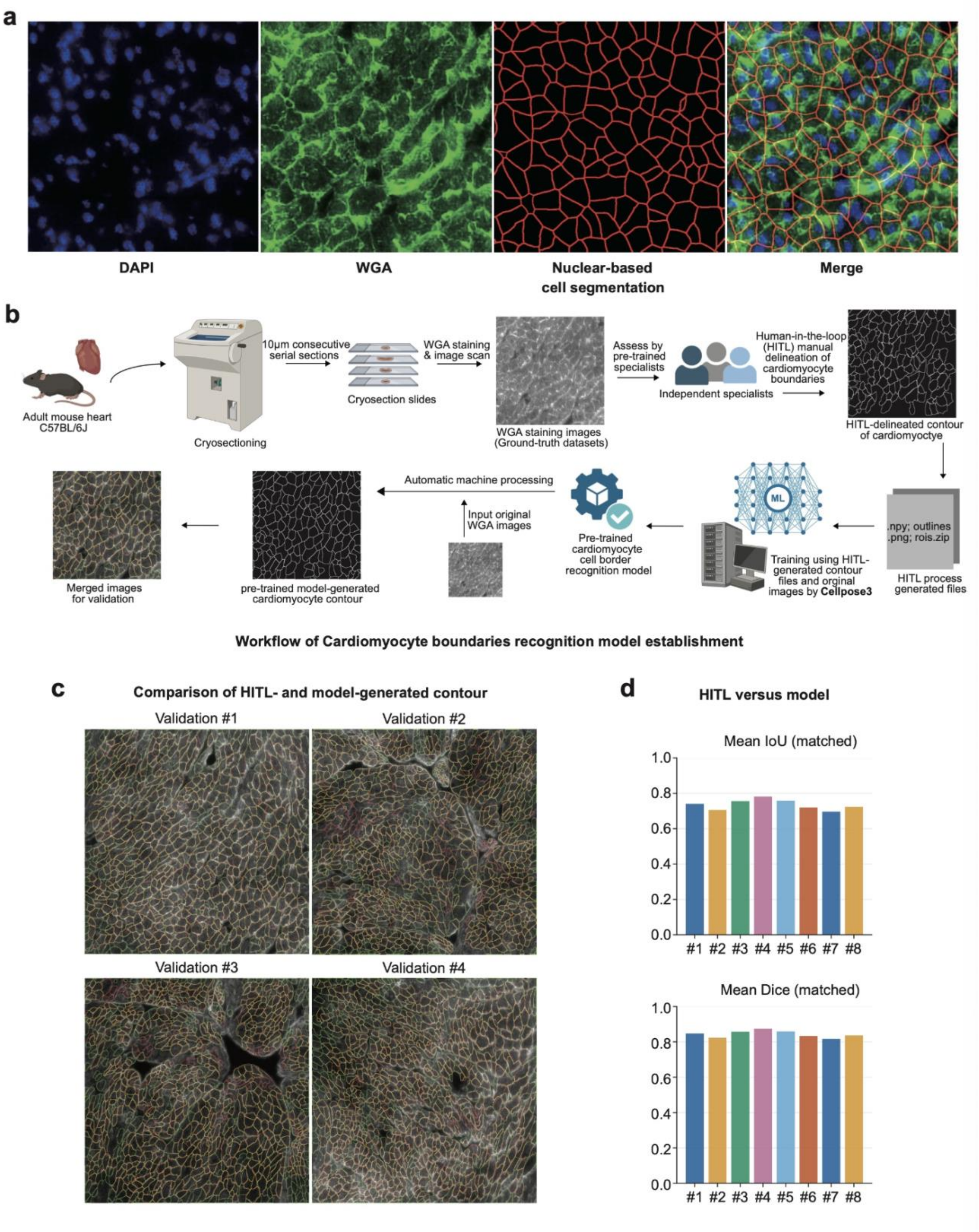

**a.** DAPI (blue) and WGA (green) staining with nuclear-based segmentation outlines (red), showing inaccurate boundary assignment in cardiac tissue. **b.** Workflow for 5 establishing an adult cardiomyocyte boundary recognition (ACMR) model. Adult mouse hearts were harvested and cryosectioned (10μm consecutive sections), WGA-stained, and imaged. Cardiomyocyte borders were manually delineated using a human-in-the-loop (HITL) strategy to generate training files, which were used to train a Cellpose-based model. The pre-trained model was then applied to WGA images to generate contours and benchmark against the validation dataset. **c.** Representative validation images showing consistency between HITL- delineated contours and model-generated contours (green line: manually delineated; red lines: Cellpose generated, yellow lines: merged). **d.** Quantitative comparison of HITL versus model segmentations across validation images using mean Intersection-over-Union (IoU) and mean Dice coefficients for matched cells.

**Extended Data Fig. S2. Ground-truth matches cardiomyocyte pairs and metric benchmarking.**

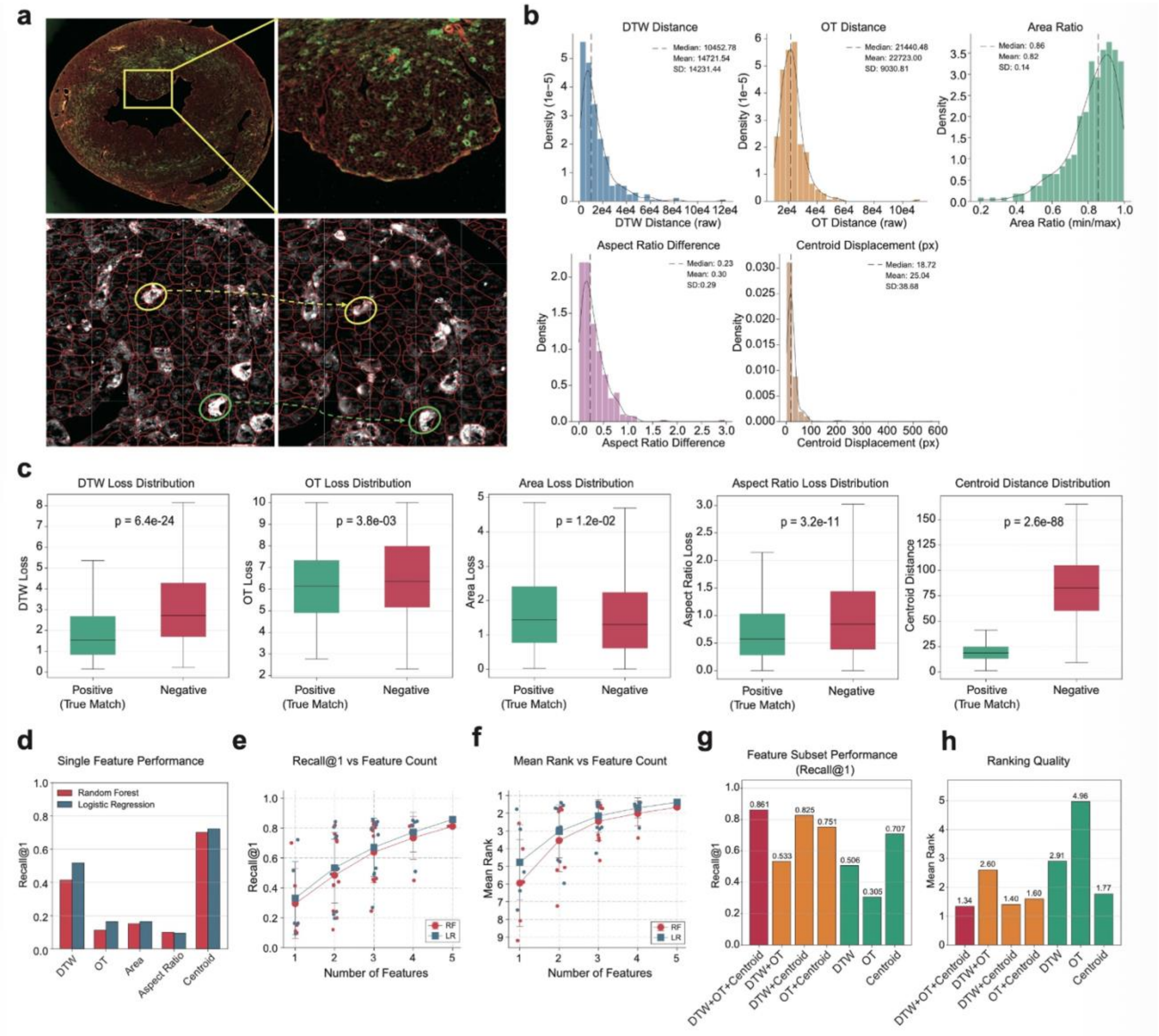

**a.** Ground-truth dataset generation from 10µm serial adult mouse heart cryosections with GFP sparsely expressed cardiomyocytes. Cellpose- proposed contours were curated by an HITL workflow; adjacent-section contours were retained as a true match only when they were morphologically similar, enclosed with GFP+ signal, and were unanimously approved by all specialists. A total of 340 adjacent-section true-match pairs were included in the final ground-truth dataset. **b.** Distributions of HiDTW-derived metrics for ground-truth true matches. **c.** Quantitative comparison of the five metrics between positive (true-match) and negative samples, where negative candidates were sampled from contours within a 200×200µm local window. **d.** Recall@1 (top-1 accuracy) using each individual feature (DTW, OT, area ratio, aspect ratio, centroid displacement). Bars compare Random Forest (RF) and Logistic Regression (LR) and evaluation used 5-fold GroupKFold cross-validation. **e-f.** Performance versus feature count. Ground-truth Recall@1 (**e**) and mean rank (**f**) across models using 1-5 features. Points denote individual combinations and error bars indicate mean ± SD. **g-h.** Core-feature subset analysis (LR). Performance of representative feature subsets, including DTW+OT, DTW+Centroid displacement, OT+Centroid displacement, DTW, OT, and Centroid displacement. Left, Recall@1; right, mean rank.

**Extended Data Fig. S3. Optimal transportation improves the performance of contour matching.**

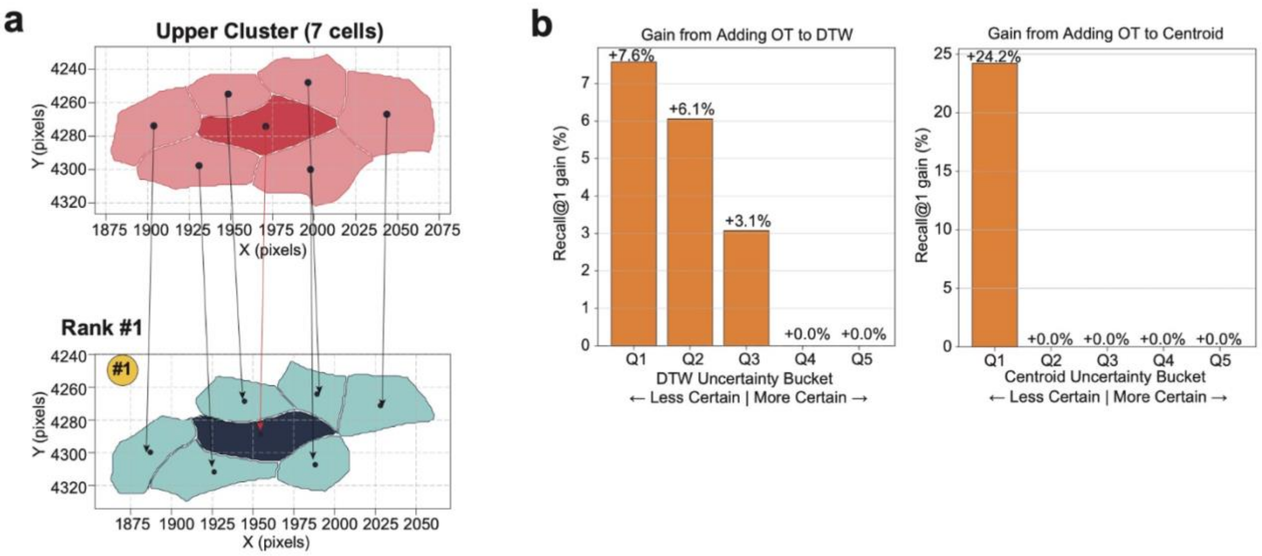


**a.** Representative example of OT-based selection of the corresponding cardiomyocyte in an adjacent section for a query cell (deep pink). Within a 100 × 100 px window, OT scores were computed and ranked across candidate cell clusters (blue) using an OT cost matrix derived from DTW distances between neighboring cells; candidate #1 was selected. **b.** OT gain under uncertainty. Improvement in Recall@1 after adding OT to a baseline feature. Ground-truth pairs were binned into five quantiles (Q1-Q5) by the margin between the top-1 and top-2 candidate scores from the baseline feature alone (smaller margin indicates lower certainty).

**Extended Data Fig. S4. Margin-based confidence control and stability analysis of HiDTW.**

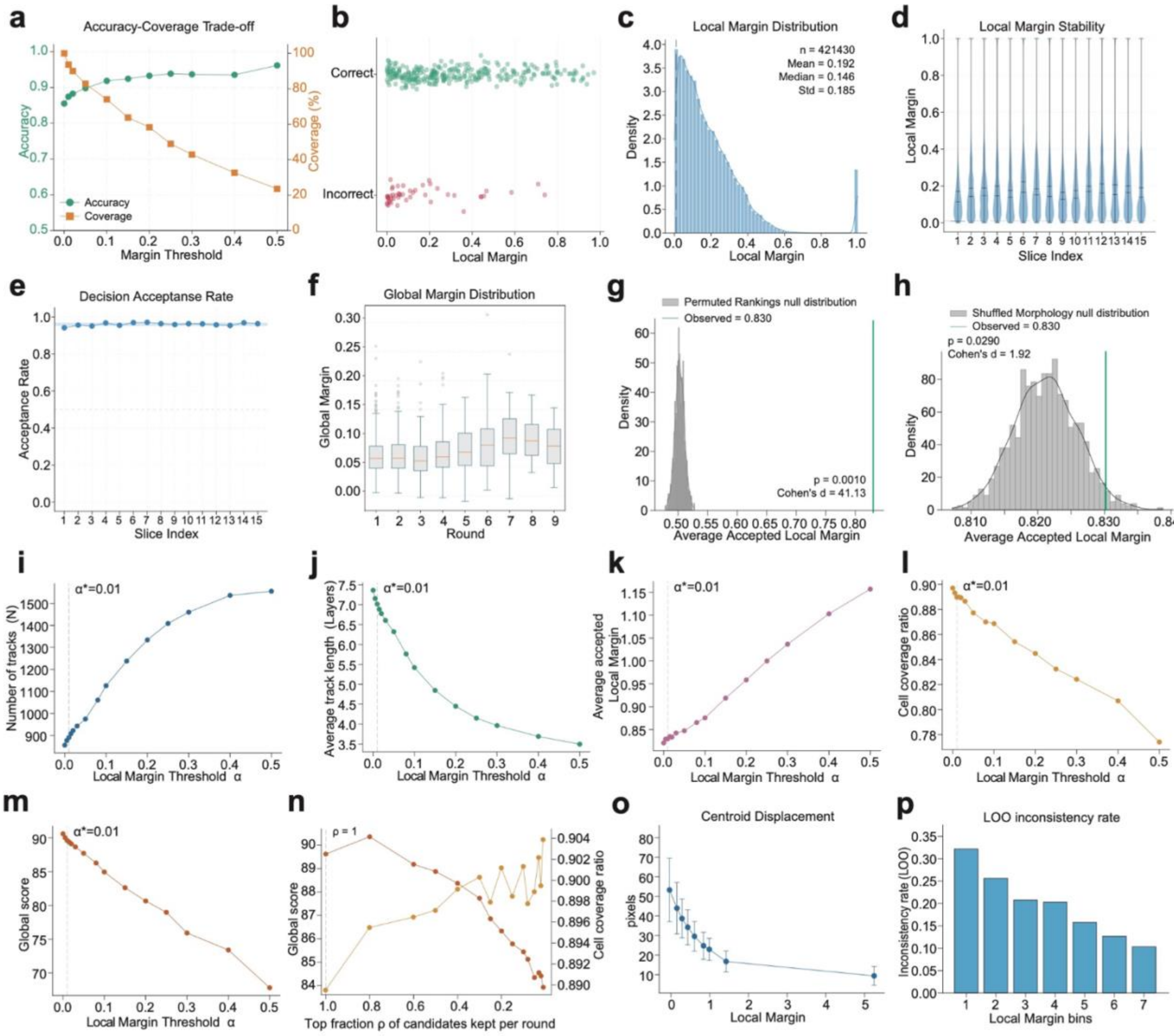


**a.** Accuracy-coverage trade-off for ground-truth pairs under local margin thresholding. **b.** Relationship between local margin and correctness. **c.** Distribution of the local margin at each stitching decision. **d.** Violin plots of local margins stratified by slice index, indicating stable margin scale and dispersion across adjacent slices. **e.** Per-slice decision acceptance rate, defined as the fraction of matches passing the local margin criterion during stitching. **f.** Boxplots of the global margin across iterative stitching rounds. All statistics in **c-f** are calculated from the best-scoring Bayesian-optimization run. **g-h.** Null distribution of the average accepted local margin under the permuted-rankings null model, which preserves candidate set size and the matching procedure but randomly permutes candidate-loss rankings; the vertical green line indicates the observed value from the real algorithm; empirical one-sided p values and Cohen’s d are shown. **i.** Correlation of the local margin threshold α and number tracks. **j.** Correlation of the local margin threshold α and average track length. **k.** Correlation of the local margin threshold α and the average accepted local margin. **l.** Correlation of the local margin threshold α and coverage ratio. **m.** Correlation of the local margin threshold α and global score. **n.** Sensitivity analysis of the candidate-retention parameter, ρ, where ρ denotes the top fraction of candidates retained at each round. **o.** External calibration of local margin against geometric consistency, showing that centroid displacement decreases as local margin increases. **p.** Leave-one-out (LOO) inconsistency rate stratified by local margin percentile bins.

**Extended Data Fig. S5. Bayesian optimization of matching hyperparameters and global score as a surrogate for accuracy.**

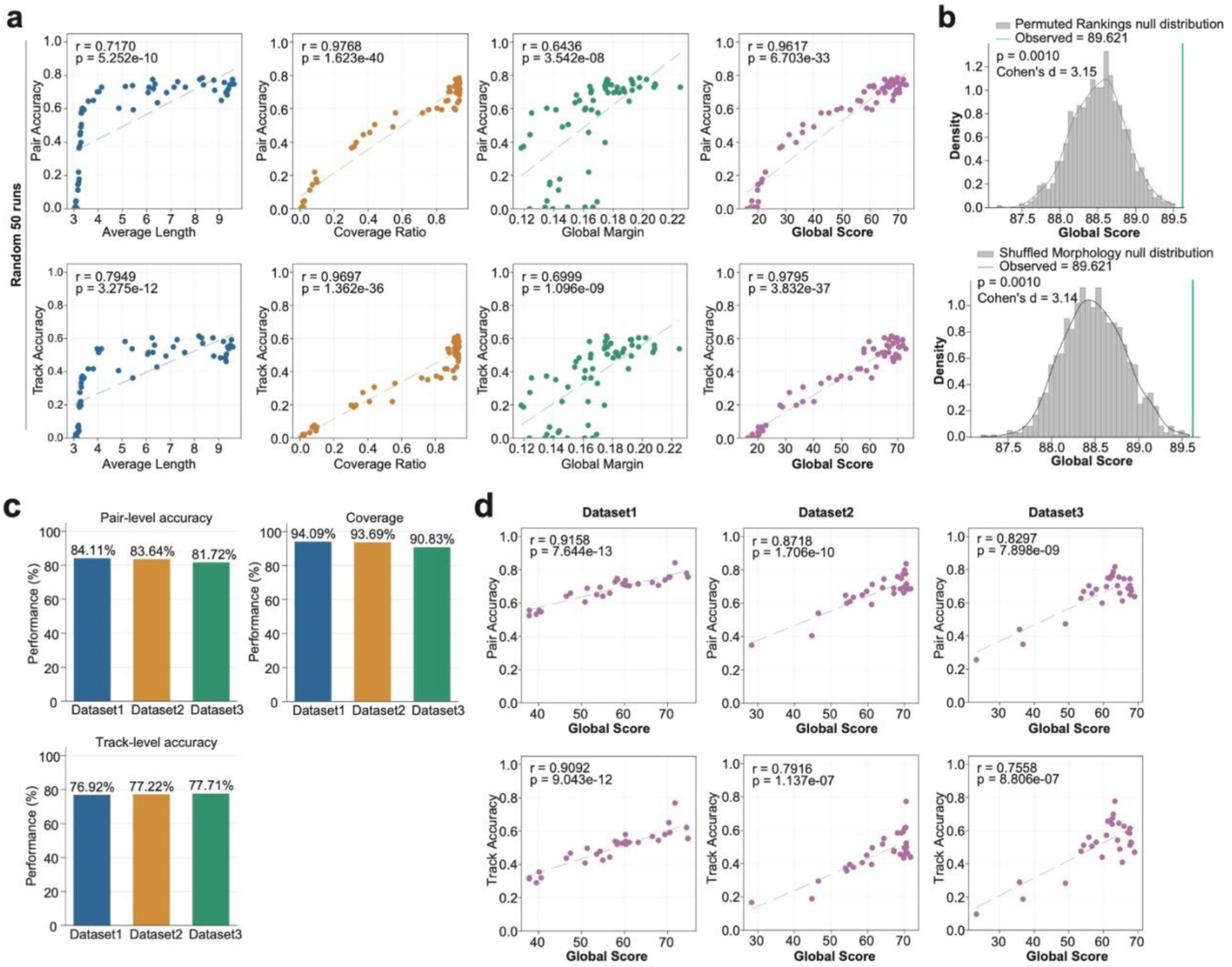


**a.** Random parameter sampling (50 runs) showing relationships between summary statistics (Average Length, Coverage Ratio, Global Average Margin, and Global Score) and Pair Accuracy (top) or Track Accuracy (bottom), evaluated on gold-standard pairs. Dashed lines indicate linear fits; Pearson r and P values are shown. **b.** Null distribution of the average accepted local margin and global score under the shuffled-morphology null model, which preserves centroid distance while randomizing morphology-related DTW/OT information; the vertical green line indicates the observed value from the real algorithm; empirical one-sided P values and Cohen's d are shown. **c.** The performance comparison of pair-level accuracy, track-level accuracy, and coverage ratio in three datasets. **d.** Validation on three ground-truth datasets. Scatter plots show the relationship between Global Score and Pair Accuracy (top) or Track Accuracy (bottom). Dashed lines indicate linear fits; Pearson's r and p values are shown.

**Extended Data Fig. S6. Quantitative evaluation of failure modes in the benchmark datasets.**

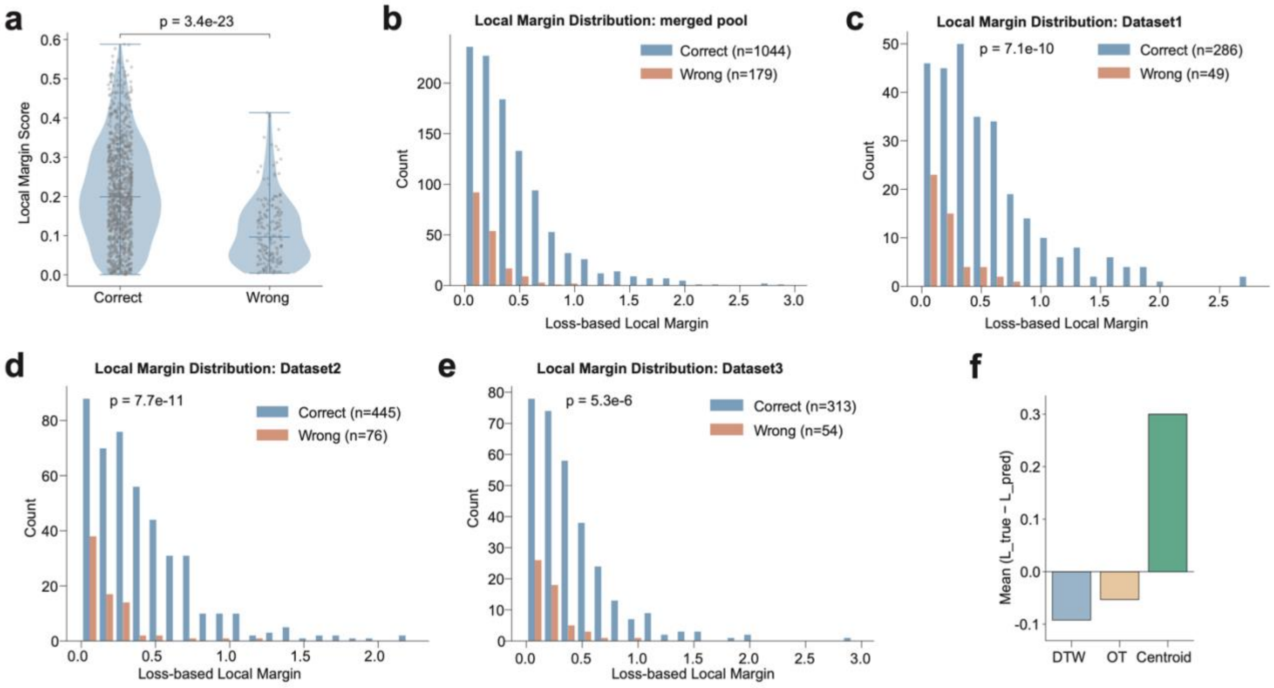


**a.** Violin plot of loss-based local margin in the pooled data; a two-sided t-test was used for statistical analysis. **b-e.** The local margin distribution in pooled and individual datasets. **f.** Mean contribution of DTW, OT, and centroid displacement (Centroid) components to the loss difference in incorrectly predicted matches.

**Extended Data Fig. S7. Representative cardiomyocyte track visualized across a two-photon microscopy z-stack acquired at 2μm axial steps.**

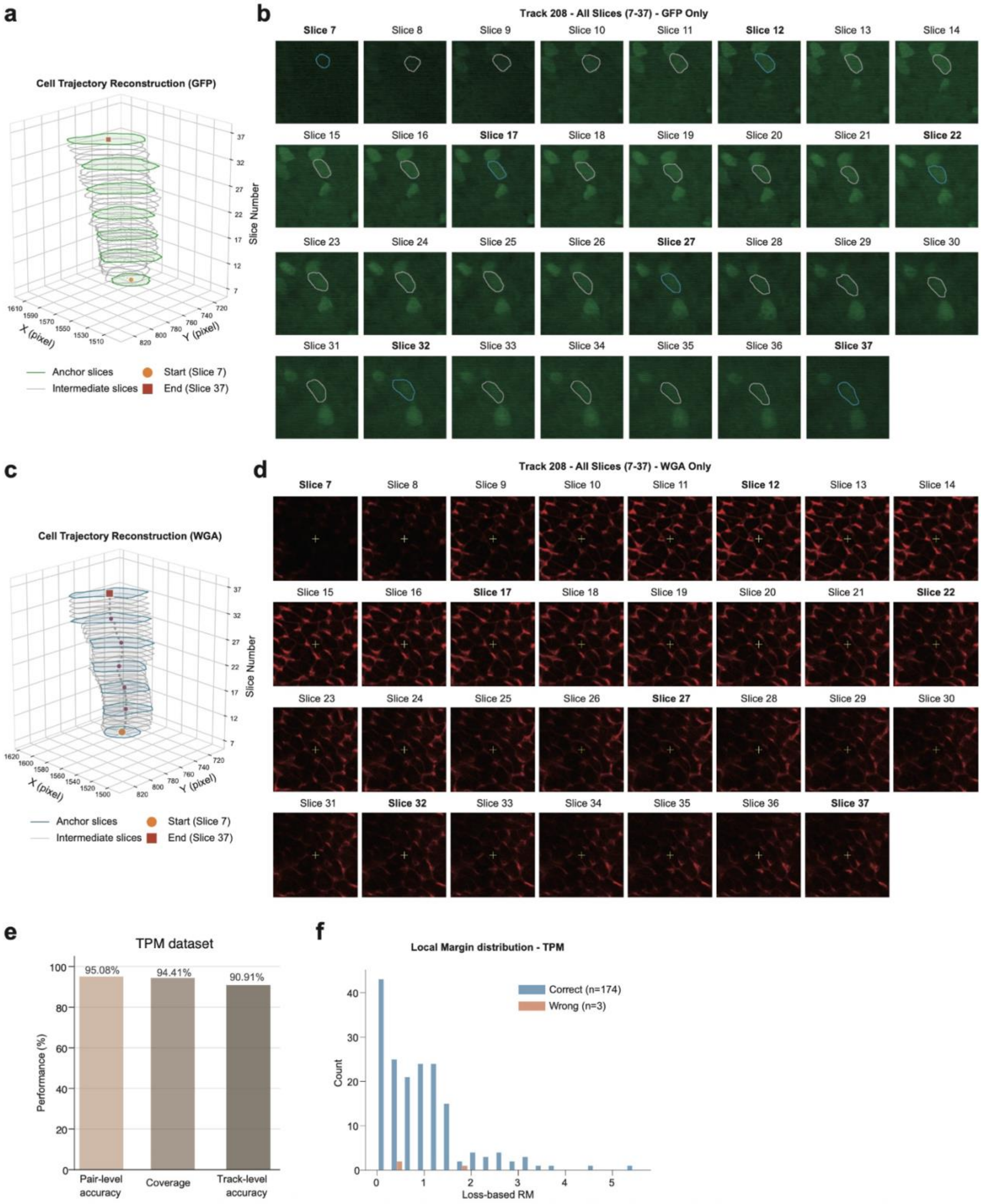


**a.** Three-dimensional reconstruction of a representative cardiomyocyte track using TPM-derived GFP signals. **b.** GFP channel showing the sparsely labeled cell with its contour overlaid across slices. **c.** Three-dimensional reconstruction of a representative cardiomyocyte track using TPM-derived WGA signals. **d.** Corresponding WGA membrane channel providing cell-boundary context. For stitching, we subsampled the stack at a 10μm interval by matching slices 7, 12, 17, 22, 27, 32, and 37. The intervening slices were then used to back-fill. **e.** Performance benchmark including coverage, pair-/track-level accuracy of HiDTW using TPM dataset. **f.** Local margin distribution in the TPM benchmark.

**Extended Data Fig. S8. Serial-section ST data generation and image-bead co-registration for post-MI mouse hearts.**

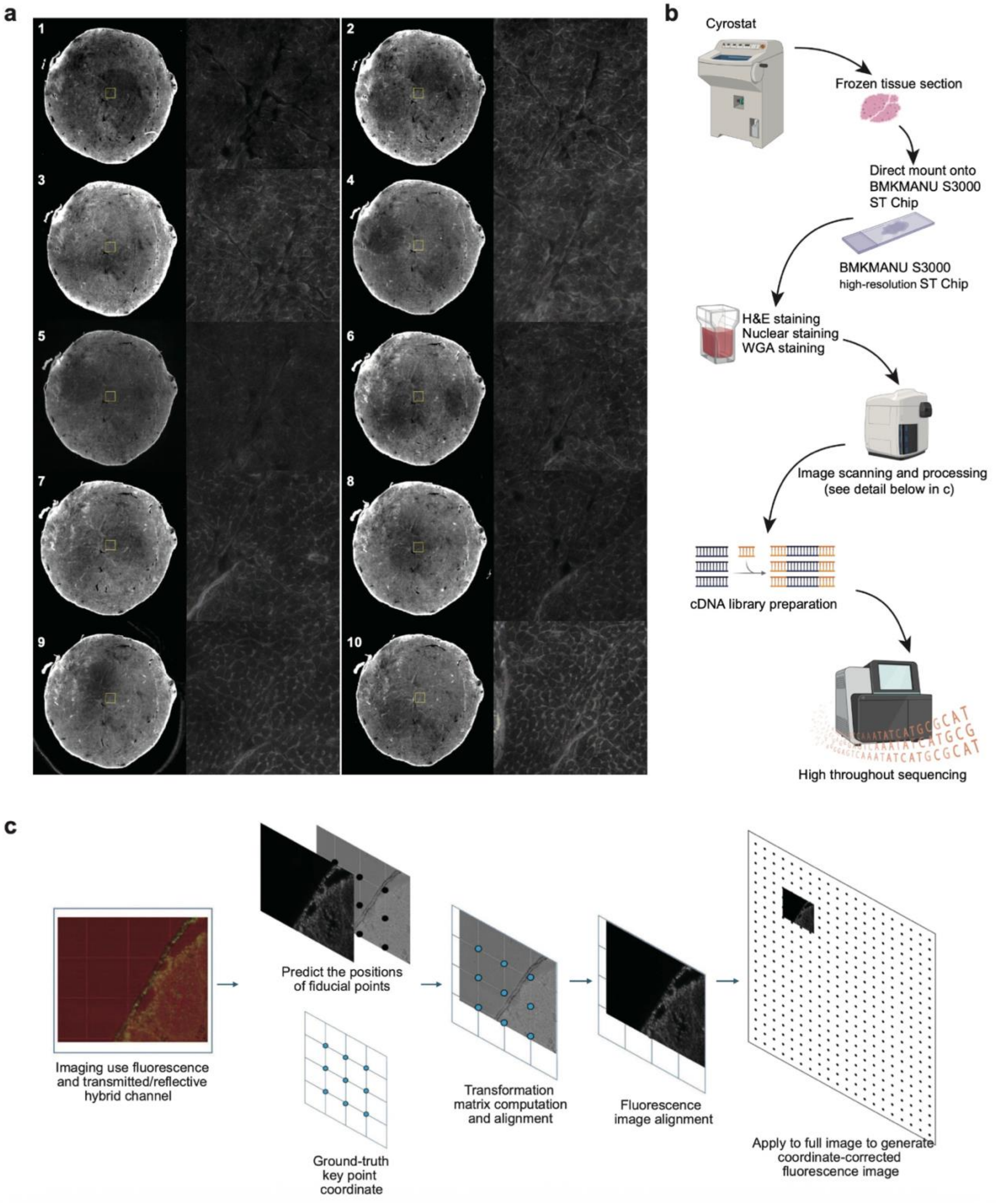


**a.** Ten sequential 10μm cryosections from mouse hearts after myocardial infarction (MI) surgery mounted onto single- cell-resolution (3.5μm) ST chips; representative staining images are shown for each section, with boxed regions indicating zoomed views. **b.**

Experimental workflow: cryosectioning, 14 direct mounting onto BMKMANU S3000 high-resolution ST chips, nuclear and WGA staining, image scanning/processing, cDNA library preparation, and high-throughput sequencing. **c.** Co-registration pipeline aligns staining images with barcoded nanobead coordinates. Fiducial points are detected, a transformation matrix is computed. The transform is applied to generate coordinate-corrected fluorescence images, enabling retrieval of corresponding RNA-sequencing reads in a common coordinate system.

**Extended Data Fig. S9. Macro-scale alignment of serial heart sections using SIFT-based rigid registration.**

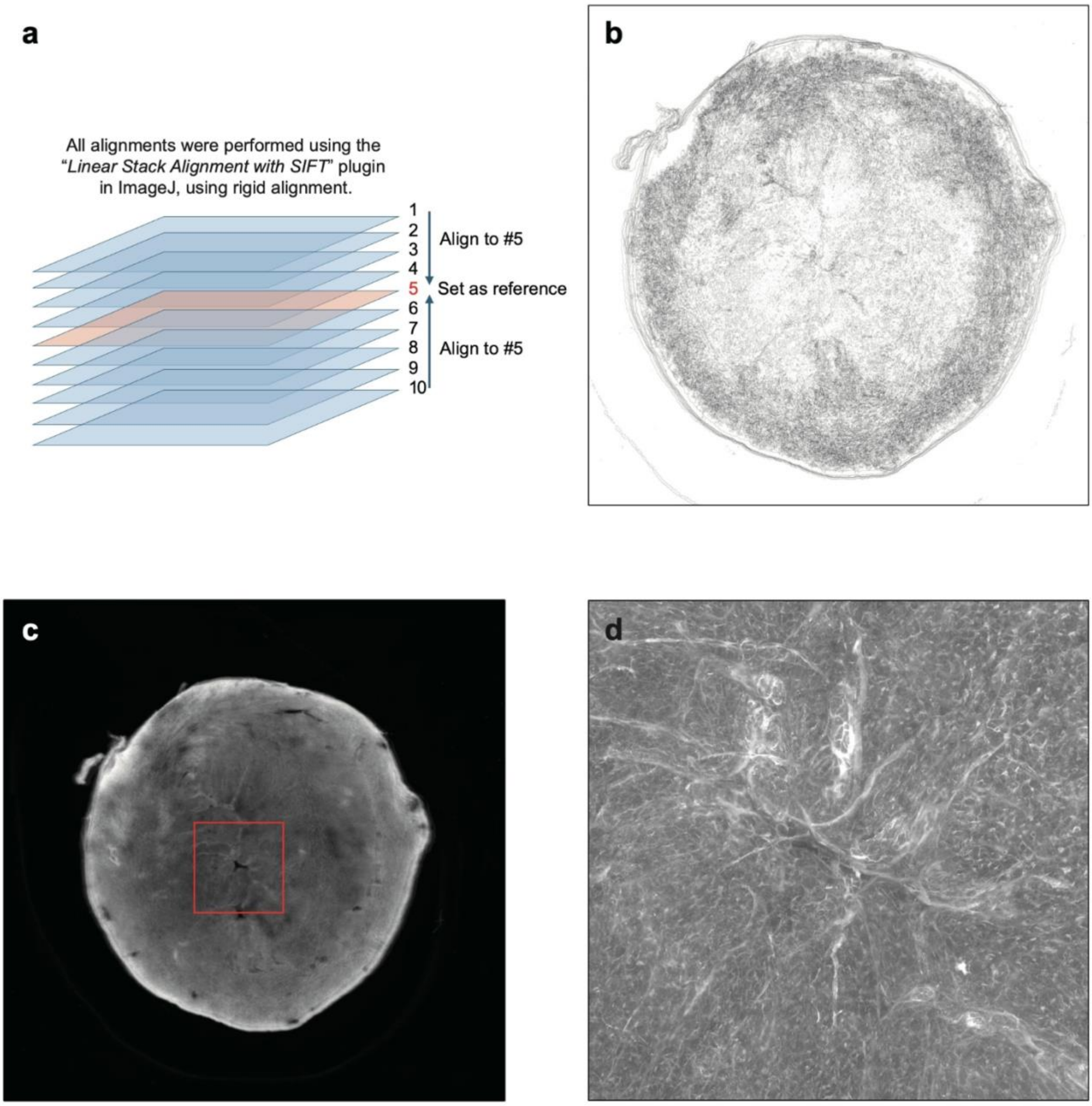


**a.** Schematic of macro-scale image alignment across 10 sequential sections using the ImageJ plugin Linear Stack Alignment with SIFT (rigid alignment), with section #5 used as the reference. **b.** Overlay of the aligned image stack showing consistent global registration across all 10 sections. **c.** Representative aligned whole-section image. **d.** Zoomed view from **c**, illustrating preserved local tissue features after macro-scale alignment.

**Extended Data Fig. S10. Track length and margin statistics across serial-section stitching.**

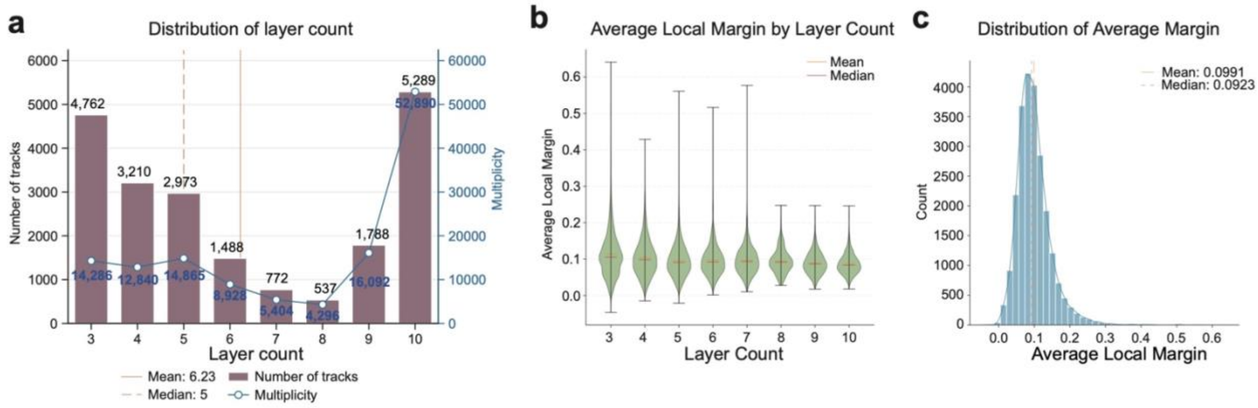


**a.** Distribution of 3D-VirtualCM cardiomyocytes' layer count (number of sections spanned by each 3D-VirtualCM cardiomyocyte) and tracks multiplicity, defined as the number of two-dimensional contours used to reconstruct each track. **b.** Violin plots of average local margin stratified by layer count. **c.** Global distribution of average local margin counts of all tracks.

**Extended Data Fig. S11. Benchmarking of HiDTW against representative cross-slice alignment methods and morphology-based baselines.**

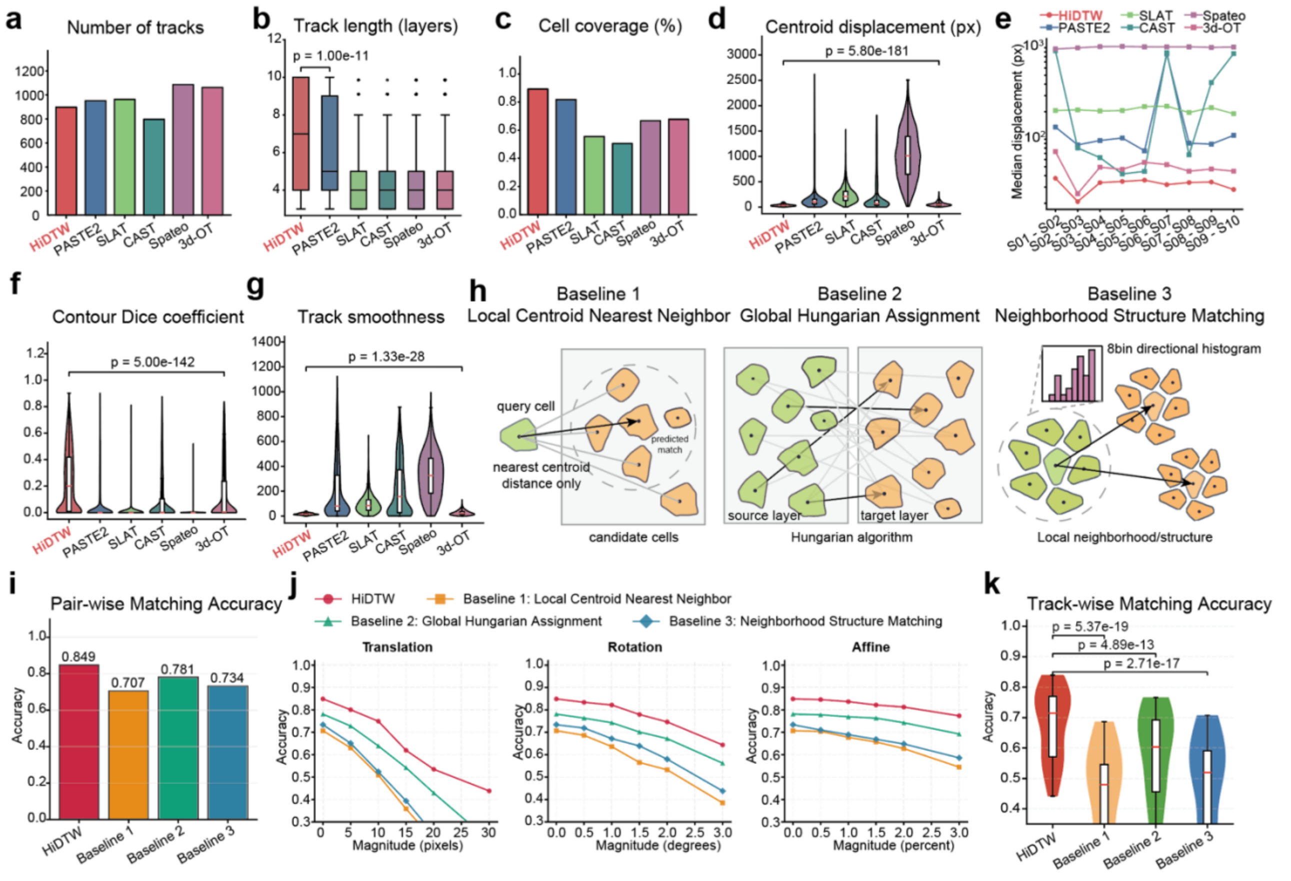


**a.** Number of reconstructed trajectories produced by each method. **b.** Distribution of trajectory-length comparisons across methods. **c.** Cell coverage comparison across methods. **d.** Distribution of centroid displacement between matched contours in adjacent sections. **e.** Comparison of cross-sectional median centroid displacement between layers using different methods. **f.** Distribution of contour Dice coefficients for matched contours in adjacent sections. **g.** Track smoothness assessment for all trajectories. Track smoothness was defined as the standard deviation of centroid displacement distances between adjacent layers within each track. **h.** Schematic overview of the three morphology-based baseline methods: local centroid nearest neighbor, global Hungarian assignment, and neighborhood structure matching. **i.** Overall matching accuracy of HiDTW and the three morphology-based baseline methods. **j.** Matching accuracy under increasing translation, rotation, and affine perturbations. **k.** Distribution of matching accuracies across 30 random combined perturbation settings composed of translation, rotation, and affine deformation. The p-values were calculated between HiDTW and the best-performing non-HiDTW method using a two-sided t-test.

**Extended Data Fig. S12. Estimation of non-cardiomyocyte gene contamination in 3D-VirtualCM.**

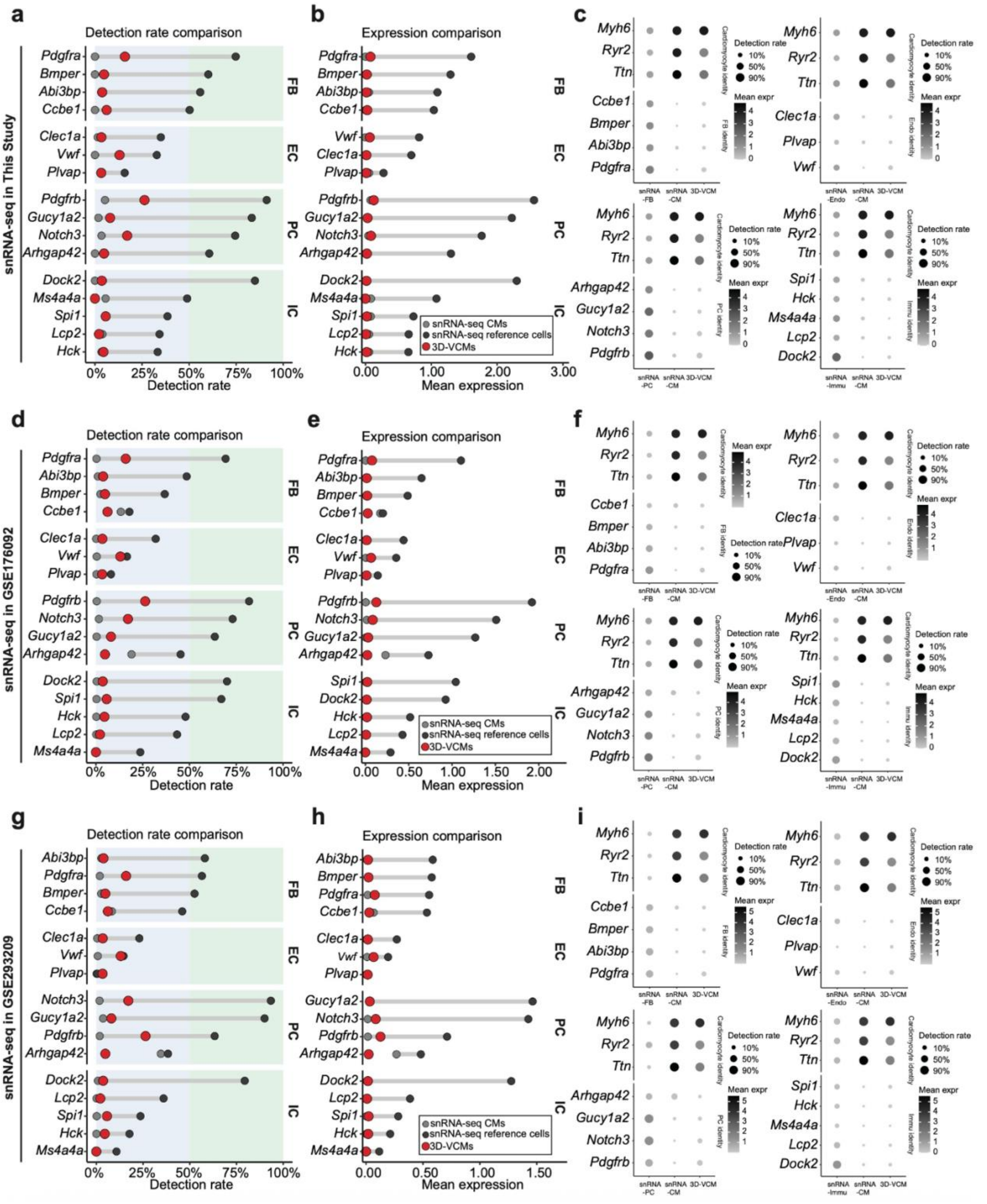


**a-b.** Detection rate **(a)** and mean expression **(b)** of representative non-cardiomyocyte lineage genes in 3D-VirtualCM cardiomyocytes (3D-VCM; red), compared with snRNA-seq cardiomyocytes (light grey) and snRNA-seq reference fibroblasts, endothelial cells, pericytes, and immune cells from this study (dark grey). Genes examined included fibroblast-associated genes (*Ccbe1, Bmper, Abi3bp, Pdgfra*), endothelial-associated genes (*Clec1a, Plvap, Vwf)*, pericyte-associated genes (*Arhgap42, Gucy1a2, Notch3, Pdgfrb*), immune-associated genes (*Spi1, Hck, Ms4a4a, Lcp2, Dock2*), and cardiomyocyte genes (*Myh6, Ryr2, Ttn*). **C.** Dot plot of lineage-gene expression across 3D-VCM cardiomyocytes, snRNA-seq cardiomyocytes (snRNA-CM), and snRNA-seq non-cardiomyocyte populations (snRNA-FB, snRNA-Endo, snRNA-PC, snRNA-Immune). Dot size denotes detection rate and color denotes mean expression. **d-f.** Same analyses as in **(a-c)** using an external snRNA-seq reference dataset (GSE176092). **g-i.** Same analyses as in **(a-c)** using an external snRNA-seq reference dataset (GSE293209).

**Extended Data Fig. S13. Integrated cardiomyocyte map and spatial gene modules.**

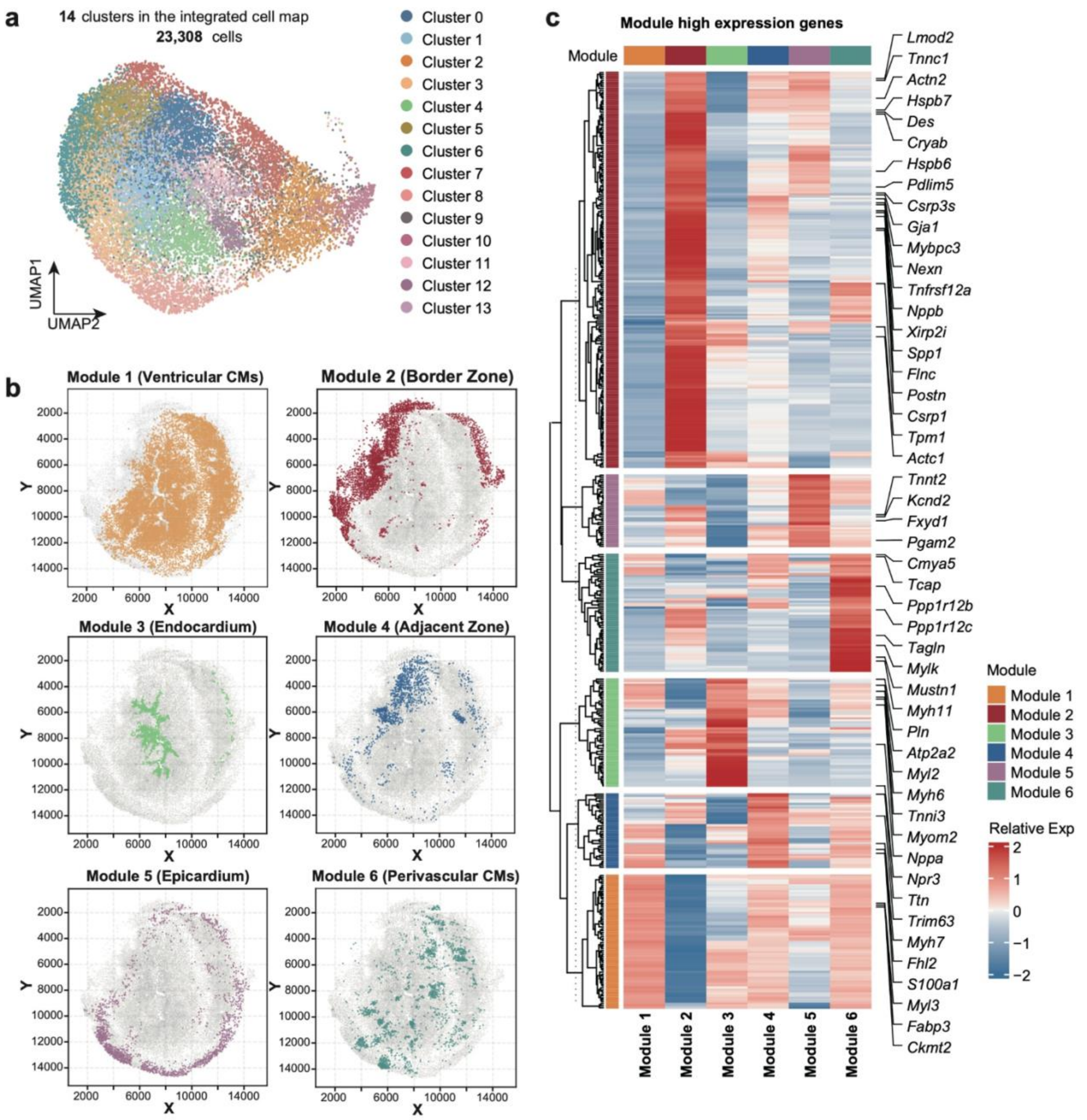


**a.** UMAP of the integrated cardiomyocyte atlas showing 14 clusters (23,308 cells). **b.** Spatial distributions of six cardiomyocyte gene modules over the tissue section: Module 1 (Ventricular CMs), Module 2 (Border Zone), Module 3 (Endocardium), Module 4 (Adjacent Zone), Module 5 (Epicardium), and Module 6 (Perivascular CMs). **c.** Heatmap of module high-expression genes. Rows show genes and columns show modules (Module 1-Module 6); colors indicate relative expression (z-scored), with selected marker genes annotated.

**Extended Data Fig. S14. Transcriptional differences of cardiomyocytes between Border Zone and Adjacent Zone.**

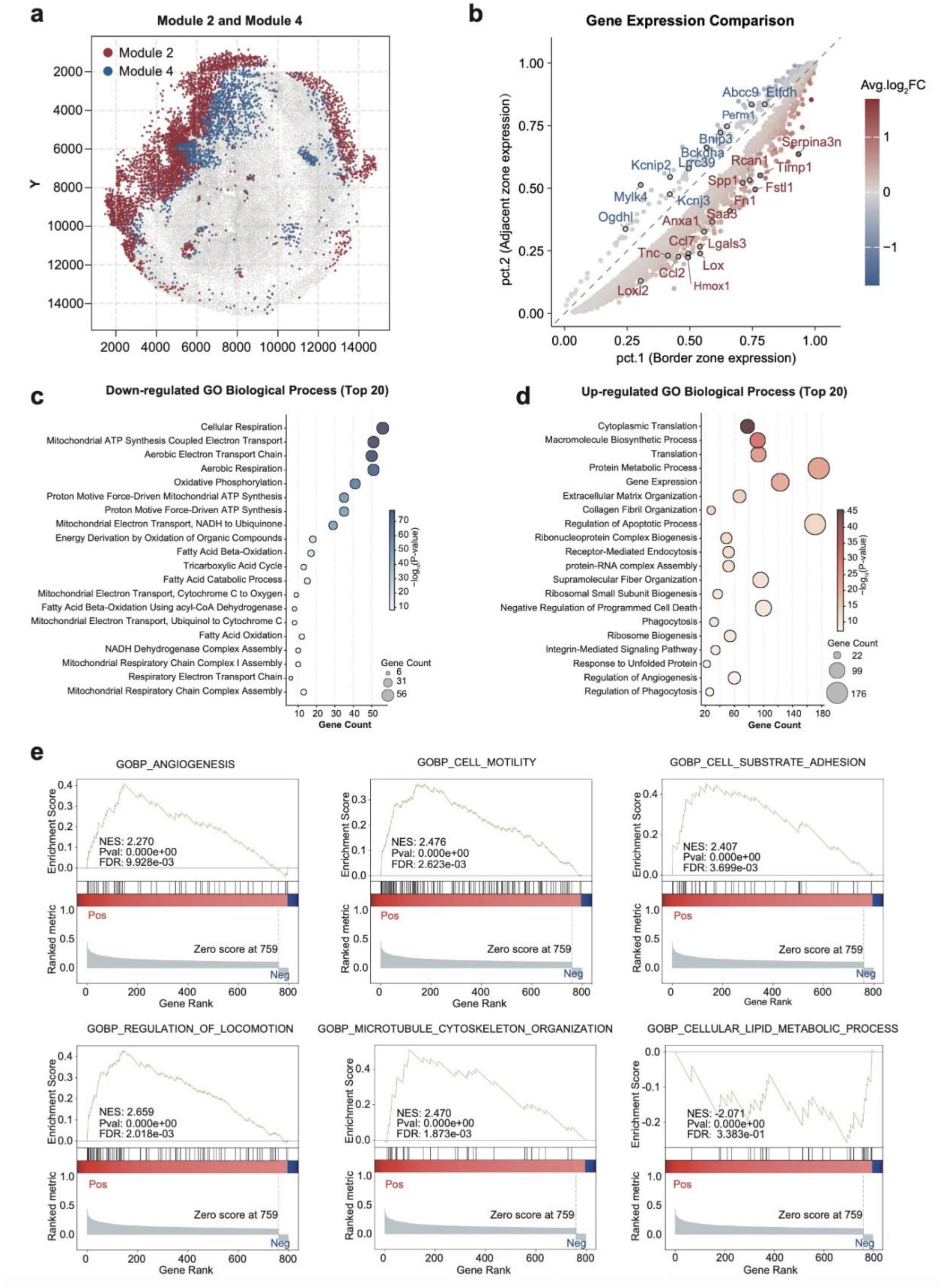

**a.** Spatial map showing the distribution of Module 2 (Border Zone CMs; red) and Module 4 (Adjacent Zone CMs; blue) overlaid on the tissue section. **b.** Gene-level comparison of expression between Module 2 and Module 4. Each point represents a gene plotted by its expression in Border Zone (pc.1; x-axis) versus Adjacent Zone (pc.2; y-axis); the dashed diagonal indicates equal expression. Colors denote average log2(fold change) (Module 2 vs Module 4). **c.** GO Biological Process enrichment of genes downregulated in Module 2 relative to Module 4. Dot size indicates gene count; color indicates significance. **d.** GO Biological Process enrichment of genes upregulated in Module 2 relative to Module 4. Dot size indicates gene count; color indicates significance. **e.** GSEA of differential expression of cardiomyocytes between Module 2 and Module 4.

**Extended Data Fig. S15. Spatial feature plots showing the expression patterns of canonical cell cycle genes across the tissue section.**

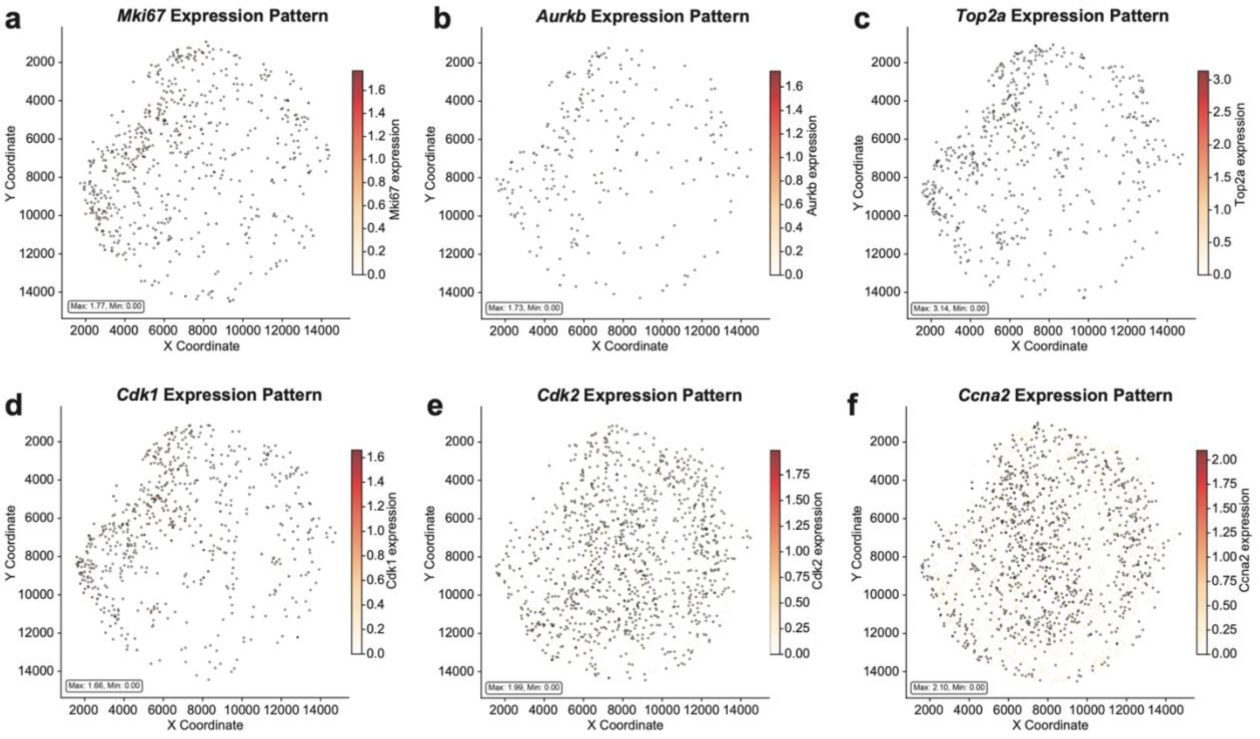


**a-f.** The gene expression distribution pattern of the indicated genes. Each point denotes a mapped 3D-VirtualCM positioned by its x-y coordinates. Highlighted points represent cells in the top 5% of expression for the indicated gene; color intensity reflects normalized expression among the highlighted cells.

**Extended Data Fig. S16. RNAscope-based identification of cycling cardiomyocytes using *Mki67* and *Aurkb*/*Top2a* with WGA-defined cell boundaries.**

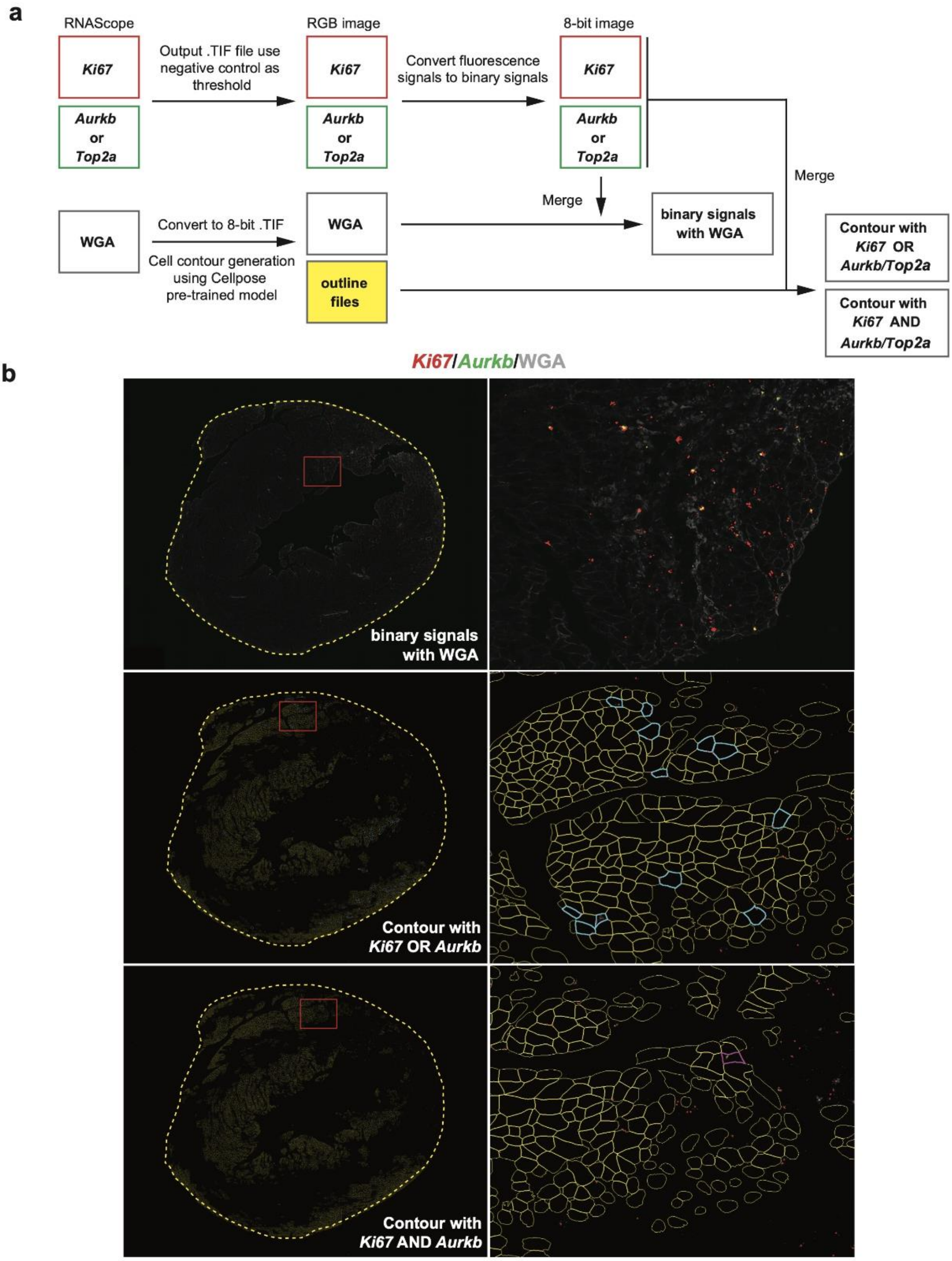

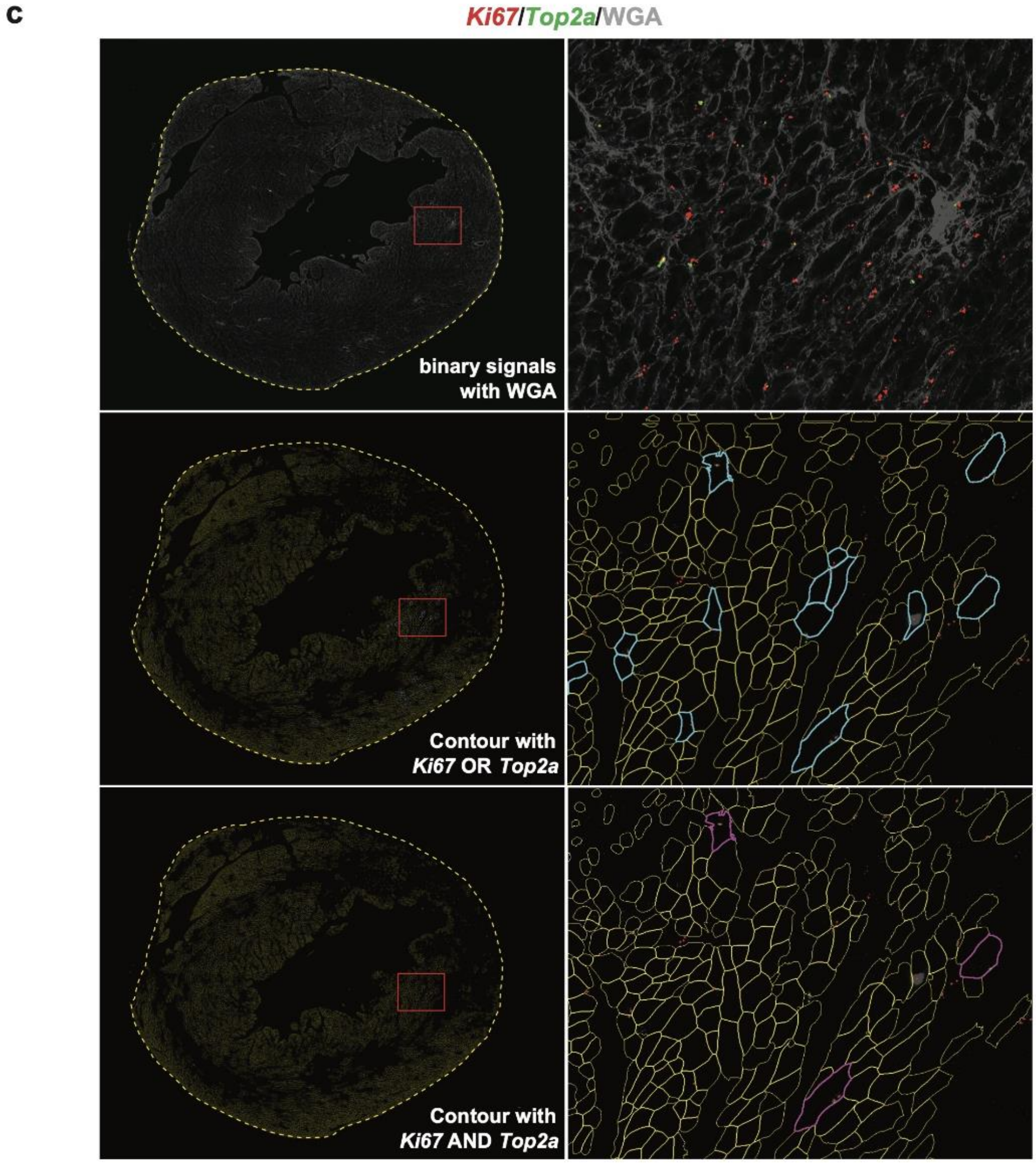


**a.** Image-processing workflow for calling cycling cardiomyocytes from RNAscope fluorescence. *Mki67* and *Aurkb* or *Top2a* channels were exported as RGB images, a threshold was set up using the negative control to generate binary masks, and merged with WGA images. Cell boundaries were obtained from WGA images by converting the images to 8-bit and segmenting them with Cellpose3 (pretrained model) to generate cell-contour files. RNAscope binary masks were intersected with WGA-derived contours to classify cells positive for Mki67 OR *Aurkb*/*Top2a* (union) or *Mki67* AND *Aurkb*/*Top2a* (intersection). **b.** Representative whole-section and zoomed views for *Mki67*/*Aurkb*/WGA. Top row, merged RNAscope binary masks overlaid with WGA; middle row, Cellpose-derived contours with cells called positive under the 25 *Mki67* OR *Aurkb* criterion; bottom row, the same contours with cells called positive under the *Mki67* AND *Aurkb* criterion. **c.** Representative whole-section and zoomed views for *Mki67*/*Top2a*/WGA, displayed as in b.

**Extended Data Fig. S17. Technical consistency across serial slices.**

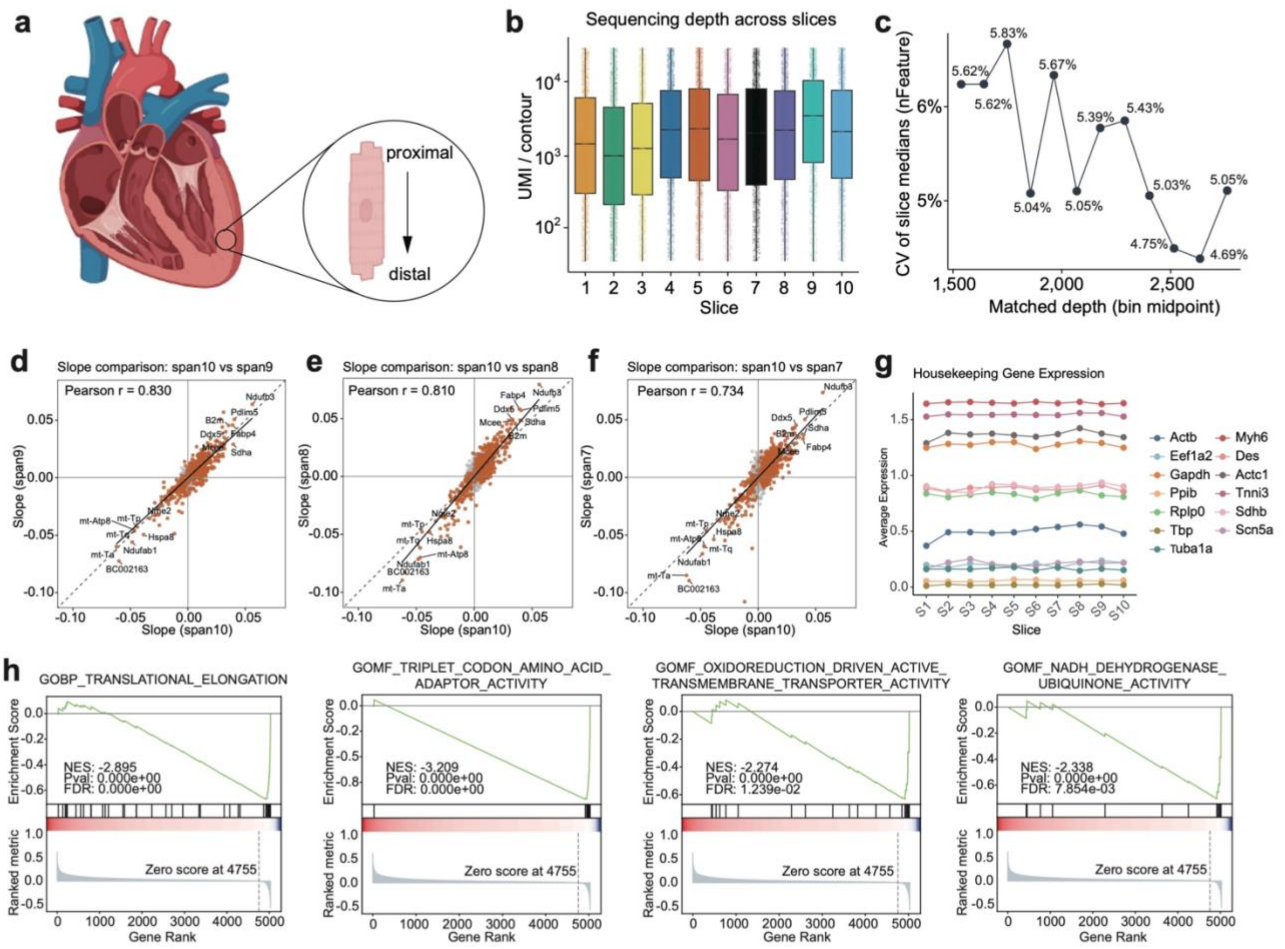


**a.** Schematic of the reconstructed cardiomyocyte axis across 10 serial heart sections, oriented from proximal to distal along the reconstruction span. **b.** Sequencing depth across slices for Module 1 spanning all 10 layers (span10 3D-VirtualCM cardiomyocytes). Boxplots show UMI counts per reconstructed contour for each slice. **c.** Depth-matched technical consistency across slices. Contours were stratified into 12 global sequencing-depth quantile bins; within each bin, the coefficient of variation (CV) of the per-slice median detected genes across the 10 slices was computed, indicating low across-slice variability after depth matching. **d-f.** Longitudinal expression-gradient slope robustness across reconstruction spans. For each gene, a linear model was used to estimate the expression-gradient slope along the reconstructed layer axis. Gene-wise slopes computed from span10 3D-VirtualCM cardiomyocytes were used as reference (x-axis) and compared against slopes from span9 **(d)**, span8 **(e)**, and span7 **(f)** cardiomyocytes (y-axis). Each panel reports Pearson correlation r and p-value. **g.** Stability of housekeeping gene expression across serial slices. (h). Representative GSEA plots for downregulated pathways. **h.** The representative gene ontology biological processes (GO-BP) of the downregulated functional pattern in the proximal end. NES, normalized enrichment score; FDR, false discovery rate;

**Extended Data Fig. S18. The Heatmap of featured genes at the proximal versus distal ends of adult cardiomyocytes.**

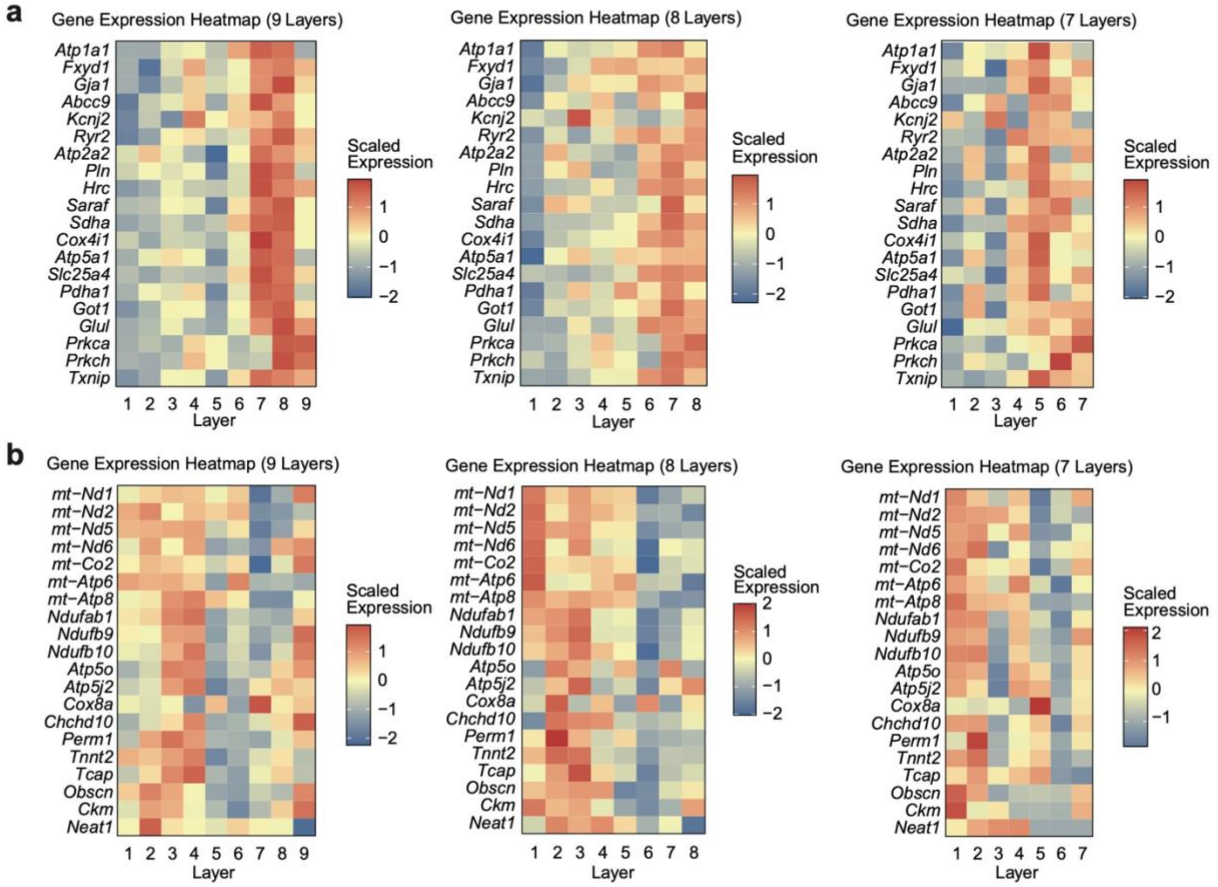


**a.** Row-scaled gene-expression heatmaps of featured distal-end enriched genes in adult cardiomyocytes spanning 9, 8, or 7 layers (left to right). **b.** Row-scaled gene-expression heatmaps of featured proximal-end enriched genes in adult cardiomyocytes spanning 9, 8, or 7 layers (left to right).
