## Supplementary material for "Three-dimensional Virtual Adult Cardiomyocyte Transcriptomics": All suppl files: 518649_3_related_ms_5215329_tkd6d1_convrt.pdf

**Supplementary Table S1. Representative methods compared with 3D-VirtualCM at the level of individual reconstructed cells**

| Approach | Primary 3D strategy | Membrane-based cell segmentation | Serial-section reconstruction | Cell-matched transcriptomes | Intracellular RNA heterogeneity | Suitability for large cells |
|---|---|---|---|---|---|---|
| **CODA** *Kiemen et al., 2022* | **Tissue reconstruction:** Serial histology registration and tissue-label interpolation | N/A | ✓ Tissue-level serial-section reconstruction | — No matched transcriptome | N/A | △ Tissue-scale; no elongated-cell identity tracking |
| **PASTE** *Zeira et al., 2022* | **Tissue reconstruction:** Optimal-transport alignment of serial ST slices | N/A | ✓ Slice alignment, not individual-cell tracking | △ Integrates spot-level expression without stable cell identity | N/A | N/A |
| **3d-OT** *Dai et al., 2026* | **Tissue reconstruction:** Deep geometry-aware spatial multi-omics slice alignment | N/A | ✓ 3D reconstruction through heterogeneous slice alignment | △Supports spot/cell correspondence for alignment | N/A | △ General tissues; not tailored to multinucleated cardiomyocytes |
| **Functional connectomics** *MICrONS Consortium, 2025* | **Connectomics**: Serial electron-microscopy volume assembly | ✓ Membrane contrast defines cellular boundaries | ✓ High-fidelity tracing across sections | N/A | N/A | △ Excellent for extended processes, but costly and low throughput |
| **Tissue clearing + LSFM** *Dodt et al., 2007; Chung et al., 2013* | **Connectomics**: Optical imaging of intact cleared tissue | △ Requires membrane or cell-specific labels | △ Direct z-stack rather than section stitching | N/A | △ Targeted RNA or protein localization only | ✓ Well suited to large labeled cells |
| **CaMVIA-3D** *Wei et al., 2026* | **volumetric imaging:** Confocal z-stacks of thick cardiac sections | ✓ WGA-guided Omnipose segmentation | △ Direct imaging of thick tissue sections | N/A | N/A | ✓ Optimized for adult cardiomyocytes |
| **3D-VirtualCM** *This study* | **Serial ST alignment plus membrane-contour tracking** | **✓ WGA-guided ACMR segmentation** | **✓ HiDTW cell-identity tracking** | **✓ Cell-matched volumetric transcriptomes** | **✓ Longitudinal intracellular RNA gradients*** | **✓ Optimized for large, elongated, multinucleated adult cardiomyocytes** |

✓ native or direct capability △ partial, conditional or indirect capability — absent or outside the method's design scope N/A not available

* **Resolution note.** For 3D-VirtualCM, intracellular RNA mapping is constrained by the 10-μm axial sampling interval and the lateral resolution of the spatial transcriptomics platform.
