## Supplementary material for "Three-dimensional Virtual Adult Cardiomyocyte Transcriptomics": All suppl files: 518649_3_supp_5189940_t6d5v8_convrt.pdf

### Supplementary Methods

#### 1. Cell Contour Feature Extraction and Adjacency Graph Construction

To reconstruct 3D trajectories from 2D sections, we first convert each cell boundary into a standardized contour representation with a fixed number of ordered points, then compute per-cell geometric descriptors, and finally construct an intra-slice adjacency graph for neighborhood-based OT computation. This design separates three roles of the contour data: geometric summarization, contour-to-contour shape comparison, and within-slice spatial proximity estimation.

##### 1.1 Contour Reading and Equal-distance Resampling

For each cell $j$, we represent its original contour as a point set $P_j = \{p_{i,j} = (x_{i,j}, y_{i,j})\}_{i=1}^{n_j}$, where $n_j$ denotes the number of contour vertices for cell $j$ and may vary across cells. Since the raw contour vertices may not provide a consistent starting point or ordering across cells, we convert them into a cyclically ordered contour sequence before subsequent processing. Specifically, points are sorted in ascending order of the centroid-referenced clockwise polar angle defined as:

$$\theta_{i,j}^{\mathrm{cw}} = \mathrm{mod}\left\{\, \mathrm{atan2}\left[x_{i,j} - \bar{x}_j, -\left(y_{i,j} - \bar{y}_j\right)\right], 2\pi \,\right\}.$$

Here, the arguments of atan2 are intentionally ordered according to the image-coordinate convention, such that the upward direction corresponds to zero angle and the contour is traversed clockwise. To further ensure consistency across cells, the ordered sequence is circularly shifted so that its starting point is the vertex closest to the topmost direction (12 o'clock in image coordinates). In this way, we obtain a reordered contour sequence $\left(q_{1,j}, \ldots, q_{n_j,j}\right)$. We close the contour by setting $q_{n_j+1,j} = q_{1,j}$ and index the ordered contour by cumulative arc length:

$$s_{i,j} = \sum_{k=1}^{i} \left\| q_{k+1,j} - q_{k,j} \right\|_2, \quad i = 1, \ldots, n_j\,.$$

This arc-length parameterization provides a common one-dimensional coordinate along the closed contour. To facilitate downstream contour-based computation, each cell contour is then resampled to a fixed number $N = 200$ of boundary points used throughout this work, so that all cells are represented by fixed-length point sequences, Unless otherwise specified, $N = 200$ was used for all contour-matching computations; a different sampling density was used only for visualization-specific mesh rendering. Specifically, we resample $N$ uniformly spaced positions on $\left[0, s_{n_j,j}\right]$ via linear interpolation of $x$ and $y$; if $s_{n_j,j} = 0$ (a degenerate zero-perimeter contour), we replicate the single point $N$ times. For translation invariance, we center the sampled contour by subtracting its centroid and get $\hat{P}_j = \{\hat{p}_{1,j}, \hat{p}_{2,j}, \dots, \hat{p}_{N,j}\}$. This normalization removes absolute position while preserving contour shape and orientation in the image coordinate system.

**1.2 Geometric features.**

From the original contour, we compute per-cell features including centroid, polygon area (via the shoelace formula, with absolute value), and an ellipse-based aspect ratio obtained by least-squares fitting using OpenCV fitEllipse. We compute these summary descriptors from the original contour rather than the resampled one, so as to avoid interpolation-induced bias in area- and shape-related measurements. Ellipse fitting was attempted only when sufficient contour vertices were available. If ellipse fitting fails or the minor axis is zero, we set the aspect ratio to 1 as a neutral fallback value.

**1.3 Intra-slice adjacency graph**

To construct the intra-slice adjacency graph, we compute the boundary-to-boundary distance between cells $i$ and $j$ as the minimum Euclidean distance over all pairs of their resampled contour points in the original image coordinates, denoted by $d_{ij}$. Distances are evaluated in the original coordinate system so that spatial proximity reflects the true within-slice arrangement of cells. Cells $i$ and $j$ are defined as neighbors if $d_{ij}$ < 5 pixels; This fixed threshold was used to identify physically adjacent or nearly touching cell boundaries while excluding more distant cells. For each cell, we stored both the neighbor IDs and the corresponding neighbor count.

**2. Dynamic Time Warping (DTW)**

To quantitatively measure contour similarity between adjacent slices, we used a Soft-DTW-based contour dissimilarity measure, which is a differentiable relaxation of Dynamic Time Warping (DTW), and computed it using a CUDA-accelerated implementation for efficient and scalable batch computation. For a cell $c_i^z$ on slice $z$ and a candidate cell $c_j^{z+1}$ on slice $z+1$, we extract their 2D boundary contours, each uniformly resampled to $N$ points (Section 1) and get $\hat{P}_i^z, \hat{P}_j^{z+1}$. Here, $\hat{P}_i^z$ and $\hat{P}_j^{z+1}$ denote the centered fixed-length contour sequences obtained by the preprocessing procedure in Section 1, so the subsequent comparison is translation-invariant and operates on standardized boundary representations.

Because the contour points have been reordered to share a common traversal direction and a consistent starting point, the two cell boundaries can be used as comparable ordered sequences for DTW-based alignment. This preprocessing step is important for DTW-based matching, since it reduces artificial discrepancies caused by arbitrary contour indexing and allows the distance to reflect genuine shape differences along the boundary.

The DTW distance between the two contours is defined as:

$$D_{\text{DTW}}\left(c_i^z, c_j^{z+1}\right) = \text{SoftDTW}\left(\hat{P}_i^z,\, \hat{P}_j^{z+1}\right).$$

Here, $D_{\text{DTW}}$ denotes the Soft-DTW-based contour dissimilarity used in this study, rather than the classical hard-minimum DTW distance. In this work, Soft-DTW serves as the practical DTW-based contour dissimilarity measure used in all downstream matching steps. Compared with a strict point-to-point comparison, this formulation is more tolerant to local boundary deformation and minor local mismatches in contour parameterization. Generally, a smaller $D_{\text{DTW}}$ value indicates higher contour similarity between the two candidate cells.

#### 3. Optimal Transport (OT)

To complement the DTW contour distance with local context, we introduce a neighborhood distance based on optimal transport (OT), which measures the overall matching quality between the two surrounding neighborhood blocks. For a cell $c_i^z$ on slice $z$ and a candidate cell $c_j^{z+1}$ on slice $z+1$, their neighborhood blocks (including themselves) are denoted by $\mathcal{C}_i^z,\ \mathcal{C}_j^{z+1}$. That is, each neighborhood block is defined as the central cell together with its intra-slice adjacent neighbors from Section 1. The OT term is therefore designed to capture local structural consistency around a candidate cross-slice match, rather than only the similarity between the two central cells themselves.

**3.1 Cost matrix based on Soft-DTW**

All contours are preprocessed and uniformly resampled to $N$ points as described in Section 1. Denote the preprocessed contour point sequences of $a_r \in \mathcal{C}_i^z$ and $b_s \in \mathcal{C}_j^{z+1}$ by $\hat{P}(a_r)$ and $\hat{P}(b_s)$, respectively. Here, $\hat{P}(\cdot)$ denotes the centered fixed-length contour representation obtained after contour reordering, arc-length resampling, and centroid subtraction. Then the pairwise shape-matching cost is defined as:

$$K_{rs} = \text{SoftDTW}\left(\hat{P}(a_r),\ \hat{P}(b_s)\right),$$

and the cost matrix is $\boldsymbol{K} = (K_{rs})$. In other words, each entry of $\boldsymbol{K}$ measures the contour dissimilarity between one cell in the source neighborhood block and one cell in the target neighborhood block. This matrix serves as the ground cost for the OT problem defined below.

**3.2 Area-weighted mass distribution and entropic OT**

Because contour measurements from very small segmented objects are more easily affected by discretization artifacts and local segmentation noise. We therefore assign area-proportional masses to cells so that the OT plan favors more reliably segmented contours.

Let $A(a_r)$ and $A(b_s)$ be the polygon areas (Section 1) of $a_r$ and $b_s$. Let $m = |\mathcal{C}_i^z|$ and $n = |\mathcal{C}_j^{z+1}|$ denote the numbers of cells in the two neighborhood blocks, respectively. The source and target masses are defined as:

$$\alpha_r = \frac{A(a_r)}{\sum_{a_{r'} \in \mathcal{C}_i^z} A(a_{r'})}, \qquad \beta_s = \frac{A(b_s)}{\sum_{b_{s'} \in \mathcal{C}_j^{z+1}} A(b_{s'})}.$$

If $m = n = 1$ (i.e., both neighborhoods contain only the central cell itself, with no adjacent neighbors), the OT distance is zero directly. This is a design choice: when no neighborhood context is available on either side, the OT term is treated as non-informative and therefore contributes neither penalty nor reward to the final matching score.

Given the cost matrix $\boldsymbol{K} = (K_{rs})$, the source mass distribution $\boldsymbol{\alpha} = (\alpha_r)_{r=1}^{m}$, and the target mass distribution $\boldsymbol{\beta} = (\beta_s)_{s=1}^{n}$, we compute the entropy-regularized OT plan $\boldsymbol{T}^*$ using a GPU-accelerated Sinkhorn algorithm:

$$\boldsymbol{T}^* = \arg\min_{\boldsymbol{T}\in\mathbb{R}_{\geq 0}^{m\times n}} \langle \boldsymbol{T}, \boldsymbol{K} \rangle + \varepsilon \sum_{r=1}^{m}\sum_{s=1}^{n} T_{rs}(\log T_{rs} - 1) \quad s.t. \quad \boldsymbol{T}\mathbf{1}_{\boldsymbol{n}} = \boldsymbol{\alpha}, \quad \boldsymbol{T}^{\top}\mathbf{1}_{\boldsymbol{m}} = \boldsymbol{\beta}.$$

Here, $\boldsymbol{T} = (T_{rs}) \in \mathbb{R}_{\geq 0}^{m\times n}$ is the transport plan between the two neighborhood distributions, where $T_{rs}$ denotes the amount of mass transported from source cell $a_r$ to target cell $b_s$. The marginal constraints enforce consistency with the area-weighted masses $\boldsymbol{\alpha}$ and $\boldsymbol{\beta}$. The entropic regularization term improves numerical stability and enables efficient computation via Sinkhorn iterations.

The adaptive regularization strength is defined as:

$$\varepsilon = \max\big(\varepsilon_0,\ 0.05 \cdot \mathrm{median}(\boldsymbol{K})\big), \quad \varepsilon_0 = 0.01\,.$$

The floor $\varepsilon_0 = 0.01$ prevents numerical instability when the median cost is near zero. In implementation, $\varepsilon$ is additionally capped at 10000 to avoid over-regularization. The adaptive form further keeps the regularization strength on a scale comparable to the current neighborhood cost matrix, which improves robustness across cell pairs with different contour dissimilarity levels. The neighborhood OT cost is then:

$$D_{\mathrm{OT}}\big(c_i^z, c_j^{z+1}\big) = \langle \boldsymbol{T}^{\star}, \boldsymbol{K} \rangle = \sum_{r,s} T_{rs}^{\star} K_{rs}\,.$$

Thus, $D_{\mathrm{OT}}$ summarizes how well the local neighborhood around $c_i^z$ can be matched to that around $c_j^{z+1}$ under the Soft-DTW cost and the mass distributions. Smaller values indicate better neighborhood-level agreement between the two candidate cells.

##### 3.3 Additional analyses of the OT-based neighborhood score

Resampling each contour to a fixed number of boundary points makes the Soft-DTW comparison well defined by equalizing sequence length, but it does not by itself remove scale dependence. To assess the practical role of cell size in the present formulation, we performed additional control analyses. In a 2×2 ablation crossing contour scale treatment (raw versus $\sqrt{\text{area}}$-normalized contours) with OT marginal definition (area-weighted versus uniform), while keeping candidate generation, gold-standard definition, and the quantile-normalized scoring framework unchanged (n=851), explicit contour scale normalization substantially reduced the Top-1 hit rate against the gold-standard target, whereas replacing area-weighted OT marginals with uniform marginals under raw Soft-DTW produced only a small change. Specifically, the Top-1 hit rate decreased from 0.744 to 0.378 (McNemar p = $1.23 \times 10^{-53}$) after contour scale normalization, but changed only from 0.7438 to 0.7368 (McNemar p = 0.118) when area-weighted marginals were replaced by uniform marginals. These results indicate that absolute cross-sectional extent carries useful biological and geometric information for matching adjacent sections of the same cardiomyocyte, whereas area-weighted OT marginals contribute mainly as a weak neighborhood-level prior that down-weights very small fragmented or truncated objects rather than serving as a second dominant size-based matching criterion.

With candidate generation, gold-standard definition, and the quantile-normalized scoring framework held unchanged (n=851), we compared top-1 hit rates against the gold-standard target under four combinations

In the present implementation, the transport coupling $T_\varepsilon^*$ is obtained by solving an entropy-regularized OT problem, but the reported neighborhood quantity is the primal transport cost evaluated at that regularized solution, $\langle T_\varepsilon^*, K\rangle$, where $K$ denotes the pairwise Soft-DTW cost matrix. This quantity is therefore not identical to the full entropy-regularized OT objective and should not be interpreted as the classical unregularized OT/EMD distance. Instead, $\langle T_\varepsilon^*, K\rangle$ is used as a regularized OT-derived neighborhood matching score for local candidate ranking. We report the primal transport term because it reflects the geometric discrepancy encoded in $K$, whereas the entropy term is introduced primarily for numerical smoothing and computational stability. Additional stability analyses showed that the induced rankings were highly robust across the regularization range used in this study and that the reported score retained substantial agreement with exact unregularized Earth Mover’s Distance on neighborhood pairs of tractable size.

#### 4. Dual-Scale Normalization for Stitching

During stitching, we jointly use DTW-, OT-, and centroid-based Euclidean distances for correspondence estimation, while explicitly handling their scale mismatch. Specifically, we used local distance-scale normalization for candidate ranking and acceptance decisions, and global distance-scale normalization for trajectory-level and cross-region quality evaluation. Local normalization was used only for matching, whereas global normalization was used only for reporting and comparing quality scores across trajectories, regions, and parameter settings.

For local matching, the normalization factor is estimated on-the-fly from the raw DTW/OT values computed in the current matching step (per source cell and its candidate set) by using the 75th percentile of the valid samples as the scale, so that DTW, OT, and centroid distance become numerically comparable when forming the weighted matching cost for candidate ranking and acceptance decisions; for efficiency, OT is evaluated only for better-ranked candidates and approximated by a constant penalty for the rest, and the locally normalized values are additionally clipped to reduce the impact of outliers.

In parallel, for global quality description, we determine a fixed, globally comparable normalization factor before each stitching stage by randomly sampling pairs in the current region and computing their raw DTW/OT distances, again setting the scale by the 75th percentile of these samples; this factor is kept unchanged within the stage and is used to map raw DTW/OT measurements into globally consistent scores that are recorded for subsequent trajectory-level and overall quality assessment and for cross-region/parameter comparisons.

#### 5. Match quality estimation based on margin

After obtaining the DTW-based contour dissimilarity, OT-based neighborhood dissimilarity, and centroid displacement, we make matching decisions for each candidate linkage under a relative local-margin framework (hereafter referred to as the local margin), using the same three distance components defined in the previous sections. For each current cell, the candidate linkages are first ranked within its own candidate set, and the separation between the best and second-best candidates is then used as a measure of decision confidence. Based on this unified margin statistic, we define both linkage-level trajectory quality and dataset-level global scores.

##### 5.1 Local adaptive normalization within the candidate set

For a current cell $c_i^z$ and its candidate set $\{c_j^{z+1}\}_{j=1}^{J}$ within a spatial window, we compute per-candidate distances:

$$\{D_{\mathrm{DTW}}^{(j)},\ D_{\mathrm{OT}}^{(j)},\ D_{\mathrm{cen}}^{(j)}\}_{j=1}^{J}\ .$$

Here, the candidate set is the set of cells on slice $z\ +\ 1$ that fall inside the pre-defined spatial search window of $c_i^z$. The three distances respectively quantify contour-shape discrepancy, neighborhood-structure discrepancy, and centroid displacement for each candidate linkage $(c_i^z, c_j^{z+1})$ .

We form a locally normalized loss for candidate ranking:

$$L_j^{(\mathrm{loc})} = w_{\mathrm{DTW}}\, \widehat{D_{\mathrm{DTW},J}^{(\mathrm{loc})}} + w_{\mathrm{OT}}\, \widehat{D_{\mathrm{OT},J}^{(\mathrm{loc})}} + w_{\mathrm{cen}}\, \widehat{D_{\mathrm{cen},J}^{(\mathrm{loc})}}\ ,$$

where $\widehat{D_{\mathrm{DTW},J}^{(\mathrm{loc})}}$ , $\widehat{D_{\mathrm{OT},J}^{(\mathrm{loc})}}$ and $\widehat{D_{\mathrm{cen},J}^{(\mathrm{loc})}}$ are the three distance terms normalized within the current candidate set of $c_i^z$. The weights $w_{\mathrm{DTW}}\,, w_{\mathrm{OT}}, w_{\mathrm{cen}}$ are fixed nonnegative coefficients shared across all candidate evaluations. and were optimized subsequently by Bayesian optimization. In parallel, we keep a globally normalized loss (using the same raw distances but a fixed, pre-estimated set of dataset-level normalization constants) for comparable quality reporting:

$$L_j^{(\mathrm{glob})} = w_{\mathrm{DTW}}\, \widehat{D_{\mathrm{DTW},J}^{(\mathrm{glob})}} + w_{\mathrm{OT}}\, \widehat{D_{\mathrm{OT},J}^{(\mathrm{glob})}} + w_{\mathrm{cen}}\, \widehat{D_{\mathrm{cen},J}^{(\mathrm{glob})}}\ .$$

Here, the globally normalized terms are obtained using a fixed set of dataset-level normalization constants estimated in advance, so that the resulting values remain comparable across different decision points, trajectories, and blocks. In our framework, the local loss is used for within-set candidate selection, whereas the global loss is retained for subsequent confidence reporting and trajectory-level aggregation. The local margin was used for acceptance because it reflects within-candidate-set separability, whereas the global margin was recorded for cross-linkage and trajectory-level comparison because it is computed on a fixed normalization scale.

##### 5.2 Local-margin matching criterion

We map losses to scores via a monotone decreasing transform:

$$S_j^{(\mathrm{loc})} = \frac{1}{1 + L_j^{(\mathrm{loc})}}, \quad S_j^{(\mathrm{glob})} = \frac{1}{1 + L_j^{(\mathrm{glob})}}.$$

Candidates are ranked by $S_j^{(\mathrm{loc})}$. Let $S_1^{(\mathrm{loc})} \geq S_2^{(\mathrm{loc})}$ denote the top-1 and top-2 scores. We define:

$$\Delta S^{(\mathrm{loc})} = S_1^{(\mathrm{loc})} - S_2^{(\mathrm{loc})}, \quad \mathrm{LocalMargin} = \frac{\Delta S^{(\mathrm{loc})}}{S_1^{(\mathrm{loc})}}.$$

That is, $\mathrm{LocalMargin}$ measures the local relative advantage of the best candidate over the runner-up within the current candidate set. A larger value indicates a more separated and hence more reliable local decision, whereas a small value indicates ambiguity between the top two candidates. We accept the top-1 match if $\mathrm{LocalMargin} \geq \tau_{\mathrm{rm}}$; otherwise the linkage is rejected (treated as unreliable for trajectory extension). Here, $\tau_{\mathrm{rm}}$ denotes a fixed local-margin threshold. For each accepted linkage, we assign a linkage index $k$ and additionally record its $\mathrm{GlobalMargin}$ based on the globally normalized scores. The globally normalized scores were computed using fixed DTW and OT value ranges estimated from a set of pre-sampled candidate linkages, so that the resulting GlobalMargin was measured on a dataset-level scale rather than on the local candidate-set scale.

$$\Delta S^{(\mathrm{glob})} = S_1^{(\mathrm{glob})} - S_2^{(\mathrm{glob})}, \quad \mathrm{GlobalMargin} = \frac{\Delta S^{(\mathrm{glob})}}{S_1^{(\mathrm{glob})}}$$

##### 5.3 Trajectory-level quality evaluation and conflict resolution

For a trajectory $\mathcal{T}$ that has already been constructed, suppose there are $M$ accepted valid linkages between adjacent slices, and the global boundary margin of the $m$-th linkage is $\mathrm{GlobalMargin_m}$. Then the margin quality of this trajectory is defined as

$$Q_{\text{margin}}(\mathcal{T}) = \frac{1}{M}\sum_{m=1}^{M} \text{GlobalMargin}_{\text{m}} .$$

This definition only considers accepted matches and summarizes the overall consistency of linkages along a trajectory. A larger average margin means that, for locally selected linkages, the locally top-ranked candidate is separated from the local runner-up by a wider gap on the globally normalized score scale, suggesting that most accepted linkages are unambiguous and that the trajectory is therefore more reliable.

When performing conflict resolution and high-quality trajectory selection, we combine $Q_{\text{margin}}(\mathcal{T})$ with the trajectory length $L(\mathcal{T})$ to construct a normalized composite quality score, so as to avoid favoring trajectories with very high confidence but very short length:

$$\widehat{Q_{\text{margin}}}(\mathcal{T}) = \frac{Q_{\text{margin}}(\mathcal{T}) - Q_{min}}{Q_{max} - Q_{min}}, \quad \hat{L}(\mathcal{T}) = \min\left(1, \frac{L(\mathcal{T})}{L_{\text{ref}}}\right),$$

where $Q_{max}$ and $Q_{min}$ are, respectively, the maximum and minimum margin quality among all trajectories in the current round, and $L_{\text{ref}}$ is the maximum trajectory length (this study is ten). The final score for each trajectory is then defined as

$$Q_{\text{track}}(\mathcal{T}) = \lambda_{\text{M}}\widehat{Q_{\text{margin}}}(\mathcal{T}) + \lambda_{\text{L}}\hat{L}(\mathcal{T}) .$$

In our experiments, we set $\lambda_{\text{M}}$ and $\lambda_{\text{L}}$ to 0.7 and 0.3, respectively. During conflict resolution for trajectory matching, when multiple candidate trajectories compete to occupy the same cell at a certain slice, we preferentially retain the trajectory with the higher $Q_{\text{track}}(\mathcal{T})$. This ensures that each cell participates in at most one trajectory, while favoring trajectories with higher composite quality.

**5.4 Global margin statistics and overall scoring**

The trajectory-level margin $Q_{\text{margin}}$ only considers accepted linkages; if we want to perform global margin statistics, then difficult-to-match or ambiguous samples, as well as segments where no trajectory is constructed, would be ignored. To more faithfully reflect the overall matching ambiguity, we record one margin statistic for every decision point in the procedure.

a) If there exists at least one candidate on the current slice, we record the corresponding

$\mathrm{GlobalMargin}$ and whether this match passes the threshold $\tau_{\mathrm{rm}}$ ;

b) If the current slice has no candidate at all, we record this decision point with $\mathrm{GlobalMargin}$ = 0 and mark it as “no candidate”.

In this way, over the entire dataset we obtain a sequence of global-level margins $\{\mathrm{GlobalMargin_t}\}_{t=1}^{T}$ , where $T$ denotes the total number of decision points evaluated during the whole procedure, and define the global average margin as:

$$\overline{m}_{\text{glob}} = \frac{1}{T}\sum_{t=1}^{T} \mathrm{GlobalMargin_t} .$$

$\overline{m}_{\text{glob}}$ can be interpreted as the average global confidence over the entire dataset. For convenient parameter tuning and comparisons across different datasets or blocks, we linearly rescale it to a global confidence score $\text{Conf}_{\text{glob}}$ in $[0,100]$. Meanwhile, we define the trajectory coverage rate as the proportion of cells covered by trajectories and map it directly to a coverage score CovScore in $[0,100]$. We also normalize the average trajectory length by a reference length (set to 10 layers) and map it to a length score LenScore in $[0,100]$. The final global score is obtained as a weighted sum of these three scores.

#### 6. Trajectory extension

We formulate the construction of cell trajectories as an iterative extension process based on the local-margin criterion defined above. Given an unassigned cell $c_i^z$, the algorithm takes this cell as the seed to initialize a candidate trajectory, then searches the adjacent slice for candidate successor cells and accepts the top-ranked candidate only when the aforementioned matching criterion is satisfied.

More specifically, the trajectory grows step-by-step along the $z$-axis, with the search restricted to the adjacent slice $z+1$ (and symmetrically, $\mathrm{z}-1$). In the target slice, we use the cell centroid coordinates in 3D space and restrict the search window to a fixed spatial radius, retaining only the cells whose spatial distance is sufficiently close as the candidate set $\{c_j^{z+1}\}_{j=1}^{J}$.

For the matching between the current cell and all its candidates, we compute the local confidence scores $S^{(\mathrm{loc})}$ as described previously, the candidate with the highest local score is selected as the tentative successor, while the second-best candidate is used only to compute the local margin. The linkage between the current cell and the top-1 candidate is accepted only if the local margin exceeds the threshold. In both the main dataset and the three consecutive-slice datasets constructed in this study, we retained only trajectories spanning at least three slices.

##### 7. Bayesian optimization–based parameter search framework

To obtain stable trajectory reconstruction performance across different tissue regions and imaging batches, we embed the entire outer layer (hyperparameter search) of the Margin-Based matching module into a Bayesian optimization framework, and automatically search the local-margin threshold and the three mode weights. The optimizer is based on gp_minimize (Gaussian-process–based Bayesian optimization, scikit-optimize) and uses multi-generation evaluation to maximize a comprehensive reconstruction-quality score defined on a given region.

###### 7.1 Parameter space and constraints

In the trajectory-construction and matching process described above, each single-step matching decision is jointly determined by the local-margin threshold $\tau_{\mathrm{rm}}$ and the linear combination weights ( $w_{\mathrm{DTW}}, w_{\mathrm{OT}}, w_{\mathrm{cen}}$ ) of the three distance modes (contour DTW distance, neighborhood OT distance, and centroid distance). We treat them as continuous hyperparameters to be optimized. Specifically, Bayesian optimization searches in the following 4-dimensional continuous space:

- Local-margin threshold $\tau_{\mathrm{rm}}$ , search range $[0,0.25]$, controlling the minimum confidence level of each step's linkage;
- Contour-similarity weight $w_{\mathrm{DTW}}$ , search range set to $[0.1,0.6]$;
- Neighborhood OT distance weight $w_{\mathrm{OT}}$ , search range set to $[0.1,0.6]$;
- Centroid-distance weight $w_{\mathrm{cen}}$ , search range set to $[0.1,0.6]$.

For each evaluation, we normalize the three weights to sum to 1 so that only their relative proportions affect the combined distance. The normalized weights are then used in the reconstruction pipeline to compute the local and global matching terms.

**7.2 Outer-loop objective for the reconstruction pipeline**

We treat each complete run of the 3D cell-trajectory reconstruction pipeline as a single evaluation of a given hyperparameter configuration $\boldsymbol{\theta} = (\tau_{\mathrm{rm}}, w_{\mathrm{DTW}}, w_{\mathrm{OT}}, w_{\mathrm{cen}})$.

For each hyperparameter configuration, we run the full 3D cell-trajectory reconstruction pipeline on the specified region and compute three predefined global evaluation metrics: $\mathrm{Conf}_{\mathrm{glob}}$ , CovScore, and LenScore. These three metrics are linearly combined into a scalar quality score:

$$\mathrm{GlobalScore}(\theta) = \gamma_C\,\mathrm{Conf}_{\mathrm{glob}} + \gamma_V\,\mathrm{CovScore} + \gamma_L\,\mathrm{LenScore}, \quad \gamma_C = 0.4,\ \gamma_V = 0.4,\ \gamma_L = 0.2.$$

The weights reflect our preference for overall confidence and coverage over trajectory-length completeness; trajectory length only serves as an auxiliary term, preventing the optimizer from trivially generating extremely short trajectories in order to obtain artificially high coverage. We have demonstrated the robustness of this weighting scheme in the paper.

**7.3 Gaussian-process surrogate and sampling strategy**

To achieve efficient exploration in the 4-dimensional search space under a limited evaluation budget, we adopt a Gaussian-process (GP)–based Bayesian optimization strategy. The implementation uses gp_minimize provided by scikit-optimize. Apart from the number of evaluations n_calls, the number of initial points n_initial_points, and the random seed, other options follow the library defaults.

Concretely, the surrogate model uses a zero-mean Gaussian-process regressor with a Matérn kernel; the prior over function values is the commonly used zero-mean GP prior, and the acquisition function is gp_hedge, which automatically switches between lower-confidence bound (LCB), expected improvement (EI), and probability of improvement (PI). The optimizer for maximizing the acquisition function is the default random-sampling–plus–L-BFGS strategy.

In the initialization stage, the optimizer first samples several starting points uniformly at random in the parameter space (8 points in this study). For each point, it performs a full run of trajectory reconstruction and records the corresponding

$$y_i = -\overline{\text{GlobalScore}}(\theta_i), \qquad i = 1, \ldots, n_{\text{init}}$$

treating $y_i$ as the observed value for the GP. The GP then fits these observations and builds a posterior regression model for the objective function, yielding the posterior mean $\mu(\boldsymbol{\theta})$ and variance $\sigma^2(\boldsymbol{\theta})$ at any candidate point. Because gp_minimize minimizes the objective, we pass $y_i$ to the optimizer; the selected configuration is the one with the largest original GlobalScore.

At each subsequent iteration, the gp_hedge acquisition function selects the next evaluation parameter set $\boldsymbol{\theta}^*$ in the EI/PI/LCB form on the basis of the current GP posterior and historical rewards. This process is repeated until the evaluation budget is exhausted.

##### 7.4 Determining the optimal parameters and final evaluation

After completing Bayesian optimization within the preset budget, we select from all historical evaluations the parameter set $\boldsymbol{\theta}^\star$ that achieves the highest overall score GlobalScore$(\boldsymbol{\theta})$, and treat it as the optimal configuration for that region.

#### 8. Linear-model-based feature ablation and fusion experiments

To systematically evaluate the contribution of different geometric and shape-similarity features to the cell-matching task, we conducted feature-combination and ablation experiments using the gold-standard dataset constructed from C57BL/6J mouse hearts. Five features were considered: DTW distance, OT distance, centroid distance, area difference, and aspect-ratio difference.

##### 8.1 Feature construction

The DTW distance, OT distance, centroid distance, area difference, and aspect-ratio difference for each candidate pair are computed exactly as described above, and for each pair of cells we finally obtain a 5-dimensional feature vector:

$$x = (\text{dtw\_loss}, \text{ot\_loss}, \text{centroid\_loss}, \text{area\_loss}, \text{aspect\_ratio\_loss})\,.$$

##### 8.2 Linear model and cross-validation protocol

In the feature-combination and ablation experiments, we use a linear logistic-regression model, implemented with LogisticRegression from scikit-learn. To avoid data leakage, all candidate pairs associated with the same query cell are assigned to the same fold; accordingly, we adopt 5-fold GroupKFold. In each fold, we train the logistic-regression model on the training folds; on the validation fold, we predict for every pair the probability of being a true match, and then aggregate the predictions from all folds to obtain the out-of-fold predicted score over the entire dataset.

##### 8.3 Evaluation methodology

For each query/source cell, the model outputs a matching score for all candidate target cells in its candidate set. We treat this as a ranking task of “retrieving the true matched cell” for each candidate, and define the following ranking-based evaluation metrics:

a) Recall@K: For each query/source cell, the model outputs scores for all cells in its candidate set. The candidates are ranked by predicted score, and Recall@K is counted as a hit if the true matched target cell appears within the top K candidates for that query cell.

b) Mean Rank / Median Rank: For every positive sample, we record its rank position (starting from 1) in the candidate ranking list, and then compute the mean and median ranks over all candidates to quantify the overall ranking quality.

##### 8.4 Feature-combination and fusion design

We exhaustively enumerate all non-empty subsets of the five basic features - DTW, OT, centroid distance, area difference, and aspect-ratio difference, and independently train and evaluate a linear model for each subset. For every feature combination, we record the following statistics: Recall@1, Recall@3, Recall@5, Mean Rank, Median Rank, and the number of Top-1 prediction hits and errors. We then rank all combinations by Recall@1 and report the top-10 combinations.

In addition, focusing on the core combination DTW + OT + Centroid, we design a more fine-grained fusion analysis by comparing: (1) the full three-feature model versus each of its two-feature subsets (dropping DTW, dropping OT, or dropping Centroid) in terms of Recall@K,

Mean Rank, and relative performance drop; and (2) the three single-feature models versus the three-feature combination, to examine their performance differences.

#### 9. Baseline-method design and robustness analysis

To systematically evaluate the effectiveness of HiDTW, we constructed three simple yet representative baseline methods and performed a systematic comparison using gold-standard pairs. Furthermore, to examine robustness to inter-slice systematic shifts and mild morphological perturbations, we designed multiple simulated perturbation scenarios and conducted comprehensive tests.

##### 9.1 Local Centroid Nearest Neighbor (Baseline 1)

Baseline 1 relies only on the 2D spatial locations of cell centroids and performs cross-section matching using a local nearest-neighbor rule, without considering the morphology or regional structural information of cells. For each golden-standard matched record, at the initial layer we locate the cell $c_a$ whose centroid is $(x_a, y_a)$. Then, when selecting candidate points within the local spatial window at the target layer, for each candidate $c$ we compute the Euclidean distance between the centroids and the candidate cell:

$$d(c_a, c) = \sqrt{(x_c - x_a)^2 + (y_c - y_a)^2}.$$

The candidate cell with the smallest distance is chosen as the predicted match for the query cell, and the predicted match is then compared with the true match.

##### 9.2 Global Hungarian Matching (Baseline 2)

Baseline 2 performs global one-to-one matching between each pair of adjacent layers using minimum-weight bipartite matching (the Hungarian algorithm). Unlike Baseline 1, which selects matches locally and independently, Baseline 2 jointly determines correspondences at the global level by minimizing the total matching cost.

For a given pair of adjacent layers, let $\mathbb{A}$ denote the set of query cells in the source layer and $\mathbb{B}$ denote the set of candidate cells in the target layer. We construct a cost matrix $C \in R^{|\mathbb{A}| \times |\mathbb{B}|}$, where each entry $C_{ij}$ represents the matching cost between cell $a_i$ and candidate cell $b_j$.

We restrict candidates using a local spatial window of size $W$. If $b_j$ lies outside the window of $a_i$, we set $C_{ij} = 10^6$ to effectively forbid this match. Otherwise, we define a normalized centroid distance and a relative area difference:

$$d_{ij}^{\text{centroid}} = \frac{\sqrt{(x_j - x_i)^2 + (y_j - y_i)^2}}{W}, \quad r_{ij}^{area} = \frac{|A_i - A_j|}{\max(A_i, A_j)}, \; max(A_i, A_j) > 0 \,.$$

The final cost is the equal-weight average of the two:

$$C_{ij} = 0.5\, d_{ij}^{\text{centroid}} + 0.5\, r_{ij}^{\text{area}}.$$

We then apply the Hungarian algorithm to $C$ to obtain the minimum-total-cost one-to-one assignment between $A$ and $B$. For each $a_i$, the assigned candidate is taken as its predicted match, and we compute Top-1 accuracy by comparing the prediction with the gold standard.

##### 9.3 Regional Structural Matching (Baseline 3)

Baseline 3 augments the centroid-distance measure in Baseline 1 with local neighborhood structure, including (i) the number of neighbors and (ii) the angular distribution of neighbor directions. This additional contextual information is expected to be more stable across adjacent slices than raw appearance, and helps disambiguate candidates that are close in Euclidean space but belong to different local structures.

**Neighborhood definition**. For each cell $c$, we define its neighbor set $\mathcal{N}(c)$ as all cells whose centroid-to-centroid distance to $c$ is within a fixed radius $R = 100$ pixels.

**Directional histogram**. For each neighbor $\mathrm{n} \in \mathcal{N}(c)$, let $(\Delta x, \Delta y)$ be the relative displacement from the centroid $c$ to the centroid of $n$. We compute the polar angle $\theta = \mathrm{atan2}(\Delta y, \Delta x)$ and map it to $[0, 2\pi)$. We then divide $[0, 2\pi)$ into 8 equal angular bins and count how many neighbors fall into each bin, forming an 8-dimensional histogram $h_c$ for cell $c$. If $|\mathcal{N}(c)| > 0$, we normalize $h_c$ by the neighbor count so that the histogram entries sum to 1; otherwise we set $h_c$ = 0.

**Matching cost.** For a query cell $c_a$ in the source layer and a candidate cell $c$ in the target layer, we define three cost terms. The first term is the centroid distance $d(c_a, c)$ as described previously. The second term is the difference in neighbor count; let $n_a = |\mathcal{N}(c_a)|$, $n_c = |\mathcal{N}(c)|$, respectively, and define:

$$\Delta n = |n_a - n_c|, \quad \Delta n_{norm} = \frac{|n_a - n_c|}{\max(1, n_a, n_c)}.$$

The third term is the L2 distance between directional histograms:

$$\Delta h = ||h_a - h_c||_2 .$$

The final matching cost is defined as the weighted sum of the three parts:

$$\text{score}(c_a, c) = 0.5 \cdot d(c_a, c) + 0.25 \cdot \Delta n_{\text{norm}} + 0.25 \cdot \Delta h .$$

For each golden-standard pair, we select, from all candidate points, the cell with the smallest score as the predicted match, and use this to compute the Top-1 accuracy of Baseline 3.

**9.4 Robustness analysis under simulated slice shifts and shape variations**

To evaluate the robustness of each method to possible systematic biases between slices, we apply controllable simulated perturbations to the original cell profile information, and under multiple noise levels we repeatedly compute the matching accuracy, thereby quantifying the impact of translation, rotation, and mild deformation perturbations on matching performance. In all simulation experiments, we only perturb the cell profiles in even-indexed layers while keeping the number of layers unchanged, so as to mimic the unilateral systematic errors that may occur when some slices are acquired or registered.

The **translation** transformation applies a uniform shift to the centroids of all cells in a given layer, thereby realizing rigid in-plane translation of that layer. Specifically, all cells are shifted along the positive x- and positive y-directions by the same amount:

$$(\Delta x, \Delta y) = (n, n),\ n \in \{0,5,10,15,20,25,30\} \text{ pixels.}$$

Rigid rotation (**Rotation**) takes the mean of all cell centroids in the layer, $c_{\text{cne}}$, as the rotation center, and applies an in-plane rigid rotation with angle θ ∈ 0,0.5,1.0,1.5,2.0,3.0. For each cell's centroid or contour point $p$, we perform (where $R_\theta$ is the standard 2D rotation matrix, preserving relative shape and local structure):

$$\tilde{p} = R_\theta(p - c_{\text{cne}}) + c_{\text{cne}}.$$

Mild affine deformation (**Affine**) uses the mean centroid within each layer as the deformation center, and applies an anisotropic scaling plus shearing linear transformation only to the even-indexed layers:

$$A = \begin{pmatrix} 1+\varepsilon_x & s \\ s & 1+\varepsilon_y \end{pmatrix}, \quad \varepsilon_x = \frac{m}{100}, \varepsilon_y = -0.5\,\varepsilon_x,\ s = 0.3\,\varepsilon_x.$$

where the deformation magnitude $\mathrm{m} \in 0,0.5,1.0,1.5,2.0,3.0\%$. The centroids and contours are first translated to the deformation-center coordinate system, then $A$ is applied, and finally they are translated back to the original coordinate system.

##### 9.5 Multi-scenario statistics under random combined perturbations

To avoid evaluation results relying excessively on a small number of manually set scenarios, we further generate $N = 30$ random combinations of perturbations. For each scenario, the translation, rotation, and affine parameters are independently sampled from the following uniform intervals:

$$\mathrm{n} \sim \mathcal{U}(0,30)\ \text{pixel}, \quad \theta \sim \mathcal{U}(0,3^\circ), \quad m \sim \mathcal{U}(0,3\%)\ .$$

For each random scenario, we run HiDTW and the three baselines on the perturbed cells, and obtain the Top-1 accuracy under that scenario. Aggregating the results from all 30 scenarios, we compute and report the accuracy distribution statistics of each method over the random scenarios.

#### 10. Pair- and Track-level Accuracy Metrics

We computed two accuracy metrics—**pair accuracy** and **track accuracy**—by comparing gold-standard pairwise relations with the stitched trajectories produced by each experiment. Gold-standard relations were represented as directed cell pairs (Start_cell, End_cell). Each experimental result was represented as a set of stitched trajectories, where each trajectory is an ordered list of cell IDs.

**Pair accuracy** was defined as the fraction of gold-standard cell pairs that were recovered by the experimental output. A gold-standard pair $(u, v)$ was counted as recovered if there existed trajectory containing both $u$ and $v$.

**Track accuracy** was defined as the fraction of gold-standard tracks that were fully matched by the experimental output. Gold-standard tracks were built from the gold-standard directed pairs by linking edges into paths starting from nodes with no predecessors. A gold-standard track $g = \{c_1, \dots, c_m\}$, was counted as matched if it appeared as a contiguous subsequence (with identical order) within any experimental trajectory.

**11. Simulation data and controlled benchmarking design**

To evaluate inter-slice cell matching under known ground truth, we generated parameterized three-dimensional cardiomyocyte phantoms within a fixed 500×500×100μm volume and virtually sectioned them into ten consecutive two-dimensional slices of 10μm thickness. Each simulated cell was represented as a capsule-like three-dimensional phantom whose cross-sectional contour was randomly sampled from a library of real cardiomyocyte contours, with configurable geometric properties, and each ground-truth trajectory comprised the observations of the same cell across adjacent sections.

The simulation included multiple controlled conditions with different spatial densities and perturbations, including Poisson-like or clustered cell placement, slice-wise rigid translation, per-cell lateral jitter across sections, and slice-specific distractor contours without cross-slice identity. We compared three single-cue baselines based on centroid proximity, contour similarity, or neighborhood optimal transport with the full HiDTW pipeline used in practice, with HiDTW parameters optimized by 20 rounds of Bayesian optimization. Performance was evaluated against the fully known ground truth using adjacent-section cell pairs and trajectory-level correspondence, reporting precision, recall, F1 score, identity-switch rate, and trajectory completeness and fragmentation after maximum-overlap assignment between predicted and true tracks.

**12. Benchmarking against alternative methods**

To compare HiDTW with published methods that could be adapted for cross-slice alignment, we benchmarked it against five representative nucleus-independent approaches based on spatial coordinates and gene expression, including PASTE2, SLAT, CAST, Spateo, and 3d-OT.

For the alternative methods, inter-slice correspondences were inferred from spatial and transcriptomic information; for HiDTW, the same cells were represented by contour geometry. To ensure comparability, all resulting adjacent-slice correspondences were normalized to one-to-one matches and linked in slice order using the same downstream trajectory-construction and conflict-resolution procedure.

Performance was assessed using label-free metrics computed from adjacent links along predicted trajectories, including centroid displacement, contour Dice coefficient, trajectory smoothness, trajectory length, cell coverage, and expression cosine similarity. These metrics were designed to quantify geometric consistency, continuity, and biological plausibility of the reconstructed cross-slice trajectories under a unified evaluation framework.

#### 13. 3D model construction

To build a 3D cell-trajectory model that carries layer index information, we construct the 3D model based on 2D cell contours and their matched cell trajectories, following the steps below.

##### 13.1 Cell-trajectory construction and identity annotation

From the trajectory summary table, we read each trajectory's track_id and its sequence of cell IDs. For each trajectory, we create a Track object, append the corresponding cell contours, and record its start- and end-slice indices. If the table provides additional tags (e.g., a Module column), we attach them to the Track for downstream colored visualization and statistics.

##### 13.2 Contour resampling and shape standardization

To support smooth 3D reconstruction, we resample each cell's 2D contour to a fixed number of points (NUM_POINTS = 100) using arc-length–based uniform resampling. Contours with fewer than three points are treated as degenerate and replaced by a regular polygon centered at the cell centroid, with radius estimated from the available points.

For valid contours, we enforce closure by setting the last point equal to the first (without adding extra points). We then compute cumulative arc length and sample evenly spaced positions along the total length; each new point is obtained by linear interpolation between neighboring original points. If needed, we truncate the final one or two samples (rather than padding) and finally force the last point to coincide with the first to keep the contour closed. This procedure yields consistent point counts, ordering, and orientation across slices, enabling one-to-one triangular mesh connections between adjacent slices.

##### 13.3 Z-direction scaling and inter-slice interpolation

To turn the stack of separated slices into a continuous 3D trajectory, we assign each slice a physical spacing along the Z-axis, and at the same time use a scaling factor to adjust the overall aspect ratio. In this experiment we set Z_DISTANCE = 5 and Z_SCALE = 8. Therefore, for a cell whose slice index is slice_idx, its corresponding Z coordinate is

$$Z = \text{slice\_idx} \times Z\text{\_DISTANCE} \times Z\text{\_SCALE}$$

For each trajectory, we sort cells by slice index and, for every pair of neighboring slices, perform inter-slice interpolation of the contours to generate a series of intermediate layers so that the cell shape varies smoothly along the Z-axis. Specifically, between every two neighboring slices we insert smooth_steps interpolation layers (in this experiment smooth_steps = 3), so that there are in total 5 cross-sections between them (including the two ends). For each interpolation layer we adopt a cubic smoothing function:

$$\text{smooth\_t} = t^2(3 - 2t), \quad t \in [0,1]$$

Here, $t \in [0,1]$ denotes the normalized interpolation position between two neighboring slices ($t = 0$ at the current slice and $t = 1$ at the next slice). We perform linear interpolation between the contour points of the current slice and the next slice according to this interpolation value, thereby generating smoothly transitioning contour points.

To further enhance the gradual shrinking or smoothing of cell shape on the intermediate layers, for the middle interpolation range $0.2 < t < 0.8$ we apply a slight centroid-contraction offset to the interpolated contours so that the transition looks more natural in 3D. This offset varies symmetrically with $t$, reaching its maximum in the middle of the trajectory and being minimal near the two ends.

##### 13.4 Triangular mesh construction and end-cap closing

After completing contour resampling and inter-slice interpolation, we stitch all contour points from all slices into a 3D vertex set and, by regular triangulation, generate a closed 3D mesh for each trajectory. The detailed procedure is as follows:

For each trajectory, we traverse adjacent slice pairs and, for every pair of corresponding indices $k$ and $k+1$, form a quadrilateral and then split it into two triangles:

- Triangle 1: ($prev\_layer_k$, $cur\_layer_k$, $prev\_layer_{k+1}$);
- Triangle 2: ($cur\_layer_k$, $cur\_layer_{k+1}$, $prev\_layer_{k+1}$)

To ensure that the mesh is closed, we additionally add end caps: we take the centroid of the contour on the start slice and end slice as the end-cap centers, respectively, add a new vertex at each center in the Z plane, and then connect this center with the contour vertices in sequence to form a fan of triangles, thereby sealing the opening of the cell at that end.

##### 13.5 Mesh smoothing and global merging

To reduce the faceted appearance at multi-slice connections, we apply Laplacian smoothing to the initial triangular mesh generated for each trajectory (using the smoothed interface in trimesh, with 3 iterations). After independently building and smoothing the mesh for every trajectory, we then use mesh-merging tools to combine the multiple trimesh objects corresponding to a single trajectory into an integral mesh, thus obtaining a complete dataset-level 3D model.

#### 14. 3D-Virtual CM expression data filling

To estimate gene expression at the single-cell level from spatial transcriptomics data, we mapped spatial barcodes to corresponding cell contours using segmentation masks derived from tissue images.

For each cell contour, we aggregated the expression values of all barcodes assigned to that contour. Gene expression for each contour was computed by grouping assigned barcodes by gene and summing their expression values within the contour. This procedure generated a contour-by-gene expression matrix, where each entry represents the total barcode expression of a given gene in a given contour. All steps, including image registration, barcode assignment, and expression aggregation, were implemented using custom scripts available in the accompanying code repository.

**15. 3D-Virtual CM Data Pre-processing**

For spatial transcriptomic reconstruction, each tissue section was subdivided into a 10 × 10 grid, and stitching was performed separately within each subregion to improve computational efficiency and preserve local transcriptomic structure. For each subregion, raw gene expression count matrices were imported into R (v4.1.3) and converted into Seurat objects (Seurat v4.3.0). Quality control (QC) metrics were computed for each spot/cell, including the number of detected genes (nFeature_RNA), total UMI counts (nCount_RNA), and the percentage of mitochondrial transcripts (percent.mt). Cells from all subregions were then merged into a single Seurat object and processed as follows: data were normalized using the LogNormalize method with a scale factor of 10,000; 2,000 highly variable genes were identified using the “vst” method; data were scaled and principal component analysis (PCA) was performed (20 PCs). A shared nearest neighbor graph was constructed using the first 10 PCs (FindNeighbors, dims = 1:10), and graph-based clustering was performed (FindClusters, resolution = 1.0). UMAP was computed using the first 10 PCs (dims = 1:10) for visualization. All downstream analyses were based on this processed dataset.

Because fibroblasts were the dominant source of non-cardiomyocyte contamination in cardiomyocyte data, we performed a fibroblast-focused QC. We computed Seurat module scores (AddModuleScore) for a cardiomyocyte gene program (CM_Score, *Tnnt2, Tnni3, Myh6/7, Actc1, Ryr2, Pln, Ttn*) and a fibroblast-lineage gene program (FibLineage_Score, *Pdgfra, Pi16, Dpt, C7, Col5a1/2, Fbln1/2, Thy1*). To account for differences in overall cardiomyocyte program strength, we fit a linear model across cells: FibLineage_Score~CM_Score. We then defined a fibroblast-contamination risk score as the residual fibroblast-lineage score from this regression (i.e., FibLineage_Score after regressing out CM_Score). Cells in the upper tail of this residual risk score (top 10%; 90th percentile) were labeled high risk and excluded from downstream analyses.

#### 16. Differential expression and module-enriched genes

Differential expression analysis was performed in Seurat (v4.1.3) using the Wilcoxon rank-sum test. P-values were adjusted using the Benjamini–Hochberg procedure to control the false discovery rate (FDR). Unless otherwise stated, genes with adjusted $p<0.05$ were considered significant.

To identify module-enriched (highly expressed) genes, we ran FindAllMarkers and retained markers with adjusted $p<0.05$. In addition, to account for expression abundance, we computed for each module the fraction of UMI counts contributed by each marker gene (UMI percentage) using the raw UMI count matrix, and summarized gene-wise UMI percentages within the target module and in the remaining cells for comparison.

#### 17. Gene set enrichment analysis (GSEA) and GO enrichment

GSEA was performed using gseapy (v1.1.8) in preranked mode. For each module, genes were ranked by the Seurat-derived average log2 fold-change (avg_log2FC) from the corresponding differential expression results. Gene sets were obtained from MSigDB for mouse (release 2024.1.Mm), using the m5.go collection (GO Biological Process, Cellular Component, and Molecular Function). Preranked GSEA was run with 1,000 permutations. Enrichment statistics (ES, NES) were computed following the standard GSEA procedure. Multiple testing across gene sets was controlled using the Benjamini–Hochberg method; unless otherwise specified, gene sets with FDR<0.05 were considered significantly enriched. GO enrichment analysis was performed using the Enrichr web platform (https://maayanlab.cloud/Enrichr/)

#### 18. pySCENIC regulon inference and activity analysis

We inferred transcriptional regulons using pySCENIC (v0.12.1) in three steps: (i) GRNBoost2 co-expression network inference on the expression matrix using a mouse TF list (mm_mgi_tfs.txt); (ii) cisTarget motif enrichment and pruning using the mm9 TSS-centered 5 kb, 7-species mc9nr rankings and v10nr motif annotations to obtain curated regulons; and (iii) AUCell scoring to compute per-cell regulon activity (AUC). For downstream comparisons between the “CC-CMs” and “BZ-CMs” groups, we loaded AUC scores and metadata, binarized regulon activities, retained shared cells, and computed per-regulon group-wise mean AUC and activation rate. Between-group differences in AUC were assessed using two-sided Mann–Whitney U tests and adjusted across regulons using the Benjamini–Hochberg procedure (statsmodels).

##### 19. 3D-Virtual CM contour-expression analysis

Contour-based expression data for 10-layers 3D-VCMs from Module1 were imported into Seurat for downstream analysis. QC metrics (nFeature_RNA, nCount_RNA, and percent.mt) were computed and visualized using violin plots. For robust downstream computation and to emphasize conservative biological signals, we focused on cells with intermediate library sizes by retaining those with nCount_RNA between the 15th and 75th percentiles.

Slice identity was extracted from cell barcodes and stored as a metadata field. The retained dataset was re-normalized and scaled while regressing out total UMI counts (vars.to.regress = nCount_RNA), followed by PCA. Graph-based clustering was performed using the first 10 principal components (FindNeighbors, dims = 1:10; FindClusters, resolution = 0.6), and UMAP embeddings were computed for visualization (RunUMAP, dims = 1:10).
